# Intermediate Filaments Associate with Aggresome-like Structures in Proteostressed *C. elegans* Neurons and Influence Large Vesicle Extrusions as Exophers

**DOI:** 10.1101/2022.08.03.501714

**Authors:** Meghan Lee Arnold, Jason Cooper, Rebecca Androwski, Sohil Ardeshna, Ilija Melentijevic, Joelle Smart, Ryan J. Guasp, Ken C.Q. Nguyen, Ge Bai, David H. Hall, Barth D. Grant, Monica Driscoll

## Abstract

Under conditions of proteostasis disequilibrium, neurons can enhance intracellular and extracellular protective mechanisms to guard against neurotoxicity. In mammals, an intracellular response to severe proteostasis imbalance that results from proteosome inhibition is the formation of juxtanuclear intermediate filament-surrounded, aggregate-filled aggresomes, which sequester threatening aggregates for later disposal via lysosomal degradation. Highly proteo-stressed neurons can also engage the assistance of neighboring cells in aggregate removal by loading threatening materials into large exopher vesicles that are transferred to neighboring cells for remote degradation of contents, a process that has been suggested to be analogous to the process that enables aggregate spreading in the human brain in neurodegenerative disease. In *C. elegans* these large extruded vesicles are called exophers.

Here we document that players involved in aggresome biology are required for the elimination of potentially deleterious materials in neuronal exophers. We show that in proteostressed *C. elegans* touch receptor neurons, intermediate filament proteins IFD-1 and IFD-2 can assemble into juxtanuclear structures with multiple molecular and cellular characteristics of mammalian aggresomes. IFD-concentrating structures depend upon orthologs of mammalian adapter proteins, dynein motors, and microtubule integrity for aggregate collection into juxtanuclear compartments where they associate with ubiquitinated and neurotoxic polyglutamine expansion proteins. Strikingly, disruption of aggresome-decoration genes encoding IFDs or disruption of the BAG/14-3-3/Hsc70 adapter that promote aggregate loading of aggresome-like organelles, lowers exopher production via a cell autonomous mechanism. Although aggresome-like structures are not mandatory exopher cargo, IFD compartments can be extruded from neurons in exophers, revealing a previously unreported strategy to eliminate neuronal aggresome-like organelles via transfer to neighboring cells. Human IF neurofilament light chain hNFL can partially substitute for *C. elegans* IFD-2 proteins in promoting exopher production, indicating conservation of the capacity of intermediate filaments to influence neuronal aggregate extrusions across phyla. In sum, we identify a requirement for specific intermediate filaments, counterparts of human biomarkers of neuronal injury and disease and major components of Parkinson’s disease Lewy bodies, in *C. elegans* neuronal aggresome-like organelle formation and large vesicle exopher extrusion from stressed neurons.

## Introduction

Disrupted proteostasis underlies cellular dysfunction in aging and neurodegenerative disease,^1^ and thus elaborating the cellular mechanisms of protein quality control is a major focus in biomedical research. The cellular commitment to protein quality control engages multiple molecular strategies, including protein refolding via the chaperone network, protein degradation via the ubiquitin proteasome system, and aggregate elimination via autophagy,^2^ all of which work to cull misfolded proteins that otherwise might impair healthy macromolecular functions.^3^ Under extreme conditions of proteostress, additional neuroprotective strategies can be engaged. For example, when the proteasome is pharmacologically inactivated, ubiquitinylated proteins are identified by molecular adaptors that link aggregates/misfolded proteins to dynein motors, which in turn transport their cargo along microtubule tracks to deliver the aggregates to a juxtanuclear domain called the aggresome.^4–6^ The aggresome concentrates and sequesters aggregates within an intermediate filament-surrounded domain that can ultimately deliver aggresome contents to lysosomes for degradation. Aggregate sequestration by aggresomes is generally considered beneficial as function-disrupting aggregates are prevented from interfering with cytoplasmic activities.

There are additional protective strategies that neurons can engage under extreme conditions of proteostasis disruption. In *C. elegans,* high proteostress that likely overwhelms chaperone, proteasome, and autophagy functions can trigger the production of large (∼3.8 um) vesicles called exophers that concentrate, and somewhat selectively remove, aggregating proteins such as transgenically expressed Huntingtin polyQ expansion proteins.^7–9^ Extruded neuron-derived exopher vesicles enter the surrounding glial-like hypodermal cell via specialized phagocytosis and exopher contents are then degraded by the hypodermal cell’s lysosomal network,^10^ transfer biology that may be conserved in flies^8, 11, 12^ and mammalian neurons.^13^ Proteostressed *C. elegans* neurons that produce exophers maintain functionality in late-life better than similarly proteostressed neurons that do not make exophers, suggesting that exophergenesis is neuroprotective.^7^ Exopher like biology has been reported in mammalian models^14–19^ and thus the discovery of neuronal extrusion in the transparent genetic model *C. elegans* enables dissection of exophergenesis in an *in vivo* context that may well inform on the fundamental biology of aggregate spreading that occurs in human neurodegenerative disease pathology.^13^

Here we report on aggregate accumulation and extrusion in proteostressed *C. elegans* touch receptor neurons. We document roles for conserved adapter proteins, dynein motors and microtubule integrity in collection and concentration of aggregating/highly expressed proteins into intermediate-filament-associated compartments that share multiple molecular features of mammalian aggresomes. We identify a cell-autonomous requirement for aggresome-associated intermediate filament (IF) D class proteins IFD-1 and IFD-2, as well as components of the 14-3-3/BAG3/Hsc70 adaptor complex that can deliver cargo to aggresome-like organelles, in efficient *C. elegans* exophergenesis. Human neuronal intermediate filament hNFL can partially execute touch neuron-autonomous exopher-related functions in *C. elegans* neurons, supporting that IF roles in aggregate transfer during proteostress may be conserved across phyla. Our data contribute molecular documentation of *C. elegans* neuronal aggresome-like organelle formation and provide the first demonstration of aggresome-like component contributions to stress-associated extrusion of compromised neuronal contents. Mechanistic linking of conserved aggresome biology with neuronal aggregate extrusion suggests novel alternative aggregate clearance options for mammalian aggresomes, and invites consideration of a new angle for therapeutic design in neurodegenerative disease.

## Results

Cellular proteostasis is maintained by balancing protein synthesis and protein degradation, with dedicated removal of misfolded and aggregated proteins via ubiquitin proteosome mediated degradation and autophagy.^2^ Under conditions of extreme proteostress, mammalian cells sequester potentially harmful aggregates in a juxtanuclear aggresome characterized by a “cage” of collapsed intermediate filament proteins. Aggresome contents eventually can undergo lysosomal degradation via autophagy.^5^ Details on how aggresomes are eliminated from cells, however, remains a poorly understood facet of proteostasis biology.

*C. elegans* has served as a highly illuminating model for the elaboration of fundamental mechanisms of aging and proteostasis.^20^ Most literature indications of *C. elegans* aggresomes.^21, 22^ refer to the presence of fluorescently tagged protein aggregates near the nucleus, but little documentation on molecular make up and contents of aggresomes is available in this model, especially for neurons. With an interest in the fundamental biology of neuronal proteostasis, we first asked whether aggresomes form in proteo-stressed *C. elegans* neurons. We chose to focus on easily visualized *C. elegans* touch neurons expressing high levels of mCherry for which previous ultrastructural examination and cell biological characterization provided evidence of high stress and neuronal exopher extrusions.^7^ The mCherry strain houses integrated transgene allele *bzIs166*[P*_mec-4_*mCherry], which produces high levels of touch neuron-specific mCherry that permits facile visualization of touch neurons and the exophers they can produce.^7, 8, 23^ For simplicity, we refer to this reporter allele as mCherry in the text that follows.

### In proteostressed neurons, intermediate filament proteins IFD-1 and IFD-2 colocalize in juxtanuclear foci that are distinct from Golgi, lysosomes and mitochondria

To begin, we sought evidence of aggresome formation by characterizing intermediate filament protein distribution in proteostressed mCherry touch receptor neurons. *ifd-1* and *ifd-2* are documented to be transcriptionally expressed in touch neurons^24–26^ and thus we studied the subcellular localization patterns of functional fluorescently tagged GFP::IFD-1 and mNeonGreen::IFD-2 proteins expressed in touch neurons from single copy transgenes. We found that GFP::IFD-1 and mNeonGreen::IFD-2 typically appear in one to three ∼ 1 μm foci located near the nuclear membrane in the mCherry background (Figure 1A-B), with two IFD foci being the number most commonly observed during early adult life (Supplementary Figure 1A). The IFD foci in mCherry-stressed neurons increase in size from L4 to adulthood (Figure 1C, Supplementary Figure 1B). We also found that additional proteostress, such as overexpression of toxic proteins or genetic impairment of autophagy with the *epg-9(bp390)* mutant, results in a modest increase in the number and size of IFD foci (Supplementary Figure 1C-E). When animals are stressed with pharmacological proteasome and autophagy inhibitors, foci can increase in size (Supplementary Figure 1F). We found that juxtanuclear IFD foci can be visualized using multiple different fluorescent reporter-tagged versions of IFD-1 and IFD-2 (*i.e.,* for mNeonGreen, RFP, and mScarlet fluorescent tags; Supplementary Figure 1G-J), indicating that the subcellular localization of IFDs is independent of fluorescent tag identity. Overall, data support that intermediate filaments IFD-1 and IFD-2 concentrate into juxtanuclear foci, the dimensions of which can increase under proteostress.

**Figure 1:**
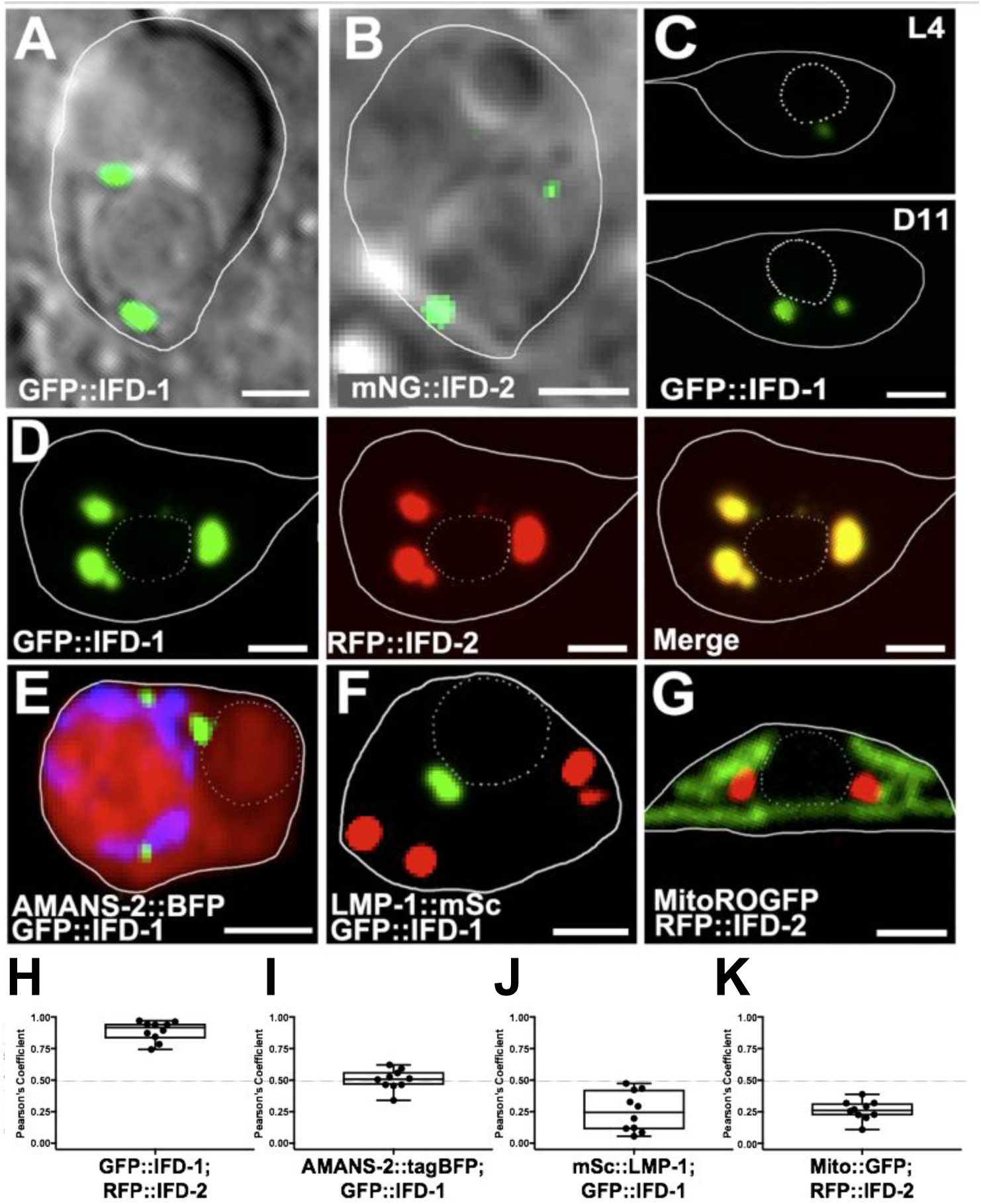
IFD-1 and IFD-2 colocalize to juxtanuclear inclusions that become larger with age and are distinct from other organelles. For images as indicated, touch cell soma outline is indicated by a continuous line and nucleus is marked by a dotted line, except in A, B where nuclear membrane is clear. **A) GFP::IFD-1 concentrates in juxtanuclear inclusions.** Shown is a representative Ad2 ALM touch neuron, DIC-GFP merge, strain expressing *bzIs166*[P*_mec-4_*mCherry]; *ifd-1*(*ok2404*)X; *bzSi3*[P*_mec-7_*GFP::IFD-1] (a rescuing single-copy GFP::IFD-1 transgene). Solid white lines outline the touch neuron soma. Image is representative of N > 40 for this strain. Scale bar = 2 µm. **B) mNeonGreen::IFD-2 concentrates in juxtanuclear inclusions.** Shown is a representative Ad2 ALM touch neuron, DIC-GFP merge, strain expressing *bzIs166*[P*_mec-4_*mCherry];*bzSi37*[P*_mec-7_*mNeonGreen::IFD-2]. Solid white lines outline the touch neuron soma. Image is representative of N > 20 with visible mNG expression for this strain. Scale bar = 2 µm. **C) IFD-1-positive inclusions become larger with age. Top:** L4 larval stage ALM neuron. **Bottom:** Ad11 ALM neuron. Strain expresses *bzIs166*[P*_mec-4_*mCherry]; *bzSi3*[P*_mec-7_*GFP::IFD-1]. Solid white lines outline the touch neuron soma. Images representative of > 100 observations. Scale bar = 2 µm. **D) IFD-1 and IFD-2 colocalize to juxtanuclear inclusions.** Representative of adult touch neuron, strain expressing *bzEx279*[P*_mec-7_*GFP::IFD-1 P*_mec-7_*RFP::IFD-2]. Image is representative of N > 20. Scale bar = 2 µm. Quantitation of colocalization in panel H. **E) GFP::IFD-1 does not extensively colocalize with a Golgi marker.** Representative of adult touch neuron, strain expressing *bzIs166*[P*_mec-4_*mCherry]; *bzEx265*[P*_mec-4_*TagBFP::AMANS-2];*bzSi3*[P*_mec-7_*GFP::IFD-1]. Image is representative of N > 20. Scale bar = 2 µm. Quantitation of colocalization in panel I. **F) GFP::IFD-1 does not extensively colocalize with the LMP-1::mSc lysosome marker.** Representative of adult touch neuron, strain expressing *bzIs3*[P*_mec-7_*GFP::IFD-1]; *pwSi222*[P*_mec-7_*LMP-1::mScarlet]. Representative of N > 20. Scale bar = 2 µm. Quantitation of colocalization in panel J. **G) IFD inclusion does not colocalize with mitoROGFP mitochondria marker.** Representative of adult touch neuron, strain expressing *bzEx253*[P*_mec-7_*RFP::IFD-2]; *zhsEx17*[P*_mec-4_*mitoLS::ROGFP]. Representative of N > 20. Scale bar = 2 µm. Quantitation of colocalization in panel K. Note: We did note that IFD inclusions were often found next to Golgi or lysosomes, and thus the possibility that the IFD compartments might interact with these organelles, potentially via transient associations, should be entertained in future study. **H)** Colocalization correlation of GFP::IFD-1 and RFP::IFD-2 signals (panel D) graphed as Pearson’s Coefficient of red and green channel. N = 10, error bars are SEM. **I)** Colocalization correlation of GFP::IFD-1 and TagBFP::AMANS-2 signals (panel E) graphed as Pearson’s Coefficient of blue and green channel. N = 10, error bars are SEM. **J)** Colocalization correlation of GFP::IFD-1 and mSc::LMP-1 signals (panel F) graphed as Pearson’s Coefficient of red and green channel. N = 10, error bars are SEM. **K)** Colocalization correlation of RFP::IFD-2 and mitoROGFP (panel G) graphed as Pearson’s Coefficient of red and green channel. N = 10, error bars are SEM.

To determine whether IFD-1 and IFD-2 colocalize with each other in the juxtanuclear foci, we tested for co-incidence in strains that expressed both GFP::IFD-1 and RFP::IFD-2 in touch neurons (Figure 1D). We find that GFP::IFD-1 and RFP::IFD-2 signals are nearly completely colocalized (quantitation in Figure 1H). To address whether the juxtanuclear IFD foci correspond to known cell organellar compartments, we tracked IFD-1 and IFD-2 localization relative to established Golgi, lysosome, and mitochondrial reporters. We found that tagged IFD proteins did not colocalize substantially with Golgi marker AMAN-2::tagBFP (Figure 1E, I), late endosome and lysosome marker LMP-1::mScarlet (Figure 1F, J), or mitochondria tagged with MitoROGFP (Figure 1G, K). Thus, IFDs localize to a neuronal compartment distinct from Golgi, lysosomes, and mitochondria.

In sum, in proteo-stressed touch neurons, tagged intermediate filaments IFD-1 and IFD-2 concentrate into juxtanuclear foci that have subcellular localization features akin to those described for mammalian aggresomes.

### Electron microscopy reveals intermediate filament-sized aggresome-like structures in mCherry stressed touch neurons

We examined ultrastructural features of ALM neuron somata from mCherry proteo-stressed touch receptor neurons for evidence of aggresome-like structures, focusing on twelve distinct ALM touch neurons in adult day 2 (Ad2) mCherry animals prepared using high pressure freezing/freeze substitution (HPF/FS) and visualized using serial section electron microscopy and in some cases 3D electron tomography.

We noted that the proteo-stressed ALM somata often contained 1-2 μm-sized rounded structures located close to neuronal nuclei that included a central granular matrix (Supplementary Figure 2A-D). A striking feature of the juxtanuclear structures was their peripheral-association with 10 nm-wide filaments (9.99 nm; > 150 filaments measured, +/-0.80 SD), a diameter measure consistent with that of Intermediate Filaments (IF). Electron tomograms confirm that these ∼ 10 nm filaments can be readily distinguished from membrane bilayers (Supplementary Videos 1, 2).

Data from the twelve neurons we examined suggest a potential “growth/maturation” sequence for the spherical structures. Some of these spherical granular structures contain a patched meshwork-like border of single discontinuous filaments at the periphery, and a few disorganized short loose intermediate filaments inside (Supplementary Figure 2A, B). Some granular structures that may be more advanced in genesis feature a series of longer curved filaments assembled in parallel at the periphery, with each filament remaining separate from its neighbors, and fewer short filaments inside the structure (Supplementary Figure 2C). Finally, some rounded juxtanuclear structures display a periphery of loosely-parallel filaments, with some portions of the border having a densely packed layer of filaments with no internal filament structures evident (Supplementary Figure 2D). We noted that the electron density of the central granular material appears higher in samples with greater filament association.

Although technical challenges precluded our immuno-EM verification of intermediate filament identity, the structures we documented in mCherry expressing ALM neurons are strikingly reminiscent of maturing mammalian aggresomes that form under high proteostress and are characterized by intermediate filament proteins that coalesce around sequestered aggregated protein,^4, 6, 27, 28^ consistent with our florescence microscopy visualization of tagged IFD proteins in proteo-stressed neurons.

### IFD foci depend on MTs and dynein motors, and associate with adapter proteins SQST-1, HSP-1 and FTT-2/14-3-3

Under high proteostress conditions, ubiquitinated and non-ubiquitinated aggregated proteins in mammalian (and other) cells can be linked to adapters such as HDA-6/HDAC6, SQST-1/p62, and the HSP-1/BAG3/14-3-3 complex, which connect to dynein motors and move along microtubule tracks to juxtanuclear sites to deposit aggregate cargos^29–31^ (see summary diagram in Figure 2A). IFs concentrate at these juxtanuclear aggresome-like organelle sites. To address whether formation of the *C. elegans* IF compartment features represents an aggresome-like organelle, we perturbed key aspects of aggresome biogenesis and probed for colocalization of IFDs and homologs of mammalian aggresome proteins.

**Figure 2:**
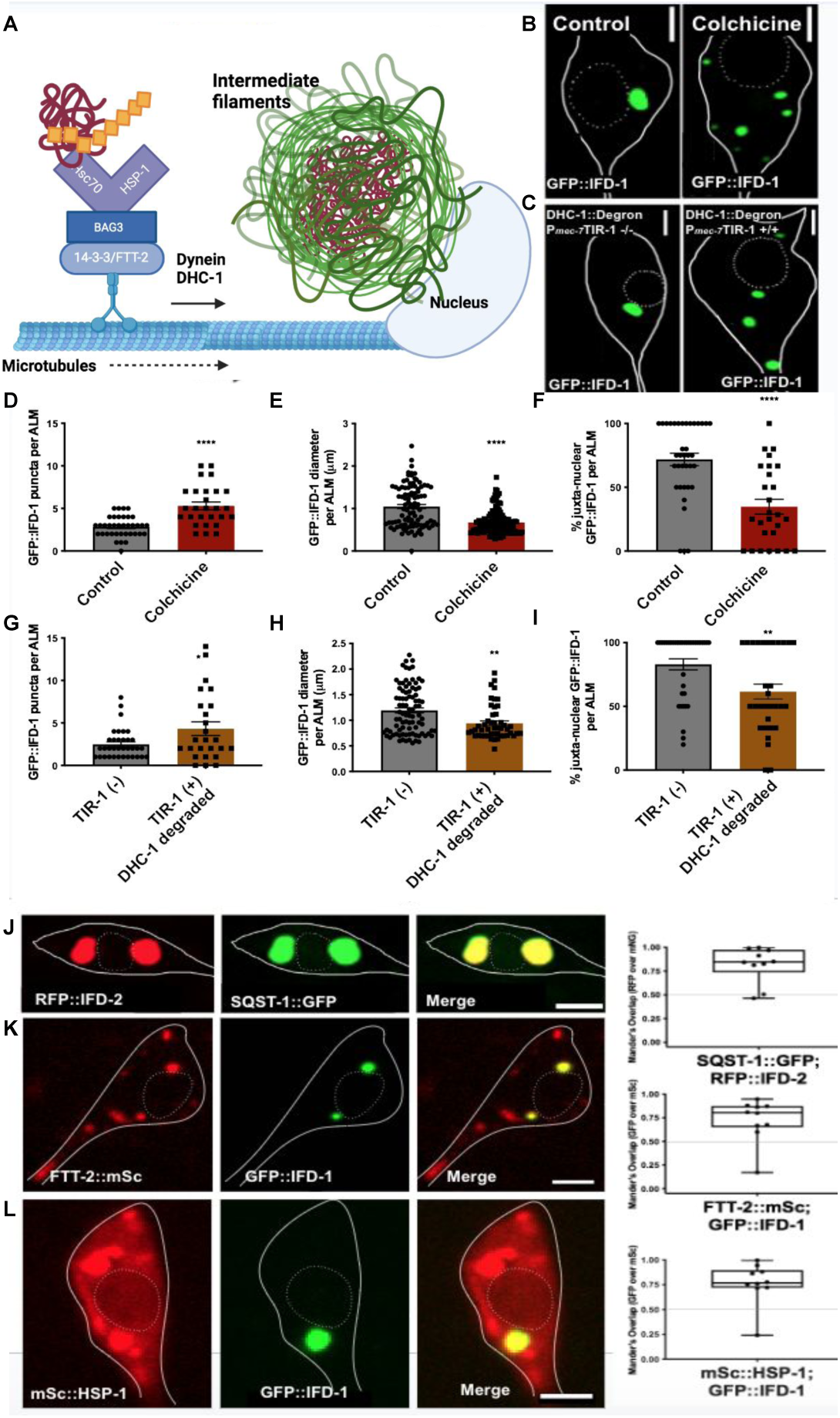
Genesis of juxtanuclear IFD inclusions depends on functional microtubules and dynein; aggregate adapter proteins 14-3-3/FTT-2 and HSP-1 colocalize with IFD-positive organelles. Solid white lines outline the soma cell body and white dashed lines outline the nucleus in panels where indicated. **A) Example of a mammalian aggregate-adapter complex.** Adapted from Xu 2013 ^36^, created with BioRender.com. 14-3-3 proteins are able to bind directly to dynein components. 14-3-3(FTT-2) binds BAG3-bound Hsc70/(HSP-1). Hsc70/HSP-1 recognizes and binds ubiquitinylated aggregates. The complex moves along the microtubule tracks to deliver the transported aggregate to the aggresome-like compartment. *C. elegans* counterpart names are indicated. **Microtubules and dynein are required for robust GFP::IFD-1 inclusion formation.** **B) Microtubule inhibition via colchicine treatment from L4 to Ad2 reduces IFD-1-positive inclusion formation.** Strain expressing *bzIs166*[P*_mec-4_*mCherry]; *bzSi3*[P*_mec-7_*GFP::IFD-1]**. Left:** Control, DMSO added from L4 to Ad2, Ad2 ALM. **Right:** 5 mM colchicine in DMSO added from L4 to Ad 2, Ad2 ALM. Scale bar = 2 µm. Image representative of N > 90. **C) Touch neuron-specific dynein knockdown via auxin inducible degradation of DHC-1::degron reduces IFD-positive puncta formation.** Knockdown strain expresses *bzIs166*[P*_mec-4_*mCherry]; *bzSi6*[P*_mec-7_*TIR-1::TagBFP]; *bzSi3*[P*_mec-7_*GFP::IFD-1]; *dhc-1*(*ie28*[DHC-1::degron::GFP]) and control strain lacks the TIR-1 component (*bzSi6*[P*_mec-7_*TIR-1::TagBFP]) and is therefore DHC-1 is not susceptible to degradation. **Left:** Control TIR-1 -/-, Ad2 ALM. **Right:** TIR-1 +/+, Ad2 ALM; 5 mM auxin exposure to the control and the experimental, from L4 to Ad2. Scale bar = 2 µm. Image representative of N > 90. **Quantification of GFP::IFD-1 inclusion formation upon microtubule and dynein knockdown.** **D) Microtubule inhibition via colchicine increases the number of IFD-1 puncta.** Number of IFD-1-positive puncta per Ad2 ALM in a strain expressing *bzIs166*[P*_mec-4_*mCherry]; *bzSi3*[P*_mec-7_*GFP::IFD-1]. Control DMSO, colchicine 5 mM. L4 to Ad2 exposure. ***P < 0.0005, N > 27/strain, 3 trials, two-tailed t-test, error bars are SEM. **E) Microtubule inhibition via colchicine treatment from L4 to Ad2 reduces ALM IFD-1 puncta size**. Diameter of IFD-1-positive organelle in ALM Ad2 in µm in a strain expressing *bzIs166*[P*_mec-4_*mCherry]; *bzSi3*[P*_mec-7_*GFP::IFD-1]. Control DMSO, colchicine 5 mM L4 to Ad2 exposure. ****P < 0.0005, N > 99/strain, 3 trials, two-tailed t-test, error bars are SEM. **F) Microtubule inhibition via colchicine treatment from L4 to Ad2 reduces the juxtanuclear location IFD-1 puncta.** Percentage of juxtanuclear foci in a strain expressing *bzIs166*[P*_mec-4_*mCherry]; *bzSi3*[P*_mec-7_*GFP::IFD-1]; control DMSO, colchicine 5 mM L4 to Ad2 exposure. ****P < 0.0005, N > 99 / strain, 3 trials, two-tailed t-test, error bars are SEM. **G) Touch neuron-specific dynein knockdown increases the number of GFP::IFD-1 puncta:** Number of IFD-1-positive puncta per Ad2 ALM. Strains are ZB4871 *bzIs166*[P*_mec-4_*mCherry]; *bzSi6*[P*_mec-7_*TIR-1::TagBFP]; *bzSi3*[P*_mec-7_*GFP::IFD-1]; *dhc-1*(*ie28*[DHC-1::degron::GFP]) and ZB4872 *bzIs166*[P*_mec-4_*mCherry]; *bzSi3*[P*_mec-7_*GFP::IFD-1]; *dhc-1*(*ie28*[DHC-1::degron::GFP]). Auxin exposure to both the control and the experimental from L4 to Ad2. *P < 0.05, N > 25 / strain, 3 trials, two-tailed t-test, error bars are SEM. **H) Cell-specific DHC-1 dynein knockdown via AID reduces ALM Ad2 IFD-1 puncta size.** Diameter of IFD-1-positive organelle in ALM Ad2 is measured in µm. Strains are ZB4871 *bzIs166*[P*_mec-4_*mCherry]; *bzSi6*[P*_mec-7_*TIR-1::TagBFP]; *bzSi3*[P*_mec-7_*GFP::IFD-1]; *dhc-1*(*ie28*[DHC-1::degron::GFP]) and ZB4872 *bzIs166*[P*_mec-4_*mCherry]; *bzSi3*[P*_mec-7_*GFP::IFD-1]; *dhc-1*(*ie28*[DHC-1::degron::GFP]). Control is -/- for P*_mec-7_*TIR-1, *dhc-1* knockdown strain is +/+ for P*_mec-7_*TIR-1. Both strains were exposed to 5 mM auxin from L4 to Ad2. **P < 0.005, N > 86 / strain, 3 trials, two-tailed t-test, error bars are SEM. **I) Cell-specific DHC-1 dynein knockdown via AID reduces the juxtanuclear location of Ad2 ALM IFD-1 puncta.** Percentage of juxtanuclear foci. Strains are ZB4871 *bzIs166*[P*_mec-4_*mCherry]; *bzSi6*[P*_mec-7_*TIR-1::TagBFP]; *bzSi3*[P*_mec-7_*GFP::IFD-1]; *dhc-1*(*ie28*[DHC-1::degron::GFP]) and ZB4872 *bzIs166*[P*_mec-4_*mCherry]; *bzSi3*[P*_mec-7_*GFP::IFD-1]; *dhc-1*(*ie28*[DHC-1::degron::GFP]). Control strains is -/-for P*_mec-7_*TIR-1, *dhc-1* knockdown strain is +/+ for P*_mec-7_*TIR-1. Both strains were exposed to 5 mM auxin from L4 to Ad2. **P < 0.005, N > 30 / strain, 3 trials, two-tailed t-test, error bars are SEM. **Homologs of aggresome adaptor proteins colocalize to IFD-1 puncta.** **J) SQST-1 colocalizes to IFD-1 puncta**. Representative of adult touch neuron expressing *bpIs151*[P*_sqst-1_*SQST-1::GFP + *unc-76*(+)]; *bzEx261*[P*_mec-7_*RFP::IFD-2]. Representative of N > 10. Scale bar = 2 µm. Quantitation of co-localization of N = 10, graphed on the right as Mander’s overlap coefficient of mScarlet channel over GFP channel. **K) FTT-2::mSc colocalizes with juxtanuclear GFP::IFD-1.** Representative of adult touch neuron expressing *bzSi51*[P*_mec-7_*FTT-2::mSc]; *bzIs3*[P*_mec-7_*GFP::IFD-1], N > 10. Scale bar = 2*µm*. Quantitation of co-localization of N = 10, graphed on the right as Mander’s overlap coefficient of GFP channel over mScarlet channel. **L) mSc::HSP-1 colocalizes with GFP::IFD-1.** HSP-1/Hsc70 is essential for recognizing ubiquitinated aggregates for aggresomal collection in mammalian biology ^36, 89^. Representative of adult touch neuron expressing *bzSi53*[P_mec-7_mSc::HSP-1]; *bzIs3*[P*_mec-7_*GFP::IFD-1]. Representative of N > 10. Scale bar = 2 *µm*. Quantitation of co-localization of N = 10, graphed as Mander’s overlap coefficient of GFP channel over mScarlet channel.

We tested for microtubule and dynein roles in IFD-1 puncta formation (Figure 2B-C). To test microtubule (MT) requirements for juxtanuclear IFD concentration, we treated P*_mec-7_*GFP::IFD-1 expressing animals with the potent microtubule inhibitor colchicine^32, 33^ from L4 to Ad2 (Figure 2B). We find that colchicine increases the number of dispersed GFP::IFD-1 foci (Figure 2D), reduces their size (Figure 2E), and disrupts their juxtanuclear positioning (Figure 2F), suggesting a requirement for MTs in the development and juxtanuclear localization of the *C. elegans* IFD compartment, as occurs for mammalian aggresomes.

Colchicine treatment acts systemically and could act via non-autonomous influences. To address cell-intrinsic requirements, we used the auxin-inducible degradation system^34^ to degrade dynein heavy chain DHC-1 specifically in the touch neurons. We introduced a degron-tagged *dhc-1* allele into a strain engineered for touch neuron-specific expression of the ubiquitin ligase adapter TIR1, which promotes rapid degradation of degron-linked sequences when auxin is added to the culture. We treated animals with auxin from L4 to Ad2 to enable auxin-promoted degradation of endogenous DHC-1::degron and scored GFP::IFD-1 inclusions (Figure 2C). We found that in the presence of auxin (i.e., when DHC-1::degron was touch neuron depleted), IFD inclusions were more numerous (Figure 2G), smaller (Figure 2H), mobile, and more dispersed (Figure 2I) as compared to non-knockdown DHC-1(+) controls. We conclude that a DHC-1-containing dynein motor acts cell autonomously in IFD collection and concentration into large juxtanuclear inclusions, similar to dynein motor roles in mammalian aggresome formation.

We also tested for molecular associations of IFDs with homologs of aggregate adapter proteins documented to contribute to, and associate with, mammalian aggresomes. Mammalian aggregate collection and delivery to aggresomes occurs via multiple distinct molecular adapters that link aggregates to dynein motors for MT transport to juxtanuclear IF-rich inclusions,^4, 31, 35, 36^ including the SQSTM1/p62 autophagy/adapter, HDAC6 deacetylase, and 14-3-3 protein, the latter of which functions in an adapter complex that includes BAG3 and chaperone Hsc70^36^ (See Figure 2A). We constructed strains harboring GFP::IFD-1 and red tagged *C. elegans* homologs of *sqst-1/*SQSTM1, *hsp-1*/Hsc70 and *ftt-2/*14-3-3. For each adapter tested, we find colocalization with the juxtanuclear GFP::IFD-1 protein, akin to their characterized mammalian counterparts (Figure 2J-L).

To test the possibility that aggresome adaptors are needed to form IFD-aggresome like structures, we examined available viable mutants for *C. elegans* counterparts of three aggregate collection pathways: *hda-6*/HDAC6, *sqst-1/*SQSTM1, and *ftt-2*/14-3-3. We scored aggresome-like organelle dimensions as reported by GFP::IFD-1 in adapter mutant backgrounds (Supplementary Figure 3). Notably, we find that no individual adapter disruption eliminates the IFD compartment. Thus, no single aggregate collection pathway appears essential for aggresome-like organelle formation, and multiple routes likely function in parallel for cargo delivery, consistent with adapter requirements reported for mammalian models.^36, 37^ We did quantitate some changes in aggresome-like organelle dimensions in *sqst-1(ok2892),* in which ALM touch neurons have significantly smaller GFP::IFD-1 puncta (Supplementary Figure 3A), with increased puncta numbers (Supplementary Figure 3B). In addition, Ad1 *ftt-2(n4426)* ALM somata have significantly smaller GFP::IFD-1 aggresome-like organelles without a change in number of aggresome-like organelles per neuron (Supplementary Figure 3C-D). *hda-6*/HDAC6 deletion did not change size or number of GFP::IFD-1 Supplementary Figure 3E,F). The changes we identified in the GFP::IFD-1 puncta are consistent with roles for SQST-1 and FTT-2 in aggresome-like organelle filling.

### *C. elegans* aggresome-like organelles concentrate ubiquitin and include neurotoxic HttPolyQ aggregates

Mammalian aggresomes can collect ubiquitinated proteins^4, 38^ and therefore we also asked whether IFD inclusions colocalize with sites of ubiquitin concentration. We co-expressed single copy transgene mNeonGreen::UBQ-2 in touch neurons together with mScarlet::IFD-1 (Figure 3A) or mScarlet::IFD-2 (Figure 3B) and found that although some fluorescently tagged ubiquitin is dispersed or concentrated elsewhere in the neuronal soma as expected, mNG-tagged ubiquitin also concentrates in the IFD-1 and IFD-2 inclusions. We conclude that, like mammalian aggresomes, the *C. elegans* neuronal juxtanuclear IFD inclusions are associated with concentrated ubiquitinylated proteins. Together, our data highlight similar homology of biogenesis and composition between *C. elegans* neuronal IFD structures and mammalian aggresomes.

**Figure 3.**
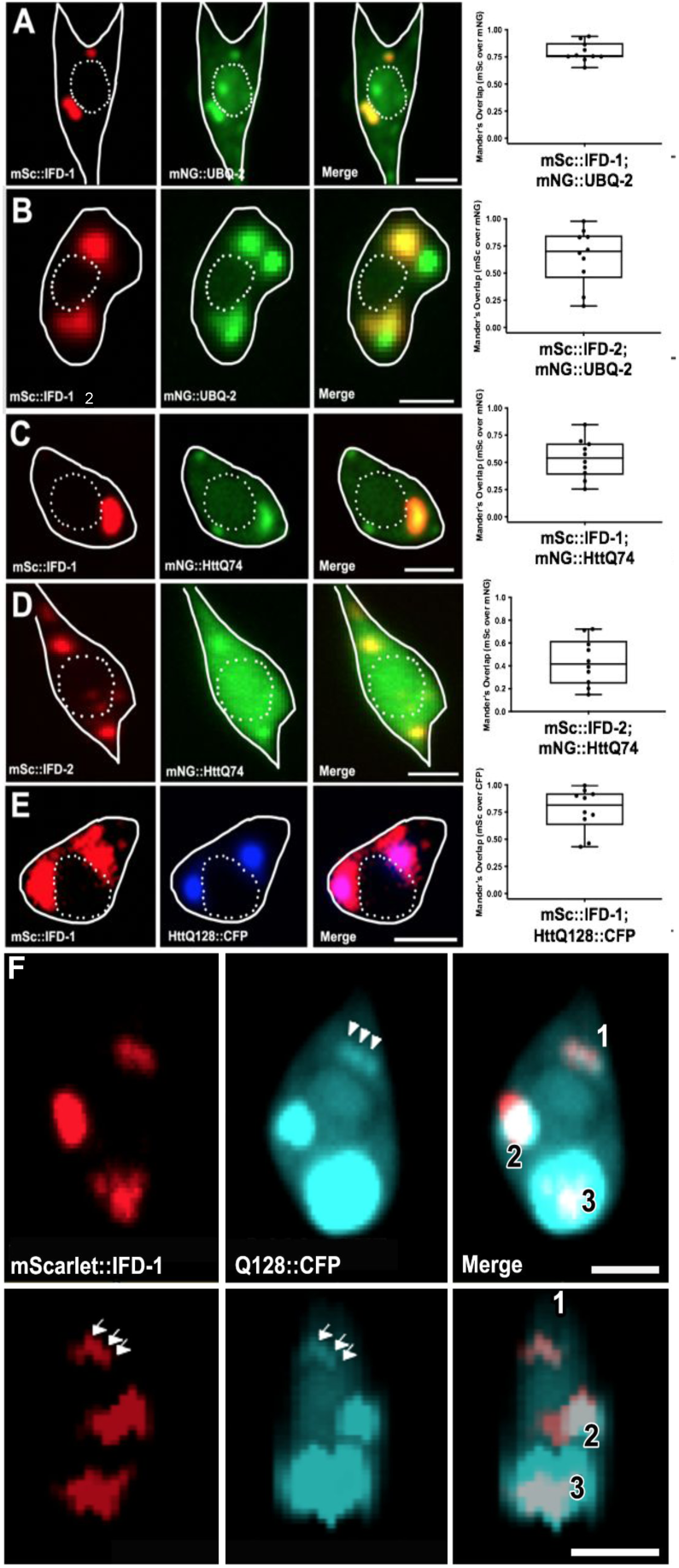
Disease-associated human Htt-polyglutamine expansion protein dynamically colocalizes with IFD-1 in touch neurons. White solid outline indicates the soma cell body and white dashed line outlines the nucleus in most panels. **A) mNG::UBQ-2 ubiquitin expressed from a single-copy-number transgene colocalizes with mSc::IFD-1.** *ubq-2* includes a canonical ubiquitin fused to the L40 ribosomal large subunit protein; the single ubiquitin polypeptide generated by cleavage can covalently attach to target proteins ^90^. We expressed mNG::UBQ-2 from single copy *bzSi38*[P*_mec-7_*mNG::UBQ-2], which expresses in touch neurons. mNG::UBQ-2 concentrates to specific subcellular structures including perinuclear IFD-1 and additional foci that are distinct, which would be expected for the range of activities/sites in which ubiquitin is engaged, such as on late endosomes. Adult touch neuron expressing *bzSi38*[P*_mec-7_*mNG::UBQ-2]; *bzSi34*[P*_mec-7_*mSc::IFD-1] image is representative of N > 10. Scale bar = 2 µm. Quantitation of co-localization of N = 10, graphed as Mander’s overlap coefficient of mScarlet channel over mNeonGreen channel. **B) mNG::UBQ-2 ubiquitin colocalizes with mSc::IFD-2.** Representative adult touch neuron expressing *bzEx422*[P*_mec-7_*mNG::UBQ-2]; *bzEx421*[P*_mec-7_*mSc::IFD-2]. mNG::UBQ-2 concentrates to specific subcellular structures including perinuclear IFD-2 and additional foci that are distinct, which would be expected for the range of activities/sites in which ubiquitin is engaged, such as on late endosomes. Image is representative of N > 10. Scale bar = 2 µm. Quantitation of co-localization of N = 10, graphed as Mander’s overlap coefficient of mScarlet channel over mNeonGreen channel. Although we cannot rule out that high expression drives the mNG::UBQ-2 to the IFD domain, the compartmentalization to specific subcellular localizations is a pattern reported in other ubiquitin studies ^91, 92^. **C) HttQ74::mNG expressed from a single-copy-number transgene colocalizes with mSc::IFD-1.** Representative adult touch neuron expressing *bzSi3*9[P*_mec-7_*HttQ74::mNG]; *bzSi34*[P*_mec-7_*mSc::IFD-1]. Images selected were those in which mNG was visibly aggregated in the cell, as opposed to homogenous dim cytosolic signal. Representative of N > 10 with mNG aggregates. Quantitation of colocalization of N = 10, graphed as Mander’s overlap coefficient of mScarlet channel over mNeonGreen channel. Scale bar = 2 µm. **D) HttQ74::mNG expressed from a single-copy-number transgene colocalizes with mSc::IFD-2.** Representative adult touch neuron expressing b*zSi40*[P*_mec-7_*mNG::HttQ74]; *bzEx420*[P*_mec-7_*mSc::IFD-2] of animals that display visible mNG aggregation. Image is representative of N > 10. Scale bar = 2 µm. Quantitation of co-localization of N = 10 of animals that display visible mNG aggregation, graphed as Mander’s overlap coefficient of mScarlet channel over mNeonGreen channel. **E) HttQ128::CFP expressed from an integrated high-copy-number array colocalizes with mSc::IFD-1.** Representative adult touch neuron expressing *igIs*1[P*_mec-7_*YFP P*_mec-3_*HttQ128::CFP]; *bzSi34*[P*_mec-7_*mSc::IFD-1]. Representative of N > 10 with an overlapping localization. Scale bar = 2 µm. Quantitation of co-localization of N = 10, graphed as Mander’s overlap coefficient of mScarlet channel over mNeonGreen channel. **F) Localization patterns of HttQ128::CFP and IFD-1 interaction.** In this strain, HttQ128::CFP is expressed from a high-copy-number integrated transgene array. Expression is higher than in the single-copy HttQ74 strain, and the polyQ tract is longer. We note that the IFD compartment often does not appear to fully surround the HttQ128 signal in this background. Biophysical analyses support heterogeneity in aggregate properties *in vivo* in polyQ touch neurons ^71^. Top row: Adult touch neuron soma displaying the three main patterns of interaction we note in lines that express mScarlet::IFD-1 and HttQ128::CFP. 1) HttQ128 and IFD overlap entirely (white arrows). 2) HttQ128 and IFD display a bi-lobed globular interaction, where part of the bi-lobed structure has overlap of HttQ128 and IFD. 3) HttQ128::CFP signal fully encompass the IFD signal. Scale bar = 2 µm. Bottom row: side view z-stack (0.2 µm) for images in the top row. Scale bar = 2 µm. Strain: ZB5104 *igIs1*[P*_mec-7_* YFP, P*_mec-3_* htt57Q128::CFP, *lin-15(+)*]; *bzSi34*[P*_mec-7_*mScarlet::IFD-1].

Ubiquitin colocalization (Figure 3A-B) is suggestive of concentrated misfolded/aggregated proteins. To address whether a characterized toxic aggregating human neurodegenerative disease protein can concentrate in the IFD-aggresome-like inclusions, we focused on polyglutamine expansion proteins, which have been found to aggregate and stress *C. elegans* neurons.^39, 40^ We constructed and co-expressed a single copy transgene encoding mNG::HttQ74 (which includes much of the first exon of the human Huntingtin gene fused to a HttQ74 expansion^41^), together with mScarlet::IFD-1 or mScarlet::IFD-2 reporters. We identified strong localization of mNG::HttQ74 to IFD-1 and IFD-2 inclusions in the touch neuron soma (Figure 3C-D) when mNG::HttQ74 is expressed at levels at which visible aggregation is present. We also examined a published HttQ128::CFP line expressed from an integrated high-copy-number array along with mScarlet::IFD-1 to record coincident fluorescence (Figure 3E). Our data establish that expanded polyglutamine proteins can concentrate at IFD inclusions, consistent with the idea that IFDs mark and sequester a domain of aggregated protein, and similar to what has been reported for mammalian aggresomes.

Interestingly, we noticed that IFDs and HttQ128::CFP can adopt multiple relative configurations, with IFDs and polyQ aggregates being mostly coincident for smaller (presumptively newly formed) aggresome-like organelles in early adult life (Figure 3E), but more frequently with HttQ128::CFP exhibiting less complete IF “covering” with advancing time or when relative expression of HttQ128::CFP and IFDs appears unbalanced, with more HttQ128::CFP evident (Figure 3F, Supplementary Figure 4). Whether these structures correspond to IF domains overrun by high aggregate concentrations, or aggresomes dissociating for degradation, remains for future testing.

We also examined localization of the highly expressed mCherry reporter relative to IFD foci, which revealed a different pattern. Our previous work showed that mCherry *bzIs166*[P*_mec-4_*mCherry] expression increases proteostress and that mCherry can concentrate into subcellular foci, although juxtanuclear positioning was not apparent^7, 8^. We found dimmer mCherry concentrations colocalized with IFD-1 in some ALM neurons, but most highly concentrated mCherry in touch neuron somata was within LMP-1::GFP positive lysosomes (LMP-1 is the lysosomal membrane protein LAMP1 ortholog;^42^ (Supplementary Figure 5A-B). In summary, colocalization analyses indicate that proteins of different character (*i.e*., expanded poly-Q proteins and mCherry) can be differentially handled, concentrated, and/or stored in proteostressed neurons (Supplementary Figure 5), and that the expanded polyQ proteins associate with, and linger within, IFD inclusions in proteostressed neurons (Figure 3, Supplementary Figure 4).

### IFD intermediate filaments are required for efficient exopher production from proteostressed neurons

In human neurodegenerative disease, aggregates can transfer among neurons and glia to promote pathological spread.^13^ The *in vivo* spreading mechanism is poorly understood, and the extent to which mammalian aggresome pathways contribute to spreading mechanisms has not been investigated. Our mCherry model is well suited to address this question, as we previously showed that adult neurons in this background can selectively eject material in large (∼3.8μm) membrane-bound vesicles called exophers that can include protein aggregates.^7^ Exopher production begins with outward budding of a nearly soma-sized membrane-bound vesicle that can selectively include damaged organelles and toxic proteins^7, 8^ (Figure 4A-B). The dynamic process of exopher-mediated expulsion involves collection and localization of noxious cellular contents, neuronal soma expansion, outward budding of a large membrane-surrounded vesicle that can preferentially concentrate specific cell contents, and the release of the loaded membrane-surrounded nascent exopher into the domain of the neighboring cell.^7, 10^ Exopher contents can be delivered directly to neighboring tissues (the glia-like hypodermis, in the case of *C. elegans* touch-neuron-derived exophers that we have studied most).^10^ We have speculated that the *C. elegans* exopher mechanism may be analogous to the poorly understood process by which neurodegenerative disease-associated aggregates spread from cell to cell to promote pathology in human neurodegenerative disease.

**Figure 4.**
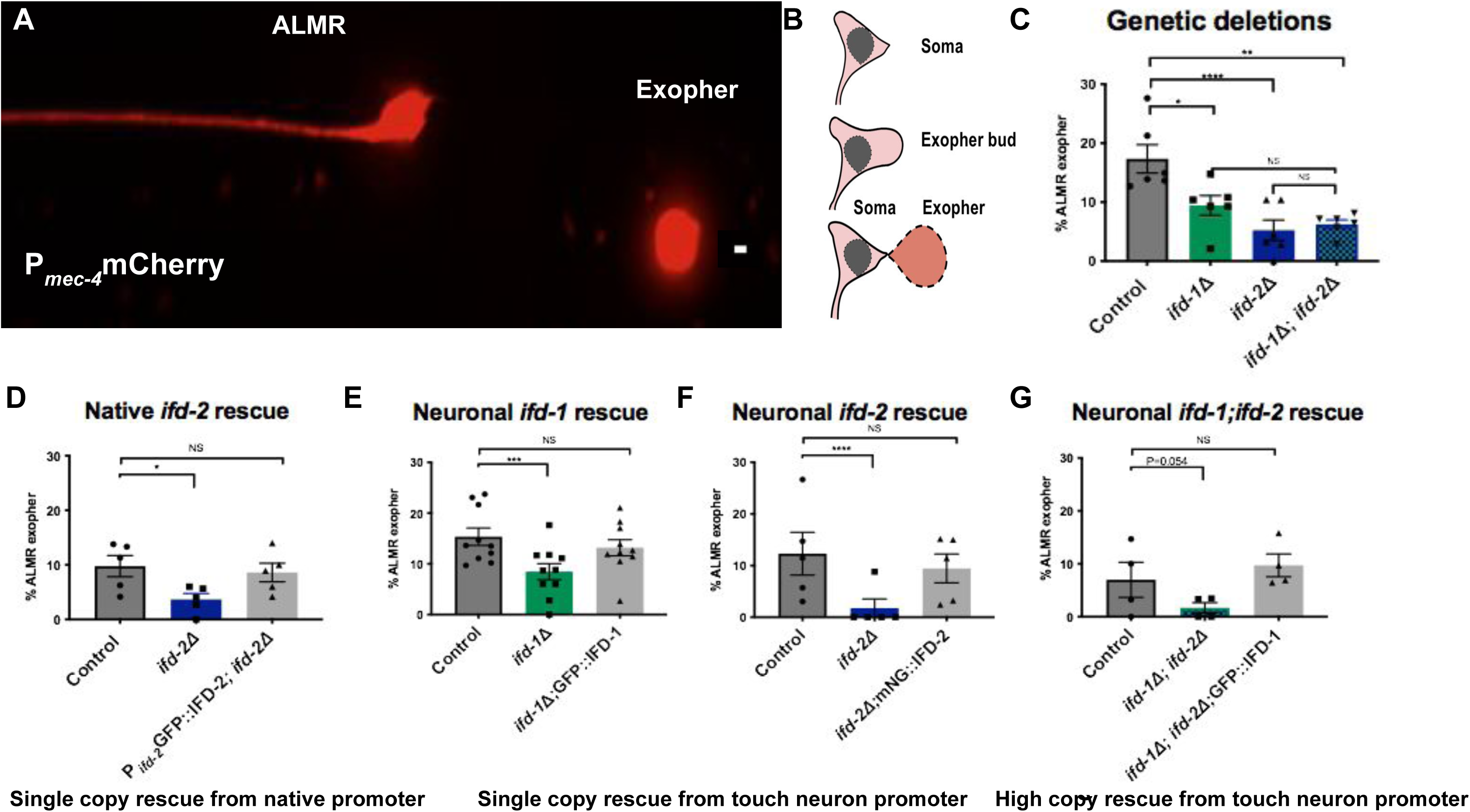
Intermediate filaments IFD-1 and IFD-2 act touch-neuron autonomously to support exopher production. Unless otherwise noted, exopher scoring throughout this study was evaluated using the Cochran-Mantel-Haenszel test (CMH), error bars are SEM; * P < 0.05, ** P < 0.005, *** P < 0.0005. **A) A typical exopher is readily visualized by mCherry** in strain ZB4065 *bzIs166*[P*_mec-4_*mCherry]. mCherry is a fluorophore expressed from an integrated P*_mec-4_* driven high-copy-number transgene array and induces exophergenesis. ALMR (**A**nterior **L**ateral **M**icrotubule, **R**ight side) mechanosensory touch neuron and an ALMR-derived exopher are pictured. Scale bar = 2 µm. Note: In general, several red fluorophores are known to aggregate in cells.^93^ (We report here that the mCherry, which is highly expressed in the *bzIs166*[P*_mec-4_*mCherry] strain, is useful as a touch neuron and exopher marker, but can concentrate in structures labelled with LMP-1::GFP and are thus presumptive lysosomes (See Supplementary Figure 5)). **B) Exophers form via a budding out mechanism.** Cartoon depiction of the exophergenesis process. 1) A normal neuronal soma depicted. 2) The neuronal soma swells, cellular contents polarize, and a swelling and budding out process generates a prominent exopher bud. 3) The exopher bud continues to extend outward; the exopher bud matures into a distinct exopher. The exopher can initially remain attached by a thin filament tube capable of transferring additional cell material; the tube commonly breaks later. **C) Deletion of *ifd-1* or *ifd-2* decreases ALMR exophers and the double mutant has exopher levels similar to single mutants**. We scored ALMR exopher production on Ad2 in the *ifd-1(ok2404)* deletion mutant, the *ifd-2(bz187)* deletion mutant, and the double *ifd* deletion mutant *ifd-1(*Δ*);ifd-2(*Δ), all of which harbored the exopher-inducing *bzIs166*[P*_mec-4_*mCherry] integrated transgene. The ALMR exopher percentage in the *ifd-1(*Δ*);ifd-2*(Δ) double mutant is not significantly different from the single mutant *ifd-1*(P = 0.351); double mutant compared to *ifd-2* P = 0.0952. ALMRs scored > 391/strain, 6 trials. **D) Single copy transgene of P*_ifd-2_* GFP::IFD-2 expressed from the native promoter rescues the *ifd-2* exopher deficit.** We expressed GFP::IFD-2 from the native *ifd-2* promoter in the mCherry background and scored for exopher levels. We compared ALMR exophers at Ad2 in *bzIs166*[P*_mec-4_*mCherry] compared to *bzIs166*[P*_mec-4_*mCherry]; *ifd-2(bz187) (P = 0.017). bzIs166*[P*_mec-4_*mCherry] compared to *bzIs166*[P*_mec-4_*mCherry]; *ifd-2(bz187); bzSi76*[P*_ifd-2_*GFP::IFD-2] (P = NS, CMS). N > 229, 5 trials; error bars are SEM. **E) Single copy P*_mec-7_*GFP::IFD-1 expressed in touch neurons rescues the exopher suppression phenotype of the *ifd-1(ok2404)* mutant.** We scored ALMR exophers at Ad2 in *bzIs166*[P*_mec-4_*mCherry] compared to *bzIs166*[P*_mec-4_*mCherry]; *ifd-1(ok2404)*. P*_mec-4_*mCherry is not significantly different compared to *bzIs166*[P*_mec-4_*mCherry];*ifd-1(ok2404);bzSi3*[P*_mec-7_*GFP::IFD-1] (P = 0.427). N > 552 ALMR/strain, 10 trials. **F) Single copy P*_mec-7_mNeonGreen*::IFD-2 expressed in touch neurons rescues the exopher suppression phenotype of the *ifd-2(bz187)* mutant.** We measured Ad2 exophers in ALMR, *bzIs166*[P*_mec-4_*mCherry] compared to *bzIs166*[P*_mec-4_*mCherry];*ifd-2(bz187). bzIs166*[P*_mec-4_*mCherry] is not significantly different from *bzIs166*[P*_mec-4_*mCherry];*ifd-2(bz187);bzSi37[*P*_mec-7_*mNeonGreen::IFD-2] (P = 0.40 CMS). N > 179 ALMR/strain, 5 trials. **G) High copy P*_mec-7_*RFP::IFD-2 expressed in touch neurons rescues the exopher suppression phenotype of the *ifd-1(ok2404)*;*ifd-2(bz187)* double mutant.** We scored ALMR exophers at Ad2 in *bzIs166*[P*_mec-4_*mCherry] compared to *bzIs166*[P*_mec-4_*mCherry]; *ifd-1(ok2404); ifd-2(bz187)* (P < 0.054). *bzIs166*[P*_mec-4_*mCherry] compared to *bzIs166*[P*_mec-4_*mCherry]; *ifd-1(ok2404); ifd-2(bz187); bzEx270*[P*_mec-7_*GFP::IFD-1 OE] is NS. N >128, 4 trials. Note: Fluorescently tagged IFD-1 and IFD-2 proteins confer functional complementation of exophergenesis and are therefore likely to localize to, and indicate the identity of, the native subcellular compartments in which they are bioactive.

Having shown that *C. elegans* neuronal aggresome-like organelles can store/associate with neurotoxic disease aggregates, we considered their potential roles in aggregate transfer biology. Potential intermediate filament requirements in exophergenesis were an initial focus given that in humans, circulating IFs are clinical biomarkers of human neurodegenerative disease,^43, 44^ and IFs are major components of α−synuclein-containing Lewy bodies that characterize Parkinson’s disease neuropathology^45, 46^ features that might intersect with extrusion biology and proteostasis balance.

We therefore tested whether null alleles of *ifd-1* and *ifd-2* act as exopher modulators. We obtained a mutant strain containing the *ifd-1(ok2404)* deletion (1863 base pairs deleted) and used CRISPR/Cas9 technology to generate deletion allele *ifd-2*(*bz187)*, which is missing the *ifd-2* start codon and first three *ifd-2* exons. The *ifd-1(Δ)* and *ifd-2(Δ)* mutants exhibited WT morphology (Supplementary Figure 6A-B) and brood size (Supplementary Figure 6C), as well as normal or near normal developmental timing (Supplementary Figure 6D-E) *ifd-2(Δ)* ∼5 hour average delay to L4 and lifespan (Supplementary Figure 6F-G). *ifd-1(Δ)* and *ifd-2(Δ)* mutants also exhibited WT osmotic stress survival (Supplementary Figure 6H), normal (*ifd-2(Δ))* or elevated oxidative stress sensitivity (for *ifd-1(Δ)*) (Supplementary Figure 6I); osmotic and oxidative stresses are associated with proteostasis disruption^47, 48^, and WT or modest suppression *(ifd-2(Δ))* of baseline *gst-4*::GFP expression (GST-4::GFP is a *daf-16*/FOXO/ SKN-1/NRF responsive glutathione-S-transferase reporter commonly used to report stress conditions^49^) (Supplementary Figure 6J-K). In terms of touch neuron-specific features, we recorded normal touch neuron response (Supplementary Figure 6L) and touch neuron morphology during aging (Supplementary Figure 6M). Likewise, in *ifd* null mutants, mCherry distribution and levels are not markedly changed (Supplementary Figure 6N-O). We conclude that *ifd-1* and *ifd-2* deletions do not confer substantial systemic physiological disruption in the whole animal; nor do *ifd-1* and *ifd-2* deletions markedly perturb the basic biology of the touch neurons in which we characterize exophergenesis.

We crossed the *bzIs166*[P*_mec-4_*mCherry] reporter into the *ifd-1(Δ)* and *ifd-2(Δ)* backgrounds and measured exophers at Ad2. We found that *ifd-1(Δ)* and *ifd-2(Δ)* each suppress exopher production (Figure 4C), establishing an impact of *C. elegans* IFDs in exophergenesis. We confirmed exopher suppression in *ifd-1(Δ)* and *ifd-2(Δ)* using alternative reporter cargo encoded by *uIs31*[P*_mec-17_*GFP]) (Supplementary Figure 7), and consequent to *ifd-1* and *ifd-2* RNAi knockdown in a pan-neuronal specific RNAi knockdown strain (Supplementary Figure 8A). Combined data define a role for *C. elegans* IFDs in normally promoting exopher levels in proteostressed neurons.

### IFD-1 and IFD-2 act in the same genetic pathway to promote exopher formation

Despite the fact that null alleles of *ifd-1* and *ifd-2* remove extensive protein coding sequences, neither *ifd-1(Δ)* nor *ifd-2(Δ)* fully eliminated exophers. To address whether *ifd-1* and *ifd-2* might act in the same, or in distinct, pathways in exophergenesis, we constructed an *ifd-1(Δ)*; *ifd-2(Δ)* double mutant and compared exopher levels to those in single *ifd* mutants (Figure 4C). We found that exopher production in the *ifd-1(Δ); ifd-2(Δ)* double mutant was suppressed to the same extent as in the single mutants, suggesting that *ifd-1* and *ifd-2* are likely to act in the same pathway to influence exopher production, and consistent with their co-localization in neurons (Figure 1D).

Some exopher production remains evident when both *ifd-1* and *ifd-2* are absent, suggesting that additional IF proteins might influence exopher production. Indeed, when both IFDs are absent, RNAi knockdown of other *C. elegans* IF genes in a strain sensitized for touch neuron RNAi suggests that more IF proteins may modulate TN exophergenesis (Supplementary Figure 8B). Given the relatively strong exopher suppression effects for *ifd-1(Δ)* and *ifd-2(Δ)*, the RNA-seq-documented *ifd* expression in touch neurons^24–26^, our initial extensive characterization of the fluorescently tagged IFD-1 and IFD-2 in TNs (Figure 1, Supplementary Figure 1), and the limited study of the biological roles of the *C. elegans* IFD genes to date^50–54^, we elected to focus on *ifd-1* and *ifd-2* contributions to exophergenesis in further studies we present here.

### Intermediate filament proteins IFD-1 and IFD-2 act cell autonomously in touch neurons to promote exopher production

To confirm that *ifd* mutations themselves, rather than unintended mutations present in the strain background, mediate the impact on exopher levels, we complemented *ifd(Δ)s* with cognate *ifd* transgenes. We used a single-copy GFP::IFD-2 transgene, expressed from the native *ifd-2* promoter, and tested for rescue of the *ifd-2(Δ)* phenotype. *bzSi76*[P*_ifd-2_*GFP::IFD-2] rescues the *ifd-2(Δ)* exopher deficit, confirming the *ifd-2* gene role in modulating exopher levels (Figure 4D). We relied on single copy touch neuron-specific expression of GFP::*ifd-1* crossed to *ifd-1(Δ)* to confirm rescue of the exopher phenotype (Figure 4E).

To address the question of whether *ifd-1* and *ifd-2* exert cell-autonomous roles for IFDs in neuronal exopher production, we expressed single-copy fluorescently-tagged reporters exclusively in the touch receptor neurons under the control of the *mec-7* touch neuron promoter. We found that single copy *bzSi3*[P*_mec-7_*GFP::IFD-1] rescues the exopher deficit phenotype of the *ifd-1(Δ)* mutant (Figure 4E). Similarly, single copy *bzSi37*[P*_mec-7_*mNeonGreen::IFD-2] rescues the exopher deficit phenotype of the *ifd-2(Δ)* mutant (Figure 4F). Touch-neuron-specific genetic complementation supports the interpretation that IFDs are required cell-autonomously for neuronal exophergenesis.

Because *ifd-2* is also expressed in the intestine (^52, 53^ and our observations not shown) and *ifd-2* mutants can have intestinal morphology defects,^53^ we also tested for rescue of the neuronal exopher phenotype when we expressed single copy mNeonGreen::IFD-2 only in the intestine (*bzSi45*[P*_vha-6_*mNeonGreen::IFD-2]). Comparing *ifd-2(Δ)* to *ifd-2(Δ)* P*_vha-6_*mNeonGreen::IFD-2, we find that intestinal expression of mNeonGreen::IFD-2 does not rescue the neuronal exopher phenotype of *ifd-2(Δ)* (Supplementary Figure 9) consistent with a predominant role for *ifd-2* in exophergenesis in the neuron (Figure 4E-G) rather than in intestine, although a trend toward partial rescue leaves open the possibility of some intestinal contribution. Overall, touch neuron-specific rescue of the exopher production deficits in *ifd* null mutants support that IFD-1 and IFD-2 primarily act autonomously within the touch neuron to influence exopher production.

### Over-expression of *ifd-1* or *ifd-2* does not markedly change exophergenesis

Our gene manipulations of *ifd-1* and *ifd-2* also enabled us to address consequences of high level expression of *ifd-1* and *ifd-2* on exopher levels (Supplementary Figures 10, Supplementary Figure 11). In brief, over-expression of functional *ifd-1* or *ifd-2* constructs in a *ifd*(+) background does not drive exopher levels significantly higher or lower than controls (Supplementary Figure 10). Lack of over-expression effects (Supplementary Figure 10) are consistent with our observation that size of the aggresome-like compartment *per se* does not correlate with generally high exopher levels nor does aggresome-like organelle size predict extrusion (Supplementary Figure 12).

We find some capacity for cross-complementation by high copy *ifd* expression (i.e., *ifd-1(Δ))* by *ifd-2*OE; Supplementary Figure 11). Thus, despite the clear genetic requirements for native levels of both *ifd-1* and *ifd-2* for normal exopher production levels (Figure 4C, Supplementary Figure 7B), the partial functional substitution conferred by cross-overexpression supports similar bioactivity for *ifd-1* and *ifd-2*. (Figure 4G, Supplementary Figure 11A).

### Adapter proteins SQST-1, HDA-6, and FTT-2 modulate exopher production

Having established that *ifd-1* and *ifd-2* can modulate exopher levels, we sought additional evidence of aggresome component impact on exophergenesis. We scored exopher levels in available mutants for genes encoding aggresome adaptor proteins that function in aggresome loading of cargo and colocalize with IF-organelles (Figure 2J-L, Supplementary Figure 3), appreciating at the outset that the genes we tested encode multi-tasking proteins that contribute in multiple facets of proteostasis.

We found that *sqst-1(ok2892)* increases variation in exopher levels, (Supplementary Figure 13A), without conferring a statistically significant change. *hda-6(ok3203)* (which did not significantly impact GFP::IFD-1 formation (Supplementary Figure 3A-B)), increased exopher levels (Supplementary Figure 13B).

We found that the 14-3-3 mutant *ftt-2(n4426)* exhibits a clear exopher suppressor phenotype on Ad1, implicating FTT-2 in exopher production (Supplementary Figure 13C). To begin to address cell autonomy of FTT-2/14-3-3 for touch neuron exopher production, we induced *ftt-2*(RNAi) knockdown in both neuron-specific and touch neuron-sensitized RNAi backgrounds. Our data reveal a suppression of exopher formation when *ftt-2* is targeted for knockdown in neurons (Supplementary Figure 13D-E). We also introduced a single copy of *ftt-2* expressed from the touch neuron *mec-7* promoter (P*_mec-7_* FTT-2::mSc; P*_mec-4_* mCherry) in an otherwise WT *ftt-2(+)* background. When we tested exopher levels in this strain, which is wild type *ftt-2* at the endogenous genomic locus and harbors a second copy of P*_mec-7_* FTT-2::mSc inserted via MiniMos single site insertion^55^ (*i.e.,* at least 2X *ftt-2* gene dose), we quantitated significantly increased TN exopher levels (Supplementary Figure 13F). Data support cell autonomous activity of FTT-2 in touch neuron exopher modulation and suggest that enhanced FTT-2 activity can drive exopher formation.

To assess requirements for the BAG3/HSP-1/14-3-3 adaptor complex in exopher biology from a second angle, we knocked down *hsp-1/Hsc70*^29, 36^ in a touch neuron-RNAi sensitized background (Supplementary Figure 13G-H). Exopher production scores suggest that the FTT-2 aggresome-related binding partner HSP-1 plays a role in touch neuron exophergenesis and further support that an FTT-2/HSP-1 complex can act to modulate exopher levels.

In sum, genetic perturbations of IF genes *ifd-1* and *ifd-2*, and aggresome adaptor genes *ftt-2* and *hsp-1*, suppress exopher levels demonstrating a potential link between these aggresome components and exopher formation.

### Aggresome-like foci can be expelled within exophers, but are not mandatory cargo of exophers

Having established that specific aggresome components likely act cell autonomously to influence exopher levels, we asked whether aggresome-like inclusions can themselves be expelled in exophers, and if so, whether aggresome inclusion is mandatory for exopher formation. We first examined exophers produced by neurons co-expressing mCherry and GFP::IFD-1 to find that exophers can contain extruded GFP::IFD-1 foci in this background (Figure 5A; GFP::IFD-1 foci in 11/61 exopher events, Figure 5B). With regard to the capacity to fully clear neurons of the aggresome-like structures, in a separate experiment, we find that 4/9 exopher events that include an aggresome-like foci, the complete visualizable GFP::IFD-1 aggresome-like organelle contents of the soma were ejected; in the remaining 5 cases, one GFP::IFD-1-aggresome-like structure was retained in the soma and one was extruded in the exopher (Figure 5C).

**Figure 5.**
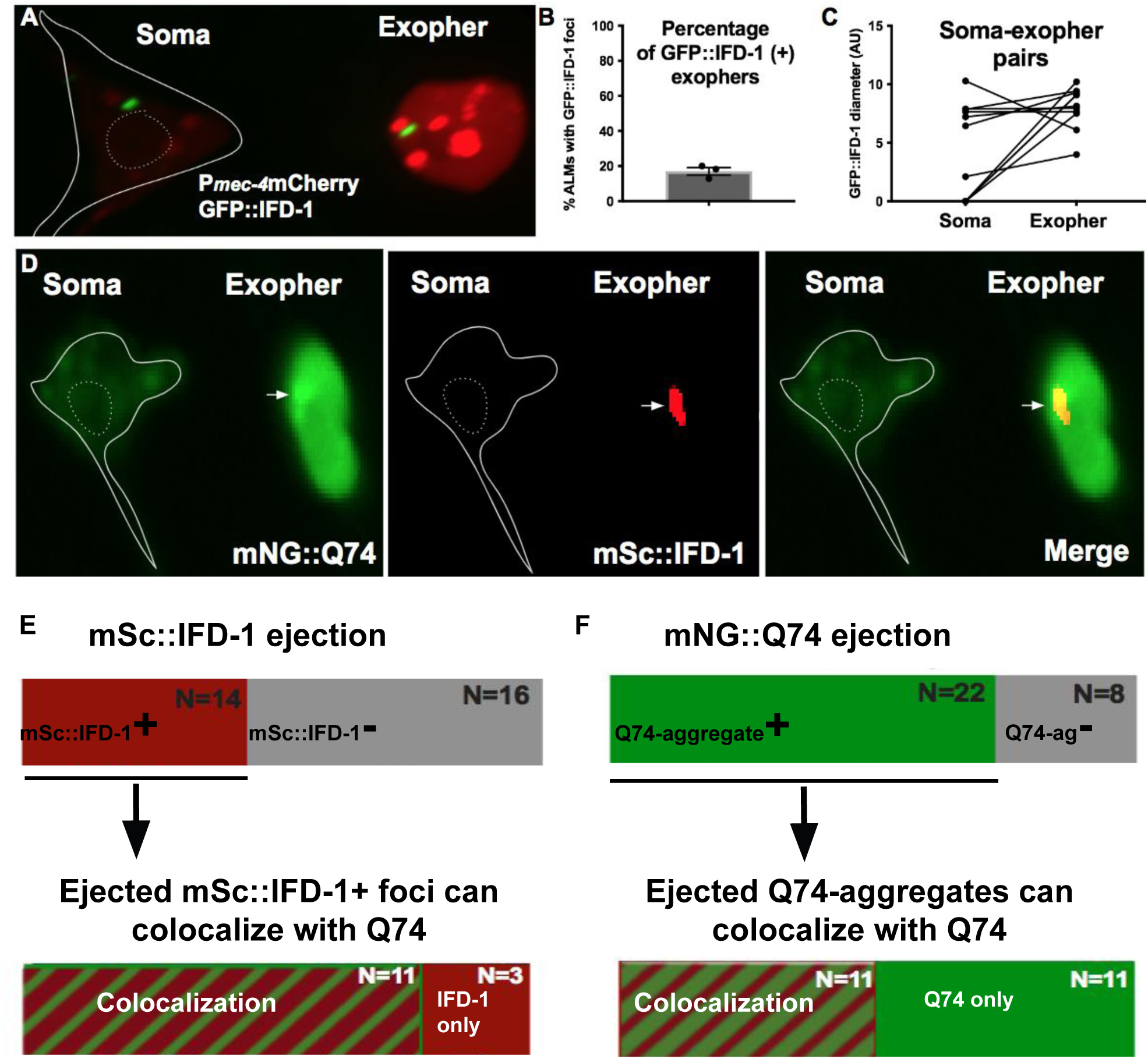
IFD proteins can be extruded in exophers. **A) IFD-1-positive puncta extruded in an exopher** in a strain that expresses *bzIs166*[P*_mec-4_*mCherry] for touch neuron/exopher visualization. Shown is an adult touch neuron from strain ZB4632 *bzIs166*[P*_mec-4_*mCherry];*bzSi3*[P*_mec-7_*GFP::IFD-1]. Soma outline is in white, nucleus is outlined with a white-dashed line. Arrowhead points to the concentrated GFP::IFD-1 organelle, one GFP::IFD-1 focus remains in the soma, and one is ejected in the exopher. Scale bar = 2 um, representative of N > 10. **B) IFD positive organelles are included in ALM exophers.** We examined strain ZB4632 *bzIs166*[P*_mec-4_*mCherry]; *bzSi3*[P*_mec-7_*GFP::IFD-1] on Ad3 and determined how often GFP::IFD-1 was extruded in exophers. In total, we observed 61 ALM Ad3 exopher events, with eleven having GFP::IFD-1 included in the exopher, 3x trials. **C) The neuron can eject the entire GFP::IFD-1-positive aggresome-like organelle**. To ask if the neuron ejected the largest GFP::IFD-1 collection of the cell, we looked at neurons that ejected a GFP:IFD-1. Of nine neurons in which GFP::IFD-1 was expelled in the exopher, four neurons ejected the only – and entire - GFP::IFD-1 organelle visibly present. Note: Excluding datapoints in which there were no foci that remained in the soma, the size of the GFP::IFD-1 that remained in the soma versus the size of the foci that were expelled did not indicate statistically significant difference. Data show that a cell can eject the entire IF-organelle-compartment, and suggest that there is no obvious size difference between the ejected aggresome versus the aggresome that remains in the neuron. **D) IFD-1-positive puncta extruded in an HttQ74-enriched exopher.** Shown is an adult touch neuron from strain ZB5171 *bzSi39*[P*_mec7_*HttQ74::mNG]; *bzSi34*[P*_mec-7_*mScarlet::IFD-1] on Ad1 after a 6 hour fast. Soma outline is in white, nucleus is outlined with a white dashed line; arrowhead points to the concentrated HttQ74 punctum that co-localizes to the IFD-1 domain. Scale bar = 2 µm, N > 10. Note that in this image much of the cytosolic mNG::HttQ74 gets extruded into the exopher domain, but there is clear concentration in the mSc::IFD-1-compartment as well. **E) IFD-1-positive organelles are included in 14/30 ALM exopher events, and most of the ejected IFD-1 puncta (11/14) colocalize with mNG::HttQ74.** We examined strain ZB5171 *bzSi39*[P*_mec7_*HttQ74::mNG]; *bzSi34*[P*_mec-7_*mScarlet::IFD-1] on Ad1 after a 6 hour fast to increase exopher yields, and asked how often mScarlet::IFD-1 was extruded in exophers. We observed 30 ALM exopher events, with 14 having mSc::IFD-1 included in the exopher. Next we asked how often the 14 mScarlet::IFD-1-positive exopher events also included at least one concentrated HttQ74 puncta overlapping with mSc::IFD-1. In 11/14 ALM IFD-1-positive exophers, there was at least one mNG::HttQ74 punctum colocalized with mSc::IFD-1. **F) HttQ74-aggregates are included in 22/30 ALM exopher events, and half of the ejected HttQ74-aggregates colocalize with mSc::IFD-1.** We examined the same strain and the same exopher events as described above. ZB5171 *bzSi39*[P*_mec7_*HttQ74::mNG]; *bzSi34*[P*_mec-7_*mScarlet::IFD-1] on Ad1, after a 6 hour fast to increase exopher yields. We determined how often HttQ74 aggregates were extruded in exophers. Out of N = 30 ALM exopher events, 22 exopher events had at least one HttQ74 aggregate included in the exopher. We asked how often mNG::HttQ74 positive exopher events (N = 22) included species where HttQ74 is a cargo of the mSc-aggresome-like organelle to find 50% (11/22) of ALM HttQ74-positive exophers had at least one incidence of HttQ74 including mSc::IFD-1.

Since IFDs colocalize with HttPolyQ aggregates in touch neurons (Figure 3C-F), we also examined exophers in a strain that co-expressed mSc::IFD-1 and mNG::HttQ74 (Figure 5D). In these studies, we scored exophers after 6 hours of food withdrawal, a culture condition that increases exopher production^23^. Of 30 total exopher events, 14 exophers included detectable mSc::IFD-1, establishing that tagged intermediate filament mSc::IFD-1 can be extruded in exophers in nutrient-stressed mNG::HttQ74-expressing neurons (Figure 5E, top) (image of polyQ and mSc::IFD-1 extrusion in Figure 5D). Our observation that mSc::IFD-1 and GFP::IFD-1 are included in some, but not all, of the exophers formed in the two different backgrounds (∼15% mCherry; ∼ 50% HttQ74), make the important point that significant IFD-1 extrusion is not a mandatory requirement in all exophergenesis events.

### Ejected IFD foci colocalize with mNG::HttQ74 in most extrusion events

To assess the IFD-1 status relative to ejected cargo, we measured the frequency of mNG::HttQ74 overlap with the mSc::IFD-1 that were ejected in exophers (*i.e*., in exophers that included mSc::IFD-1 puncta/ae). Of the 14 mSc::IFD-1-positive exophers we reported in Figure 5E (top), 11 exophers contained species in which mSc::IFD-1 and mNG::HttQ74 colocalized to the same puncta; only 3 exopher events had mSc::IFD-1 foci in exophers(s) that lacked co-enrichment of mNG::HttQ74 (Figure 5E, bottom). We conclude that IFD-1-aggresome-like organelles hosting HttQ74-cargo are often components of extruded exophers.

In a complementary analysis alternatively focused on cases in which HttQ74 aggregates leave the cell, we looked at the same 30 exopher events with an initial scoring of the mNG::HttQ74 reporter, and later checked for co-association of the IFD reporter. We found that 22 of the 30 mNG::HttQ74-identified exophers contained at least one identifiable mNG::HttQ74-enriched punctum (Figure 5F, top). Of those 22 exopher events that included mNG::HttQ74 punctae, we asked whether the ejected HttQ74 was accompanied by IFD-1. We found 11 out of 22 exophers also displayed mSc::IFD-1 colocalized with mNG::HttQ74 (Figure 5F, bottom). Data indicate that HttQ74-aggregates are often extruded into exophers in IFD-1+ aggresome-like organelles, although HttQ74 can leave the soma in the exopher compartment nearly as frequently without co-IFD-1 association. We conclude that IFD-1 association is not strictly required for HttQ74 extrusion.

### Human neuronal intermediate filament protein hNFL can partially complement *C. elegans ifd-2* deletion to promote exopher formation

Mammalian intermediate filament proteins are integrated into the neuronal proteostasis network and can associate with juxtanuclear aggresomes^4, 56–58^. Roles for mammalian IFs in neuronal aggregate extrusion have not been reported, but our *C. elegans* observations raise the possibility that mammalian neuronal IFs might act in exopher-related biology^14, 19^. To begin to address this possibility, we asked if a human neuronally-expressed intermediate filament protein can complement an *ifd(Δ)* for exopher production deficits.

We focused on testing human neurofilament light chain (hNFL) because human NFL is clinically evaluated as a circulating blood/CSF biomarker of neuronal injury and neurodegenerative disease^43, 44^, despite mysteries surrounding the mechanistic origins of NFL that circulates. hNFL is a neuron-specific intermediate filament protein and exhibits ∼24/45% sequence identity/similarity to *C. elegans ifd-1* and *ifd-2*.

We optimized the human NFL gene for expression in *C. elegans* based on codon bias and inclusion of short introns using *C. elegans* Codon Adapter Software^59^. We first documented the subcellular distribution of the tagged mNG::hNFL expressed under the direction of a *mec-7* promoter from a single copy insertion in touch neurons. (Supplementary Figure 14). mNG::hNFL concentrated to juxtanuclear foci that colocalize with *C. elegans* intermediate filament reporter mSc::IFD-1 (Figure 6A). mScarlet::hNFL colocalizes with ubiquitin, mNG::UBQ-2 (Figure 6B), and with mNG::HttQ74 (Figure 6C) in aggresome-like compartments. Although the predominant localization of mNG::hNFL is to one or two punctate juxtanuclear locations (Supplementary Figure 14) with both single copy and high copy mNG::hNFL reporters, we did note that FP-tagged hNFL could also be distributed more broadly in the neuron as compared to *C. elegans* FP::IFDs, also appearing in the axon, nucleus, and in filamentous patterns in the cytoplasm in some neurons (Supplementary Figure 14A-C). Nonetheless, *C. elegans* IFDs and hNFL clearly localize similarly to aggresome-like compartments (Figure 6A). Akin to what we documented for the *C. elegans* IFD-aggresome-like organelle (Figure 5), we found that hNFL inclusions are expelled within approximately half of *C. elegans* exophers (Figure 6D).

**Figure 6.**
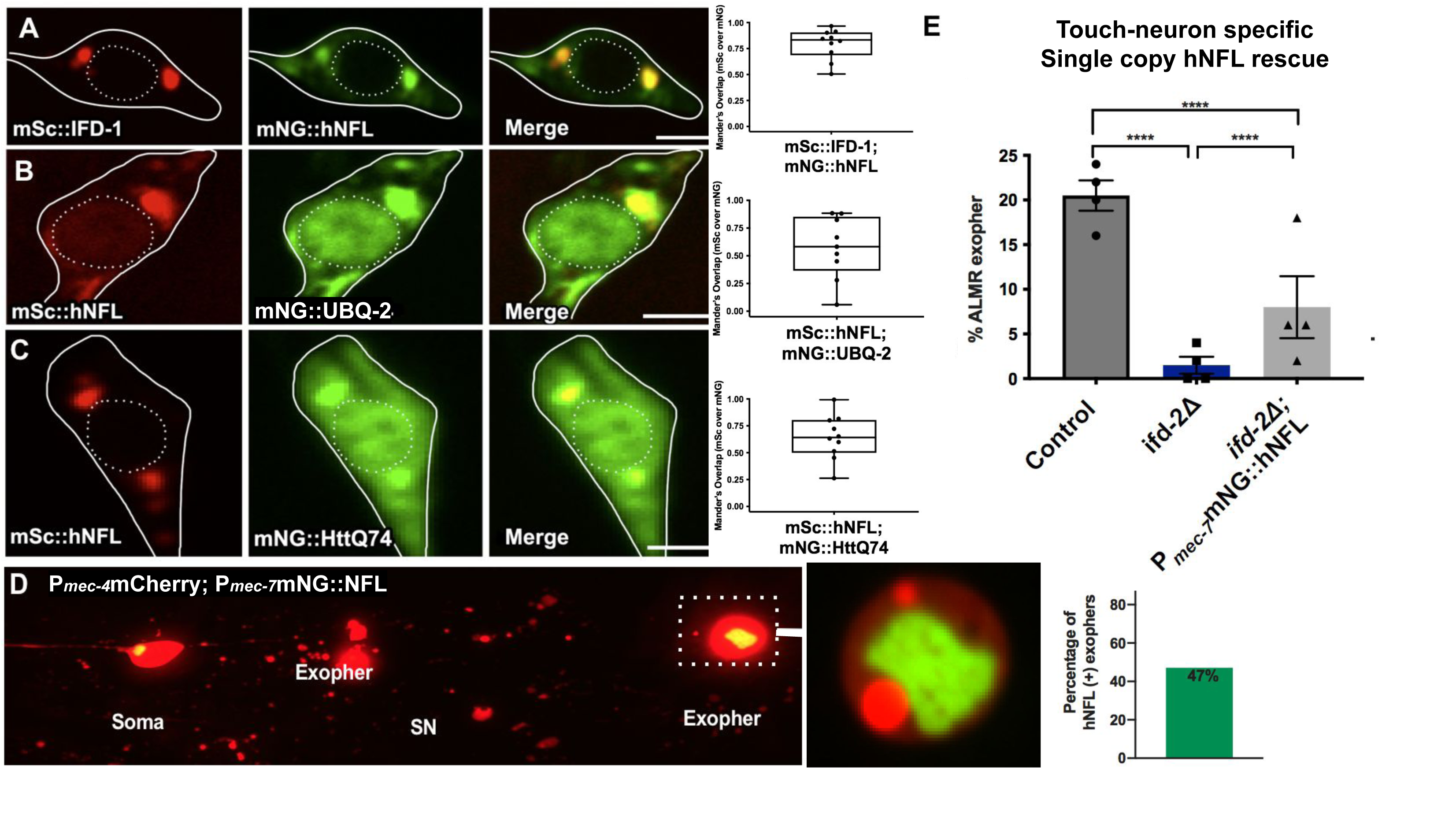
IF roles in aggregate collection and neuronal aggregate expulsion may be partially conserved. **A) mNG::hNFL localizes to the IFD-1 positive aggresome-like organelle.** Shown is a representative ALM touch neuron. We optimized the coding sequence of human Neurofilament Light Chain, hNFL, NM_006158.5, for *C. elegans* expression based on published codon biases as described in Methods. We added mNeonGreen fluorophore and expressed the construct under the *mec-7* touch neuron promoter. Strain is expressing *bzEx269*[P*_mec-7_*mNG::hNFL] and *bzSi34*[P*_mec-7_*mSc::IFD-1]. Solid white lines outline the touch neuron soma, dotted outline is the nucleus. Image is representative of N > 20. Scale bar = 2 µm. Graphed as Mander’s overlap coefficient of mScarlet channel over mNeonGreen channel, N = 10 graphed, error bars are SEM. **B) mSc::hNFL colocalizes with mNG::UBQ-2 ubiquitin.** Representative adult touch neuron expressing *bzEx313*[P*_mec-7_*mNG::UBQ-2]; *bzSi43*[P*_mec-7_*mSc::hNFL]. Image is representative of N > 10. Scale bar = 2 µm. Quantitation of co-localization of N = 10, graphed as Mander’s overlap coefficient of mScarlet channel over mNeonGreen channel. **C) mSc::hNFL colocalizes with mNG::HttQ74.** Representative adult touch neuron expressing single copy *bzSi39*[P*_mec-7_*mNG::HttQ74]; *bzSi43*[P*_mec-7_*mSc::hNFL]. Image is representative of animals that display visible mNG aggregation, N > 10. Scale bar = 2 µm. Quantitation of co-localization of N = 10 of animals that display visible mNG aggregation, graphed as Mander’s overlap coefficient of mScarlet channel over mNeonGreen channel. **D) hNFL-labeled aggresome-like foci can be extruded from touch neurons in exophers.** Representative image (N > 10) of mNG::hNFL extrusion in exophers. Strain expressing *bzEx311*[P*_mec-7_*mNG::hNFL] with *bzIs166*[P*_mec-4_*mCherry] in the *ifd-2(bz187)* background. Image shows ALMR neuron with three exophers. Two exophers are positioned in the middle of the image surrounded by scattered mCherry debris, starry night phenotype (SN) from hypodermal exopher digestion ^7^. The third exopher pictured, a mature distant exopher, contains a large mNG::hNFL inclusion, shown in focus and zoomed in. The exposure is altered to show details. **Graph on right:** Out of N = 34 Ad2 ALMs with an exopher from strain expressing *bzSi48*[P*_mec-7_* mNG::hNFL]; *bzIs166*[P*_mec-4_*mCherry], 16 exophers included mNG::hNFL puncta. **E) Touch neuron specific single copy P*_mec-_*_7_mNG::hNFL partially rescues the exopher suppression phenotype of the *ifd-2(bz187)* mutant.** We measured Ad2 exophers in ALMR. *bzIs166*[P*_mec-4_*mCherry] compared to *bzIs166*[P*_mec-4_*mCherry];*ifd-2(bz187)* and *bzIs166*[P*_mec-4_*mCherry];*ifd-2(bz187)* compared to *bzIs166*[P*_mec-4_*mCherry];*ifd-2(bz187);bzSi48[*P*_mec-7_*mNeonGreen::hNFL], both significantly different, P < 0.005, N = 200 ALMR / strain, 4 trials. * P < 0.05, ** P < 0.005, *** P < 0.0005. The images of the somas in each panel are not related.

Having documented distribution of human hNFL that reflects that of *C. elegans* IFDs at aggresome-like sites, we next wanted to investigate neuron-autonomous function of hNFL in exophergenesis. We introduced the P*_mec-7_*mNG::hNFL single copy transgene into an *ifd-2(Δ);* mCherry background and compared exopher production in *ifd-2(Δ)* vs. *ifd-2(Δ)*; P*_mec-7_*mNG::hNFL. Single copy P*_mec-7_*mNG::hNFL can partially rescue *ifd-2(Δ)* exopher deficits (P = 0.0046; Figure 6E). The mNG::hNFL single copy construct, similarly to tagged IFD constructs, does not significantly elevate or suppress exopher levels on its own, indicating that the rescuing activity is unlikely to be caused by NFL aggregation (Supplementary Figure 15). hNFL rescue of *ifd-2(Δ)* exopher deficits supports the proposition that human neuronal intermediate filament proteins might play a conserved role in exopher biology.

In sum, our finding that human hNFL shows overlapping subcellular localization with *C. elegans* IFD and aggresome-like organelle components (Figure 6A-C), is expelled from the soma in exophers (Figure 6D), and can partially compensate for *ifd-2* expulsion function *in vivo* (Figure 6E) raise the possibility that mammalian IFs may play roles in cellular extrusion that are conserved from nematodes to humans, inviting the re-examination of mammalian aggresome biology in the context of mechanisms of human aggregate spread.

## Discussion

*C. elegans* intermediate filament proteins IFD-1 and IFD-2 concentrate into stress-responsive juxtanuclear foci that are analogous in position and composition to what has been reported for mammalian aggresomes. IFD-1 and IFD-2 also act in proteostressed neurons to mediate efficient extrusion of cytoplasmic contents in large exopher vesicles (Summary in Supplementary Figure 16). Conservation of the associated biology is supported by our finding that human neuronal intermediate filament protein hNFL, expressed in *C. elegans* neurons, localizes to the IF-aggresome-like organelle, can partially functionally substitute for *C. elegans ifd-2* in exopher production, and can be extruded in exophers similarly to what we find for *C. elegans* IFDs. Our data hold implications for considerations of proteostasis, vesicle extrusion, and neuronal health.

### Stressed *C. elegans* neurons produce aggresome-like organelles

The original description of mammalian aggresomes (focused in cultured CHO and HEK cells) defined multiple aggresome features, including prominent juxtanuclear concentration of aggregates with distinct localization from Golgi and lysosomes, association with intermediate filaments under proteostress conditions (aggregate-prone protein over-expression or inhibition of the proteasome), colocalization with ubiquitinated and heterogeneous cargoes, and requirements for microtubule and dynein motor integrity for biogenesis.^4^ Follow up studies established that p62/SQSTM1(SQST-1) and components of the 14-3-3(FTT-2)/BAG3/Hsc70(HSP-1) complex associate with small foci that move into the large perinuclear aggresome-like organelle.^29, 60^ We find that most of these mammalian aggresome features are shared with the IFD-associated puncta in proteo-stressed *C. elegans* touch neurons, including collection of aggregated polyQ protein, ubiquitin enrichment, concentration of p62/SQSTM1(SQST-1) and 14-3-3(FTT-2)/Hsc70(HSP-1), and a requirement for intact microtubules and dynein activity to collect such materials into one or two juxtanuclear sites. Although aggresome-like structures associated with the expression of human neurodegenerative disease proteins^15, 61, 62^ or ROS stresses^21, 22^ have been reported for *C. elegans* cells, our studies constitute the first multi-component characterization of *C. elegans* neuronal aggresome-like organelles and definitively establish that aggresome biogenesis mechanisms are at least partially conserved from nematodes to humans. *C. elegans* proteostressed neurons thus can serve as a validated model for deciphering the relative role of aggresomes in maintaining neuronal proteostasis and health in an *in vivo* context.

### Intermediate filament and cargo dynamics in native context in a proteostressed neuron

Limited ultrastructural data inform on mammalian aggresomes^4, 6, 27, 28, 63–67^. Our *C. elegans* EM data for proteostressed neurons suggest an initial early association of short IFs within the concentrated central granular material positioned near the nucleus, followed by maturation featuring a loose discontinuous IF meshwork that surrounds proteinaceous content, eventually featuring longer more continuous fibers at the periphery (Supplementary Figure 2). Assembly of a loose cage suggests feasibility of the proposal that once the IF mesh assembles with initial aggregates to nucleate the aggresome compartment, cargo might be later able to diffuse directly into and around the aggresome.^68^

Our cell biological data report strong association of ubiquitin and polyQ aggregates with the aggresome-like IF compartment. A striking observation, however, is that as polyQ accumulates in the neuron over time, the IF/polyQ association progressively appears imbalanced such that the polyQ domain outgrows the IF domain, which can no longer surround or “cage” the polyQ domain (Supplementary Figure 4). Thus, under conditions of high expression of neurotoxic proteins, IFDs appear limited in their capacity to fully encase cargo like aggregated polyQ128, raising the possibility that imbalances in IF or aggregate levels might alter function/capacity of the IF-associated aggresome-related organelle. The cellular consequences of IF/aggregate proportion changes are not clear, and the question as to whether these structures correspond to an aggresome “uncoating” step ^69, 70^ prior to content degradation via autophagy remains for future study. Changing patterns of polyQ concentration have been reported in other *in vivo* studies in *C. elegans*^71^ and in HEK cells,^68^ suggesting dynamics may be more of a feature of aggresome biology and aggregate processing than previously thought.

### Aggresome components promote efficient exophergenesis

Our data show that aggresome-associated proteins that collect and concentrate with protein aggregates can modulate exophergenesis levels. In particular, the documentation of multiple IFs and aggregate collection factors (IFD-1/IF, IFD-2/IF, FTT-2/14-3-3, HSP-1/Hsc70) localized to the same soma compartment with shared functional capacity to modulate exopher extrusion levels, support an unanticipated intersection between aggresome-like organelle formation and exopher extrusion.

### Refining understanding of *in vivo* biology that influences exopher formation

Our genetic results address multiple hypotheses regarding the nature of the aggresome/exopher connection, suggesting how IFs might, and might not, modulate exophergenesis, raising the following points.

*1) IFDs exert neuron-intrinsic influences on exophergenesis.* Although *ifd-1* or *ifd-2* deficiency might be hypothesized to induce or inactivate a systemic stress response, we document multiple examples of touch neuron-specific rescue of exophergenesis deficits, and find few changes in stress and health indicators at the neuron or organism-wide levels in *ifd* null deletion backgrounds. Our data support a touch neuron-autonomous role in extrusion efficiency for IFD-1 and IFD-2, rather than general roles in animal or neuronal health.
*2) Aggresome dimension, per se, does not suffice to trigger expulsion.* Measures of the aggresome-like compartment relative to extrusion do not indicate a strong correlation of aggresome-like compartment dimensions with exopher levels (Figure 5C, Supplementary Figure 12). For example, aggresome-like compartment size increases with age but exopher production levels do not rise in a corresponding temporal pattern^7^ (Supplementary Figure 1B). We infer that the exophergenesis trigger is not exclusively activated by aggresome compartment size; if aggresome size influences expulsion, additional molecular conditions must also be met prior to exopher extrusion.
*3) Modulation of ifd expression levels is unlikely to drive exopher production.* While IFD-proteins are important for exopher production rate, IFD over-expression is not sufficient to further elevate exophergenesis. Thus IFD proteins are necessary but not sufficient to trigger exopher production (Supplementary Figures 10-11).
*4) IFs are not required partners for extruded aggregates or essential components of exophers.* Co-extrusion of IF and aggregates occurs in many, but not all cases, and thus IFDs do not appear to be essential structural components of all exophers (Figure 5). Thus IFs associated with aggresomes may act as part of an exopher triggering mechanism in the neuronal soma that acts prior to exopher production.

### Exopher-like mechanisms involving IFs might influence a range of disease processes and underlie the release of neurodegeneration biomarkers

Heritable mutations in neurofilaments can cause several neurological diseases including Giant Axonal Neuropathy, Charcot-Marie-Tooth (CMT) disease, Amyotrophic Lateral Sclerosis (ALS), and Parkinson’s disease (PD).^72, 73^ Neurofilaments have also been widely implicated in neurodegenerative disease pathology due to their concentrated presence within characteristic inclusions, such as Parkinson’s disease Lewey bodies.^46, 74, 75^ Our findings raise the possibility that IF-modulated content expulsion under neuronal stress conditions might be an unexpected contributor to disease pathology in these disorders. From another perspective, it is interesting that the presence of human neurofilament light protein, NFL, in CSF and blood is clinically measured as a biomarker for neuronal disease and injury.^74–76^ Exactly how hNFL is released into human fluid spaces is not well documented. We found that IFD-1 and IFD-2, sharing ∼ 24% identity and ∼ 43% similarity with human NFL, can be expelled within exophers in a significant fraction of exopher events. Thus, *C. elegans* exopher studies reveal a novel path through which IFs are released from proteo-stressed neurons and raise the possibility that the commonly used NFL biomarker of human neuronal injury and disease might also reflect active neuronal exopher expulsion mechanisms, a mechanism not generally considered. Potentially supporting this idea, intermediate filaments are also a major reported protein component of exopher-like extruded material from stressed cardiomyocytes.^14, 15^ Research into IFD or NFL expulsion in *C. elegans* may therefore both help explain basic mechanisms promoting aggresome- and Lewy body-like inclusion pathology that results in disease, and illuminate understanding of clinical findings used to assess pathology.

### Intermediate filaments in proteostasis and stress response

The identification of specific *C. elegans* intermediate filaments in a novel facet of proteostasis links to the increasing documentation of IF roles in proteostasis. Interestingly, studies on the mouse vimentin knockout reveal that IF protein vimentin is not necessary for either basic cellular function or aggregate collection at the juxtanuclear aggresome,^77^ similar to our observation in the *C. elegans* touch neuron model (Supplementary Figure 17). As in *C. elegans*, redundancy among intermediate filaments might mitigate effects. In mammals, vimentin has been implicated in stress responses that include heat shock, wound healing, and Alzheimer’s disease challenges.^78, 79^ Adult mammalian neural stem cells rely on vimentin for activation of proteostasis and aggregate elimination as the NSCs exit from quiescence.^77^ In the NSC model, vimentin is implicated in spatial organization of proteasomes and autophagosomes to the aggresome domain, possibly anchoring machinery that promotes degradation of aggresome contents.^77^ Detailed study of these processes in multiple models may further underscore commonalities and clarify mechanism.

### Aggresome-like organelles can be cleared from neurons by extrusion, holding implications for mammalian proteostasis

We document that IFD-marked aggresome-like foci can frequently leave the neuron together with cargo (Figure 5). In about half the cases in which IFD left in an exopher, the entire aggresome-like organelle content of the soma exits in the exopher; in most of the remaining cases we observe one aggresome-like organelle ejected and one left behind (Figure 5C). The point we underscore is that whole aggresome-like organelles can be removed from the neuron of origin for remote degradation, a novel concept in the field.

### Exophers may represent a conserved mechanism used to clear aggresomes

The fundamental mechanisms by which proteo-stressed neurons manage toxic aggregates and damaged organelles remain a central question in neurodegenerative disease.^13^ In mammals, processes strikingly similar to *C. elegans* exophergenesis have recently been described for HeLa cells,^80^ PC12 cells,^80^ and *in vivo* extrusions from mouse cardiomyocytes.^15, 17, 18, 81^ Moreover, *in vivo* transcellular extrusion of mitochondria from retinal ganglion cells for degradation by neighboring cells is reminiscent of exopher-like biology, and transcellular extrusion and degradation is increased in AD-model mouse brain and has been reported in human neurodegenerative disease brain.^82^ Emerging data thus highlight a newly identified cellular strategy for offloading aggregates and organelles for remote management that may operate across phyla. Although molecular links among the extrusion events across species remain to be rigorously established, initial observations are consistent with the hypothesis that exopher-like responses are within the capacity of a variety of stressed cells in a range of tissues across organisms (see review on heterophagy, ref. ^16^).

By extension, our findings that *C. elegans* exophers can clear aggresome-like organelles invite novel thinking on the cell biology of aggresomes and raise the possibility that aggresome-related mechanisms influence human neuronal aggregate spreading. Along these lines it is interesting that elevated expression of mammalian 14-3-3θ can promote alpha synuclein extrusion.^83^ This is akin to our finding that 14-3-3 protein FTT-2 is required for efficient exophergenesis and when modestly over-expressed can drive enhanced exopher production in *C. elegans.* Overall, our data invite a fresh look at how mammalian aggresomes are cleared and how exopher-like pathways contribute to spreading mechanisms.

## Additional information

Supplementary information is available for this paper.

## Acknowledgements

We thank past and present members of the Exopher Club for exopher-related discussions, and members of M.D. and B.D.G. laboratories at Rutgers University. We thank H. Ushakov for injections that created some transgenic lines. We thank Yunpeng Xu for sharing detailed control studies on neuronal RNAi efficacy. We thank Florian Geissler/Rudolf E Leube for providing plasmid DNA templates for IFD promoter amplification^84^ and Malene Hansen for neuronal RNAi strains with enhanced efficacy and RNAi-related discussions. Some strains were provided by the CGC (which is funded by NIH Office of Research Infrastructure Programs (P40 OD010440) and the OK *C. elegans* Deletion Mutant Consortium).

Research was supported by the National Institutes of Health under award numbers R37AG56510, R01AG047101, and 1F31AG066405. K.N. and D.H.H. were supported by NIH OD010943. Content is solely the responsibility of all authors and does not necessarily represent the official views of the National Institutes of Health.

## Author Contributions

M.L.A., J.F.C., R.A., S.A., I.M., A.J.S., and R.J.G., conducted and designed experiments, along with M.D., M.L.A., and B.D.G, who wrote the manuscript with input from J.F.C., R.A., and R.J.G.. G.B. provided some plasmid constructions. K.C.N. and D.H.H. carried out electron microscopy fixation, interpretation, and analysis.

## Data Availability

Correspondence and requests for materials should be addressed to Monica Driscoll via email.

**Reprints and permissions information are available at www.nature.com/reprints**

## Methods

### Strains and media

We cultured *C. elegans* strains at 20°C with standard methods as described ^8^. We fed animals *Escherichia coli* OP50 unless we initiated RNAi or a 6-hour fasting protocol to induce proteostress-exophers. Extrachromosomal arrays and drug-resistant strains were harbored on antibiotic selection plates (Hygromycin B or G418). For exopher measurements, L4 animals were removed from selection plates and placed onto standard OP50 plates for later observation.

### Strains

New transgenic alleles are provided in figure legends and in the strain list, Supplementary Table 1. Reporters for FP::IFD were expressed from the touch neuron specific *mec-7* promoter to focus on cell autonomous effects; single copy reporters from the native *ifd-1* and *ifd-2* promoters did not provide adequate neuronal signal for visualization. P*_mec-7_*FP::IFD constructs were effective at complementation of the cognate *ifd* alleles and were neither dominant negative nor exopher-inducing on their own. Animals expressing *igIs*1[P*_mec-7_* YFP P*_mec-3_*HttQ128::CFP] must be manually selected for CFP expression to avoid silencing of the CFP, which we have not been able to easily reverse (high copy number silencing has been documented ^85, 86^). PolyQ constructs are quite challenging to generate due to the extent of repeat sequences therein; still in an attempt to make and study a more co-operative polyQ in touch neurons, we constructed a single copy *bzSi39*[P*_mec-7_*mNG::HttQ74] allele. The polyQ protein aggregates minimally. We used CRISPR to generate AID degron-tagged alleles of *hsp-1* and *ftt-2* but found that these alleles were lethal (*hsp-1*) or suppressed exophers (*ftt-2*) (even without auxin-i.e., tagging induced the loss of function phenotype which prevented use for inducible tissue specific disruption).

See **Supplementary Table 1** for strain list.

### Plasmid constructions

New allele information is described in **Supplementary Table 1.** We created single-copy integration using MiniMOS technology and pCFJ1662 hygromycin or pCFJ910 G418-resistance conferring backbones^87^. For extra-chromosomal array transgenesis and selection we used hygromycin or G418 selection by exposing animals to selection plates. To make selection plates, we used 5mg/mL for hygromycin or 32mg/mL for G418 and added 200ul per 6cm OP50-seeded plate.

See **Supplementary Table 1** for RNAi bacteria clone sequences.

### Human homolog optimization and expression

Both IFD-1 and IFD-2 are most closely related to the named mammalian (taxid : 9606) neurofilament light polypeptide (<u>NP_006149.2</u>) with e-values of 2e-15 and 3e-13 for IFD-1 and IFD-2 respectively, sharing 24% identity. To optimize human genes for *C. elegans* expression based on codon and intron bias and *C. elegans* amino acid to tRNA ratio, we used *C. elegans* Codon Adapter GGA Software^59^. Nucleotide coding regions of mammalian proteins are provided. NCBI Reference Sequence: NM_006158.5 for *homo sapiens* neurofilament light. Optimized hNFL sequence available in Supplementary Table 1.

### CRISPR-CAS9 genomic *ifd-2* deletion

*ifd-2(bz187)* is a deletion of base pairs 816271-815275. The first three exons are affected by the genomic deletion which eliminates 996 base pairs, including the *ifd-2* initiation codon. Guide RNA and repair oligo available in Supplementary Table 1.

### RNAi

We achieved gene inactivation by feeding animals RNAi bacteria (HTT115) from L4-Ad2 or for 2 generations as noted, using standard methods. We performed RNAi experiments in strains in which neurons are RNAi-enhanced via SID-1 expression^7, 8^ required for effective feeding RNAi effects in neurons^55^. We used two transgenes for neuronal RNAi enhancement, initially working with the P*_mec-18_sid-1* transgene, which enhances touch neuron RNAi. RNAi studies in our group later established that the pan-neuronal P*_rgef-1_sid-1* constructs reliably enabled stronger RNAi knockdown of most touch neuron genes we targeted, and thus we focused more on use of the pan-neuronal knockdown strain. As noted in the strain list, some strains expressed pan-neuronal *sid-1* in the background of a *sid-1* null mutant, such that only transcripts in neurons could be targeted for degradation. Whenever possible, we confirmed RNAi knockdown effects using alternative approaches.

### Age synchronization

To synchronize animals, we selected L4 stage hermaphrodites and transferred them to test plates. The day after moving was considered adult day 1, Ad1. We scored animals on adult day 2 (Ad2) unless otherwise noted. For scoring exophers, we measured animals on plates using a Kramer pseudo-stereo dissecting scope (20x objective) and immobilized using 100ul of 10mM tetramisole; exophers were also visible in live animals without anesthetic. We scored ALMR neurons for exopher events in a binary (yes/no) manner.

### Exopher features for scoring

Exophers are readily visible at 400X total magnification with high power dissecting microscopes. We performed exopher characterization methods as described in detail.^8^ We note that baseline exopher levels in the P*_mec-4_*mCherry strain is generally in the range of 5-15% ALMR exophers on Ad2. The general or overall (mean) variability of other genetic backgrounds can differ. It is critical to *always* compare control and experiment on the same day because of the baseline variability—a test outcome must be compared to the background of its paired control.

### Microscopy

Most high resolution fluorescence micrograph data were captured on Zeiss Axiovert Z1 microscope equipped with X-Light V2 Spinning Disk Confocal Unit (CrestOptics), 7-line LDI Laser Launch (89 North), Photometrics Prime 95B Scientific CMOS camera, using Metamorph 7.7 software. A 100x oil immersion objective, 63x oil immersion, or 40x objective was used.

### Fluorescent colocalization analysis

We imaged touch neurons for colocalization experiments. Where needed, we used DIC confirmation to identify touch neurons of interest. For mNG::HttQ74 and HttQ128::CFP colocalization, we used only neurons in which visible mNG/CFP-aggregation was observable in the cell (as opposed to dim cytosolic mNG signal or absent CFP signal) for colocalization calculations. We analyzed thresholded multi-color image channels using ImageJ FIJI with JACoP and Coloc2 analysis software to calculate Pearson’s and Mander’s overlap coefficients.^88^

We used Pearson’s coefficient to quantitate colocalization of organelle tags and calculated Mander’s overlap of thresholded double-channel images by using the channel relevant to tag IFD over the channel of the other protein tag. We manually analyzed images in each 0.2 um Z-stack for colocalization often observing Z-stacks laterally in 3D projection for 3-dimensional observation.

### Soma GFP::IFD-1 focus measurement

We imaged whole touch neurons using 0.2 um Z-stacks. We measured the largest diameter of each inclusion by calculating the longest pixel length using ImageJ software. We omitted neurons with the presence of an exopher(s) from GFP::IFD-1 size calculations. We calculated the number of IFD inclusions using a standard threshold.

### Drug assays

For proteostasis drug treatments, we dissolved MG132 (Sigma-Aldrich C2211) and Spautin-1 (Sigma-Aldrich SML0440) in DMSO at 10mM concentration and administered by placing 40μL of each solution over the bacterial food lawn. We exposed animals to drug-plates from L4-Ad2. For DHC-1::degron AID-system treatments, we dissolved auxin in ethanol to 0.4M. We added 25μL 0.4M auxin and 175μl distilled water. We pipeted 200μl solution, per plate, on top of well-seeded OP50 plates. We exposed both control and experimental animals to auxin plates from L4-Ad2 - only the experimental animals harbored *bzSi6*[P*_mec-7_*TIR-1::TagBFP] necessary for auxin-inducible degradation.^34^

### Osmotic stress

We determined sensitivity to acute osmotic stress by moving L4 stage animals to NGM plates containing 450 mM concentrations of NaCl and scored exophers after an overnight exposure. We scored animals for exophers using ringed cytology slides on a fluorescent Kramer dissecting microscope.

### Oxidative stress experiments

We determined exopher sensitivity of animals to acute oxidative stress using paraquat (methyl viologen, Sigma). We transferred L4 animals to freshly prepared NGM plates containing paraquat 200 mM and assessed percent survival after an overnight exposure.

### Development assay

We measured post-embryonic development (PED) time by transferring eggs to an NGM plate. We performed a 50-gravid animal egg lay lasting one hour upon a new NGM plate. 60 hours after the egg lay, we checked plates every hour and the percentage of progeny to develop was assessed; we removed young adults after counting. We measured the time from hatching to the L4-adult transition, the time to development. For significance tests we used a standard two-tailed t-test comparing the mean value of three trials.

### Fertility

We measured brood size by placing individual L4 stage worms onto NGM plates. We transferred animals to new plates daily for three days, allowing progeny to develop to adulthood before paralysis and quantification.

### Lifespan

We measured lifespan was measured on NGM plates, transferring animals to fresh plates each day until day 5 of adulthood (Ad5) and after every four days thereafter. We assessed viability every two days as response to gentle prodding with a platinum pick. Animals that either had internal hatching of progeny or expulsion of internal organs were not counted as deaths.

### Mechanosensory touch function

We lightly stroked animals with a fine-hair pick behind the pharynx five times and scored a positive response as an animal movement in the reverse direction. For example, if an animal responded 2 out of 5 times, the animals has a 40% response rate.

### GST-4::GFP reporter assay

GST-4 is a glutathione S-transferase that facilitates the Phase II detoxification process. *gst-4* expression is regulated by *daf-16* and *skn-1 and* is a common reporter of cytosolic stress. We mounted animals expressing *dvIs19*[(pAF15)P*_gst-4_*GFP::NLS] +/- *ifd(Δ)* on microscope slides for 40x imaging and reported maximum intensity per animal in AU.

### Electron microscopy

We prepared animals for TEM analysis by high pressure freezing and freeze substitution (HPF/FS), and followed a standard to preserve ultrastructure. After HPF in a Baltec HPM-010, we exposed animals to 1% osmium tetroxide, 0.1% uranyl acetate in acetone with 2% water added, held at −90°C for 4 days before slowly warming back to −60°C, −30°C, and 0°C, over a 2 day period. We rinsed the samples several times in cold acetone and embedded the samples into a plastic resin before curing them at high temperatures for 1–2 days. We collected serial thin sections on plastic-coated slot grids and post-stained them with 2% uranyl acetate and then with 1:10 Reynold’s lead citrate, and examined with a JEOL JEM-1400 Plus electron microscope. By observing transverse sections for landmarks such as the 2nd bulb of the pharynx, it was possible to reach the vicinity of the ALM soma before collecting about 1,500 serial thin transverse sections.

We identified 29 ALM somata in serial sections from 23 blocks. We selected 12 candidate cells that were initially identified using light microscopy to contain touch neurons in the bud phase of exophergenesis. We imaged juxtanuclear regions of interest at high-magnification in serial sections, and we took high power electron tomograms (15K or 20K) in those neighborhoods, comparing regions showing either aggresome-like structures, mitochondria, or large lysosomes. We closely examined those regions for evidence of intermediate filaments surrounding and/or lying inside of the organelles. We used IMOD and TrakEM software for 3D analysis of the regions of interest, and for model-making.

### Statistical analysis

Each trial is graphed as a percentage of ALMR exopher occurrence (binary); we used Cochran–Mantel–Haenszel (CMH) analysis for P - value calculation of at least three or more biological trials. For IFD measurements, we used two-tailed t-tests. We analyzed the proteostress assays and comparison of number of IFD-inclusions by t-tests or one-way ANOVA (with Dunnet’s post-test) as noted.

### Blinding

For exopher measurements, we recorded strain information in a non-visible location. The data were unblinded following completion of the experiment. For blinding microscopy analysis, we mounted and imaged all animals on a slide for a given strain. Post-acquisition, the file names were assigned a random number and re-coded. The images were unblinded following analysis of the entire data set.

## Supplementary Data

**Supplementary Videos. Demonstration of the distinctive structural differences between membrane layers and intermediate filament types.**

P*_mec-4_*mCherry animals were staged as adult day 2 with an exopher bud phenotype.

**Video 1: Whorl membrane geometry.** Movie 1 shows a focus through series in which individual membranes continue through the full depth of the series without dropping out (unlike members of a filament bundle). Membranes often lie in closely packed layers but may bend sharply to double back on themselves. Scale bar is 50 nm.

**Video 2: Intermediate Filaments associated with the circular organelle.** Movie 2 shows filaments in a focus-through series demonstrates that (unlike membranes) these filaments do not continue through the full depth of the series, but individually disappear (or appear) at different depths within the series. Some longer filaments run for a distance around the organelle, bending gradually, but these filaments are loosely associated and rarely touch one another. Filaments deeper inside the organelle tend to be short, and run at odd angles to one another, rarely pairing. Scale bar is 50 nm.

**Supplementary Figure 1.**
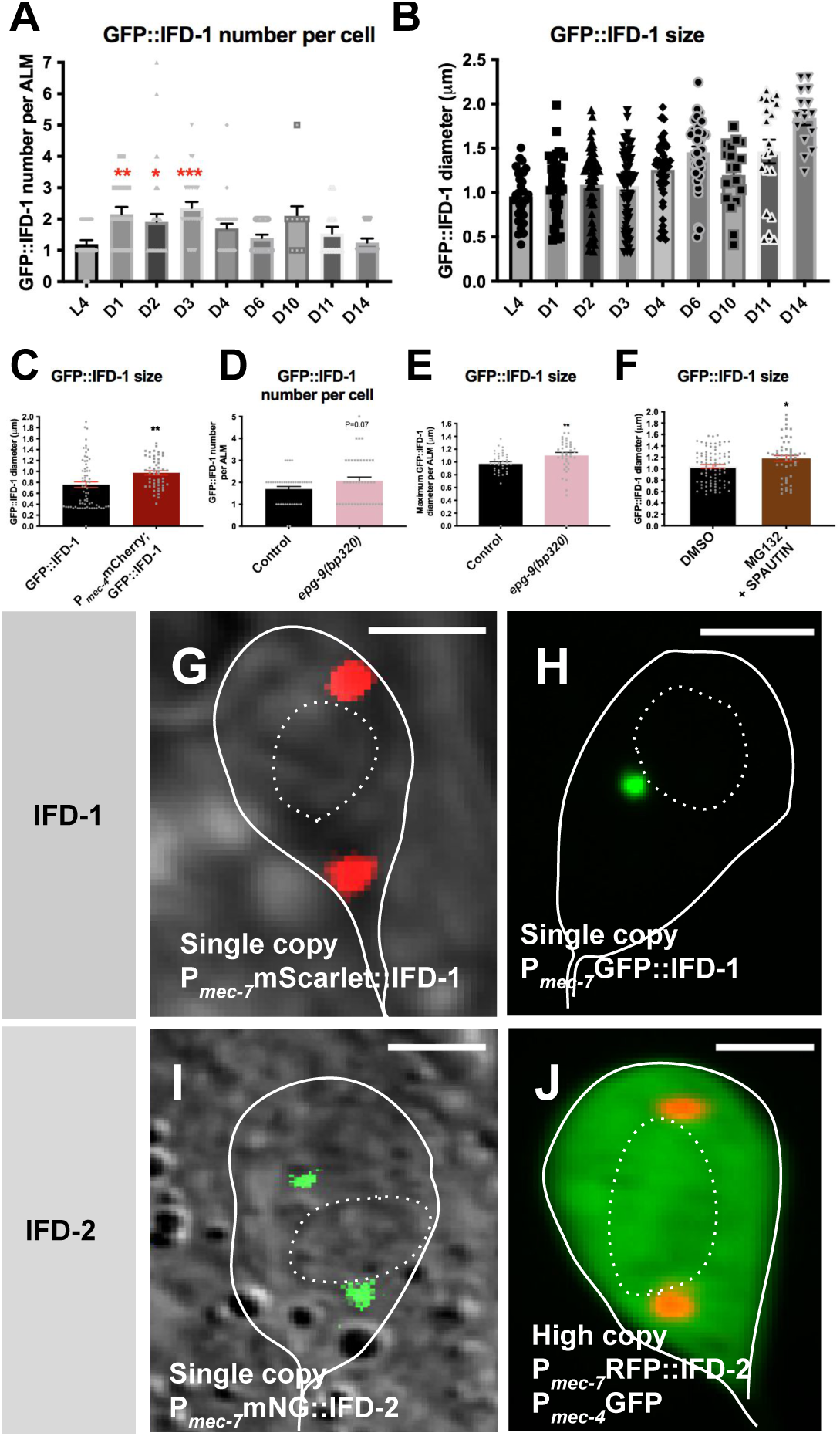
IFD-1 compartments are evident and enlarge under neuronal stress conditions. **A) Number of IFD-1 inclusions per ALM neuron with increasing age.** Strain: ZB4632 *bzIs166*[P*_mec-4_*mCherry]; *bzSi3*[P*_mec-7_*GFP::IFD-1]. Life-stage points assayed: L4, Ad1, Ad2, Ad3, Ad4, Ad6, Ad10, Ad11, Ad12, Ad14. N = 25, 19, 34, 28, 30, 25, 9, 13, 12 for lifestages respectively. One-way ANOVA, Dunnett’s post-test, error bars are SEM. *P < 0.05, *. P < 0.005, **. P < 0.0005, ***. **B) IFD-1-positive inclusions increase in size with age.** GFP::IFD-1 diameter in ALM neurons in µm. Strain: ZB4632 *bzIs166*[P*_mec-4_*mCherry];*bzSi3*[P*_mec-7_*GFP::IFD-1]. Life-stage points assayed: L4, Ad1, Ad2, Ad3, Ad4, Ad6, Ad10, Ad11, Ad12, Ad14. N = 25, 19, 34, 28, 30, 25, 9, 13, 12 for life stages, respectively. Error bars are SEM. **C) IFD-1-positive puncta increase in size under enhanced proteostress induced by mCherry expression.** GFP::IFD-1 diameter in ALM neurons on Ad2, in µm. Strain ZB4998 *bzSi3*[P*_mec-7_*GFP::IFD-1] and ZB4632 *bzIs166*[P*_mec-4_*mCherry]; *bzSi3*[P*_mec-7_*GFP::IFD-1]**, \*\***P < 0.005, N > 50/strain, 3 trials, two-tailed t-test, error bars are SEM. **D) IFD-1-positive puncta number per cell increases in the *epg-9/*ATG101 mutant.** Strain: ZB4993 *epg-9(bp320)*III; *bzIs166*[P*_mec-4_*mCherry]; *bzSi3*[P*_mec-7_*GFP::IFD-1] and ZB4632 *bzIs166*[P*_mec-4_*mCherry]; *bzSi3*[P*_mec-7_*GFP::IFD-1]. P = 0.07, N > 30/strain, 3 trials, two-tailed t-test, error bars are SEM. *P < 0.05, *. P < 0.005, **. P < 0.0005, ***. **E) Maximum IFD-1-positive puncta increase in size in *epg-9/*ATG101 mutant.** Strain: ZB4993 *epg-9(bp320)III*; *bzIs166*[P*_mec-4_*mCherry]; *bzSi3*[P*_mec-7_*GFP::IFD-1] and ZB4632 *bzIs166*[P*_mec-4_*mCherry]; *bzSi3*[P*_mec-7_*GFP::IFD-1]. Maximum GFP::IFD-1 punctae diameter per ALM, in µm. **P < 0.005, N > 30/strain, 3 trials, two-tailed t-test, error bars are SEM. **F) IFD-1-positive puncta increase in size under enhanced proteostress induced by proteasome and autophagy inhibitors.** We treated strain ZB4632 *bzIs166*[P*_mec-4_*mCherry]; *bzSi3*[P*_mec-7_*GFP::IFD-1] with both 10 mM proteasome inhibitor MG132 and 10 mM autophagy inhibitor Spautin-1, proteostressors that are known to enhance exopher production, from L4 to Ad3 and measured GFP::IFD-1 inclusion size at Ad3. GFP::IFD-1 diameter in ALM neurons in µm. P < 0.05, N > 50/strain, 3 trials, two-tailed t-test, error bars are SEM. **G-J. Multiple IFD-1 and IFD-2 fluorescently-tagged reporter proteins (tags GFP, RFP, mScarlet, mNeonGreen) exhibit similar localization to juxtanuclear foci.** Adult ALM soma outlined with continuous line, nucleus outlined with dotted-line. Scale bar = 2 µm. Images representative of N > 10. Note that G-I are single copy, J is from a high copy transgene array. **G)** ZB5021 *bzSi34*[P*_mec-7_*mSc::IFD-1] **H)** ZB4998 *bzSi3*[P*_mec-7_*GFP::IFD-1] **I)** ZB5091 *bzSi37*[P*_mec-7_*mNG::IFD-2]; *bzIs166*[P*_mec-4_*mCherry] **J)** ZB4866 *bzEx253*[P*_mec-7_*RFP::IFD-2]; *zdIs5*[P*_mec-4_*GFP]. Free touch neuron GFP diffuses throughout the cytoplasm. Note: Fluorescently tagged IFD-1 and IFD-2 proteins confer functional complementation of exophergenesis defects and are therefore likely to localize to, and indicate the identity of, the native subcellular compartments in which they are bioactive.

**Supplementary Figure 2.**
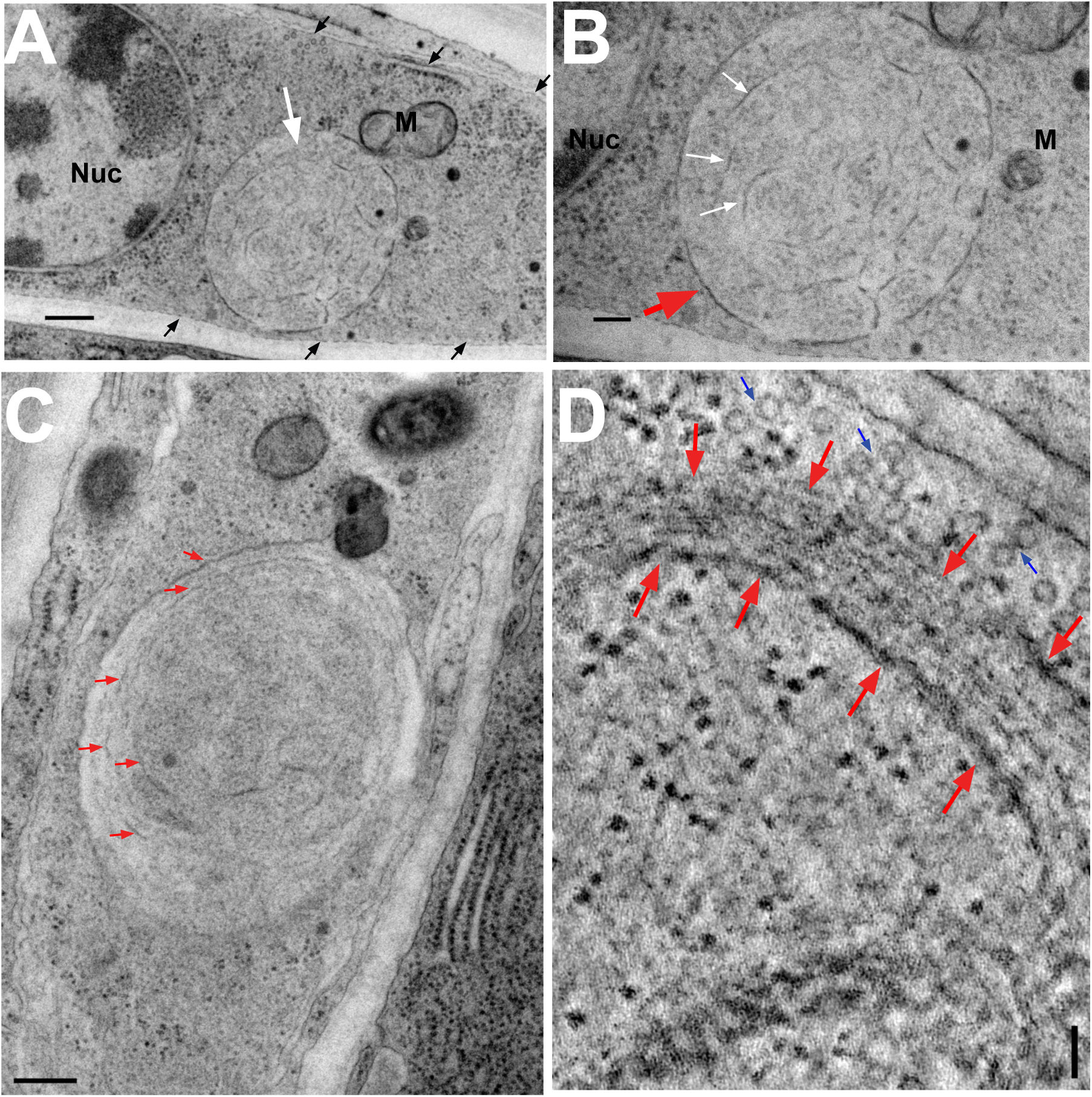
Ultrastructural analysis reveals the presence of intermediate filament assemblies in distinctive perinuclear structures in proteo-stressed ALM neurons. We selected P*_mec-4_*mCherry animals at Ad2 with touch neurons that had early exopher buds that had not separated from the neuron of origin (12 ALMs examined in EM). We identified touch neuron somata on the basis of well characterized ultrastructural features after extensive transverse thin section series. ALM neuron somata (L/R pair) lay on opposite sides of the midbody, at similar distances from the head and can be unambiguously identified via their characteristic microtubule-filled dendrite features. **A)** Lower power TEM view of an ALM neuron in transverse section, embedded in the hypodermis next to the cuticle. A distinctive circular structure (white arrow) lies close to the nucleus (Nuc) and a small mitochondrion (M). Electron lucent contents are dominated by the presence of single filaments. No ribosomes or other small organelles are detectable here, which would be expected if the contents had been derived from typical cytoplasm. Shorter filaments ∼ 10 nm in width lie at random angles within the circular organelle, while a single layer of discontinuous filaments of similar width coats the periphery of the organelle. Filament measures are consistent with their identification as intermediate filaments (9.98 nm average; > 150 filaments measured). A bundle of microtubules lies just beneath the cuticle layer (cuticle annotated with black arrowheads). Scale bar 500 nm. **B)** Higher magnification view of the circular organelle from panel A, showing the filaments in higher resolution (white arrows). Red arrow indicates one of the longer filaments taking position at the outer edge of the organelle. Filaments sometimes adopt a zig-zag pattern of bent filaments as they bundle. Note the thinness of IFs compared to the membrane bilayer surrounding the nearby mitochondrion. Demonstration of how intermediate filaments can be visually distinguished from membrane are included in Supplementary Video 1 and Supplementary Video 2. Scale bar 200 nm. **C)** Closeup view of a similar rounded organelle (soma of a different animal) in which most intermediate filaments (red arrows) have assembled into a loose bundle that curves as the bundle envelopes most of the periphery. A small number of shorter filaments lie deeper inside the organelle. Scale bar is 200 nm. **D)** Thinner view from an electron tomogram shows curving individual filaments bordering the circular structure, but with a denser, tighter accumulation of filaments (between red arrows), at the edge of the organelle. Note also the bundle of microtubules (blue arrows) lying just outside the indicated organelle. Some very dark ribosomes are now seen within the structure, as well as granular matrix material of moderate electron density that likely represents proteinaceous aggregates. Scale bar is 200 nm. **Discussion**. Although successive sequencing is not possible for individual samples, we speculate that the juxtanuclear structures might mature by increasing and expanding the proteinaceous center and adding more intermediate filaments to the periphery. Ultrastructure of the mammalian aggresome has been reported in cultured HEK cells, with aggresomes described as electron dense cores of (CFTR)-aggregated protein deposits devoid of membrane surroundings, deposited likely intimate to the nuclear membrane, and surrounded by 8 - 10 nm sized filaments that form loosely parallel bundles.^4^ Distorted ‘tangles’ of IFs within the dense aggresome core have been noted; organelles can be positioned nearby the aggresome similar to our EM observations (mitochondria in Supplementary Figure 2A) and microscopy observations (Figure 1G). Overall, limited low resolution EM data exist on mammalian aggresome-focused EM.^4, 6, 27, 28, 63–67^ Although the initial description describing IF “cages” was based on fluorescent microscopy,^4^ available EM data suggest at best loose and somewhat disorganized IF networks in aggresome structure, similar to what we report here.

**Supplementary Figure 3.**
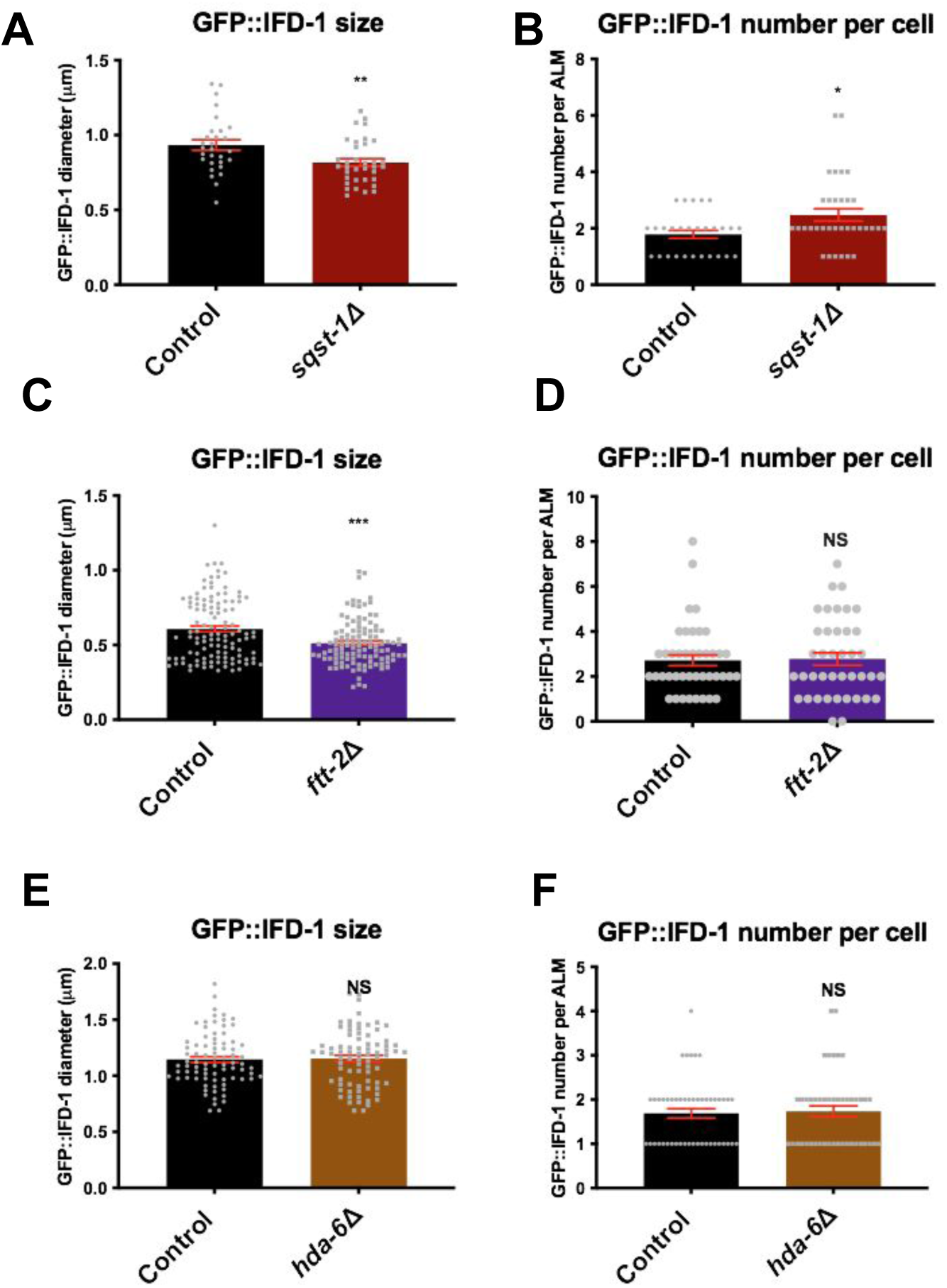
Disruption of *hda-6*, *sqst-1,* and *ftt-2/*14-3-3 can modulate IFD compartment size but no single disruption eliminates the aggresome-like compartment. **A) *hda-6(ok3203)* does not affect average IFD-1 puncta size in *bzIs166*[P*_mec-4_*mCherry] background.** Strains are ZB4801 *bzIs166*[P*_mec-4_*mCherry]; *hda-6(ok3203), bzSi3*[P*_mec-7_*GFP::IFD-1] and ZB4632 *bzIs166*[P*_mec-4_*mCherry]; *bzSi3*[P*_mec-7_*GFP::IFD-1]. Diameter of IFD-1 puncta measured in ALM on Ad2 in µm. N > 60, 4 trials, two-tailed t-test, error bars are SEM. **B) *hda-6(ok3203)* does not significantly affect IFD-1 puncta number per cell in the *bzIs166*[P*_mec-4_*mCherry] background.** Strain is ZB4801 *bzIs166*[P*_mec-4_*mCherry]; *hda-6(ok3203); bzSi3*[P*_mec-7_*GFP::IFD-1] and ZB4632 *bzIs166*[P*_mec-4_*mCherry]; *bzSi3*[P*_mec-7_*GFP::IFD-1]. HDA-6 is an ortholog of human HDAC6, a cytoplasmic deacetylase with conserved function as an adapter, microtubule modulator, and autophagy regulator. Number of IFD-1 puncta per ALM Ad2, N > 60, 4 trials, two-tailed t-test, NS, error bars are SEM. **C) Maximum IFD-1 puncta size decreases in *sqst-1*(*ok2892*).** *sqst-1* encodes the ortholog of human SQSM1, an aggregate adapter protein implicated in mammalian aggresome formation and autophagy. Strains are ZB4632 *bzIs166*[P*_mec-4_*mCherry]; *bzSi3*[P*_mec-7_*GFP::IFD-1] and ZB4838 *bzIs166*[P*_mec-4_*mCherry]; *sqst-1ok2892*) IV; *bzSi3*[P*_mec-7_*GFP::IFD-1]. Diameter of the largest IFD-1 puncta in ALM Ad2 in µm. P < 0.05, N > 25, 3 trials, two-tailed t-test, error bars are SEM. **D) IFD-1 puncta number per ALM increases in *sqst-1*(*ok2892*).** Strains are ZB4632 *bzIs166*[P*_mec-4_*mCherry];*bzSi3*[P*_mec-7_*GFP::IFD-1] and ZB4838 *bzIs166*[P*_mec-4_*mCherry]; *sqst-1(ok2892)IV*; *bzSi3*[P_mec-7_GFP::IFD-1]. Number of IFD-1 puncta per Ad2 ALM indicated. P < 0.005, N > 25, 3 trials, two-tailed t-test, error bars are SEM. **E) Maximum IFD-1-positive puncta size decreases in *ftt-2(n4426).*** Diameter, in µm, of the largest Ad1 IFD-1-positive puncta per ALM neuron decreases in the *ftt-2* predicted null background, in a strain expressing *bzIs166*[P*_mec-4_* mCherry]; *bzSi3*[P*_mec-7_* GFP::IFD-1]. P < 0.0001, N > 50, 3 trials, two-tailed t-test, error bars are SEM. We measured on Ad1 to reduce animal-bagging consequences. **F) IFD-1-positive puncta number per ALM neuron remains the same in control and *ftt-2(n4426).*** Number of IFD-1-positive inclusions in Ad1 ALM somas does not change in mutant *ftt-2(n4426)* versus WT in strains expressing *bzIs166*[P*_mec-4_* mCherry]; *bzSi3*[P*_mec-7_* GFP::IFD-1]. NS, P = 0.87, N > 50 / strain, Ad1, 3 trials, two-tailed t-test, error bars are SEM. We measured on Ad1 to reduce animal-bagging consequences.

**Supplementary Figure 4.**
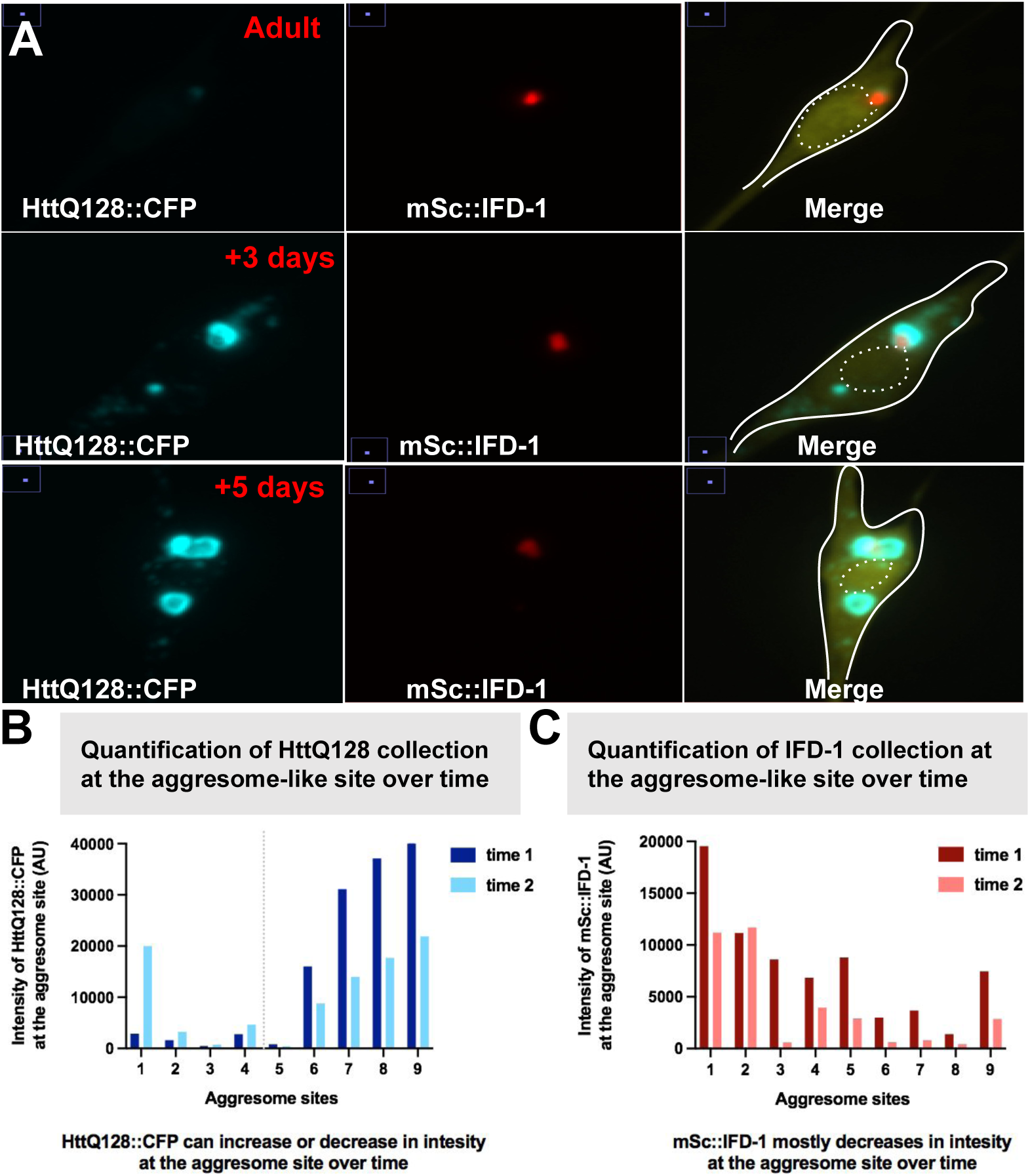
Disease-associated human Htt-polyglutamine expansion protein dynamically colocalizes with IFD-1 in touch neurons. **A) HttQ128::CFP and mSc::IFD-1 compaction over time.** Images are selected from a 5-day time-course of an ALM neuron expressing HttQ128::CFP and mSc::IFD-1. Neurons were imaged as an adult, rescued from microscope slides and placed back on plates. Three days later, the same neuron was imaged. Two more days later (+5 days) the neuron was imaged again. Strain ZB5104 *igIs1*[P*_mec-7_* YFP, P*_mec-3_* Htt57Q128::CFP, lin-15(+)]; *bzSi34* [P*_mec-7_* mScarlet::IFD-1]. Out of nine multi-day observations of when mSc and CFP interact, we find about half the time (4/9 observations), HttQ128 can globularly expand at the mSc::IFD-1-aggresome-like site. **B) Quantification of HttQ128 collection at the aggresome-like site over time.** We imaged touch neurons that co-expressed interacting HttQ128::CFP and mSc::IFD-1 at time 1 (nine animals distributed over the X-axis), then rescued the animals from microscope slides and placed animals back on typical OP50-plates for at least 2 days. We then imaged the same neurons again, at least 2 days later (time 2). Following the same aggresome-like site over a period of at least 2 days, we find that HttQ128::CFP can exhibit a dynamic collection pattern. HttQ128 can either globularly expand at the aggresome-like site (as measured by intensity, AU) in 4/9 neurons (and as depicted in Panel A), or decrease in intensity at the aggresome site, observed in 5/9 neurons. **C) Quantification of mSc::IFD-1 collection at the HttQ128-filled aggresome-like organelles.** In the same experiment as in Panel B, we scored the same aggresome-like site over a period of at least two days, we found that mSc::IFD-1 typically displays a relative reduction in collection over time, as measured by maximum intensity (AU) at the aggresome-like site (8/9 neurons displayed reduced intensity at the aggresome-like site). Interestingly, aggresome dynamics in the absence of aggresome-targeted protein HttQ128 appear different (only GFP::IFD-1 observed over time; see Supplementary Figure 1B which shows GFP::IFD-1 aggresome-like growth over time in absence of aggresome-targeting-protein, HttQ128). Our data suggest alternative dynamics of the IF-aggresome-like organelle in the presence of an elevated aggresome-targeted-aggregate like HttQ128. Note that the X-axis is plotted to show trend of growth between timepoint 1 and timepoint 2 and aggresome-like sites cannot be compared between panels B and C.

**Supplementary Figure 5.**
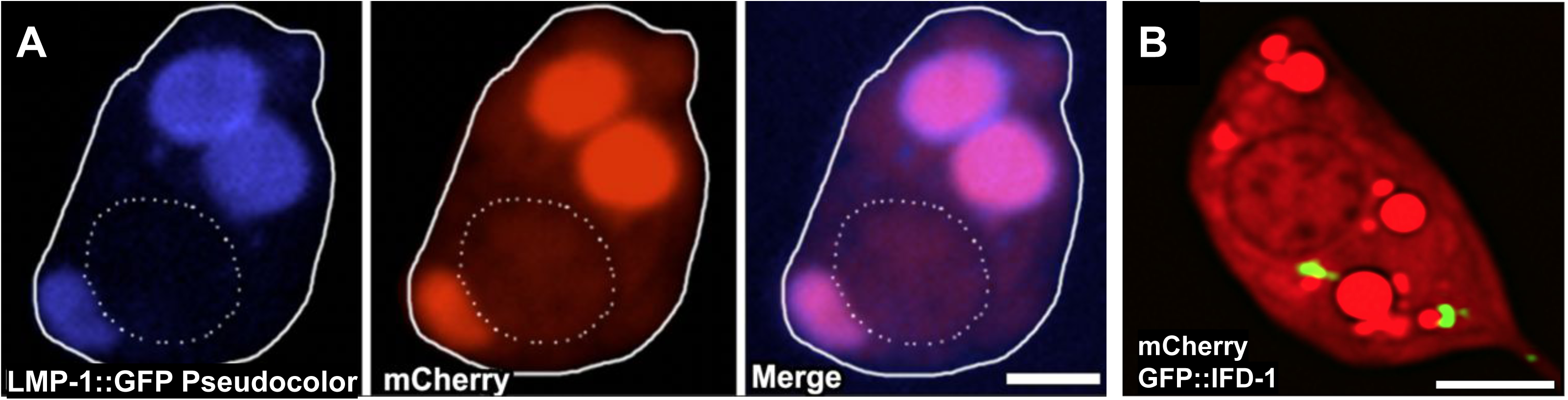
Over-expressed mCherry is predominantly housed in LMP-1-bound lysosomal-organelles, which are distinct from the IF-compartment. **A) mCherry is mostly in LMP-1-bound structures.** A representative adult ALM neuron co-expressing mCherry with the LMP-1::GFP lysosome marker (white solid line outlines the soma cell body and the dotted line outlines the nucleus). Representative of N > 250. LMP-1::GFP signal is enriched in the lysosome membrane forming a GFP ring (pseudo-colored blue) around the mCherry collection. mCherry degradation appears minimal within the LMP-1::GFP compartment as signal persists. GFP channel is pseudo-colored to blue to emphasize reporter difference between panel A and B. Strain ZB4509 *bzIs166[*P*_mec-4_*mCherry]*; bzIs168*[P*_mec-7_*LMP-1::GFP]. Scale bar = 2 µm. **B) IFD-positive organelles are often localized next to, but not overlapping with, a subset of mCherry concentrations when tagged IFD is expressed from a single copy transgene**. Representative Ad2 touch neuron, representative of N > 50. Strain ZB4632 *bzIs166[*P*_mec-4_*mCherry]*; bzSi3*[P*_mec-7_*GFP::IFD-1]. Scale bar = 2 µm

**Supplementary Figure 6.**
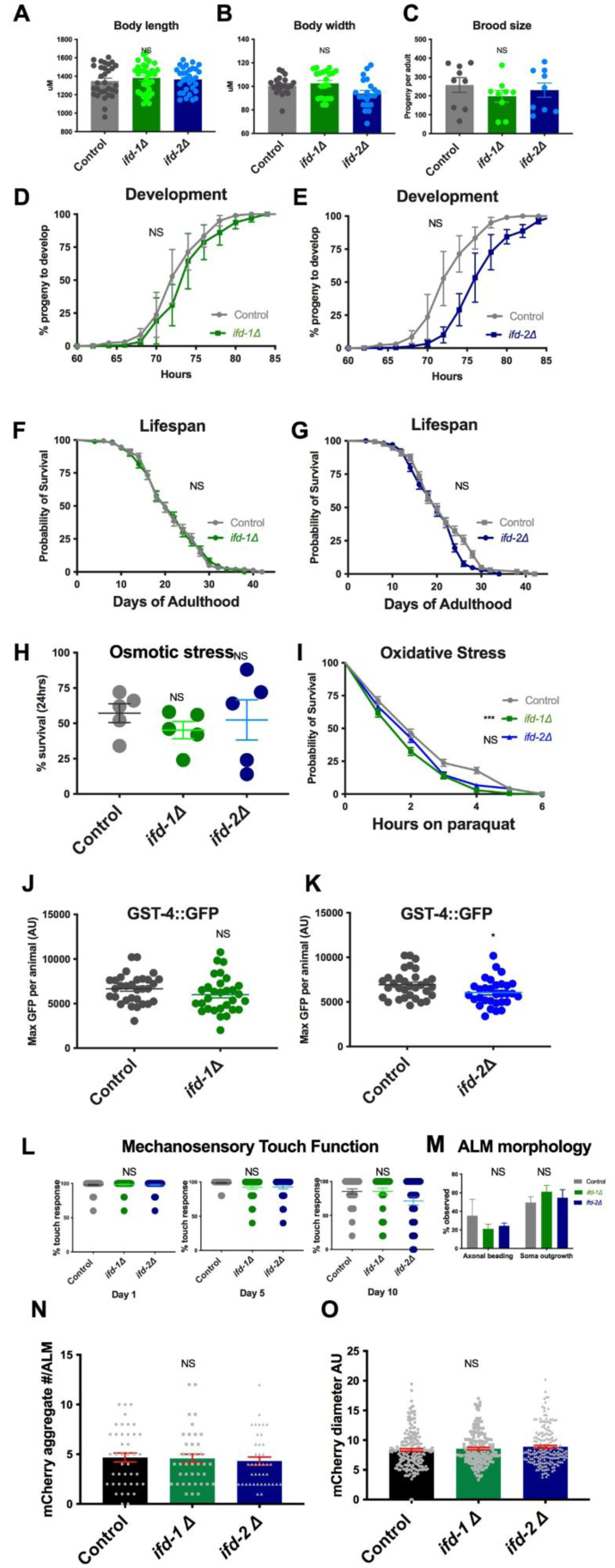
General health, development, lifespan, stress response measures and touch neuron integrity and function are largely preserved in the *ifd* null mutants. Strains were ZB4065 *bzIs166*[P*_mec-4_*mCherry], ZB4722 *ifd-1(ok2404);bzIs166*[P*_mec-4_*mCherry], and ZB4655 *ifd-2(bz187); bzIs166*[P*_mec-4_*mCherry]. **A) Body length** in µM as measured from 10x images taken from dissection microscope. Synchronized Ad2 animals. N > 30, NS, 3 trials, One-way ANOVA. **B) Body width** as measured from 10x images taken from dissection microscope. Synchronized Ad2 animals. N > 30, NS, 3 trials, One-way ANOVA. **C) Brood size**. Progeny number per adult. We placed larval animals onto individual plates. Progeny of these animals were counted during their entire life span and is given as an average number per individual. N = 9, P = NS, 3 trials, One-way ANOVA. **D) Development for *ifd-1(ok2404)*.** Hours from egg to develop into young adult stage. Isolated embryos (synchronized by an hour egg lay using 50 animals) were placed onto each plate. We monitored Individual larvae every day to identify L4 stage animals. We determined time of development by calculating the average number of days animals needed to reach adulthood. N > 150, P = 0.72, 3 trials. We calculated statistical significance using an unpaired two-tailed t-test where mean values of 3 trials were considered using GraphPad Prism. **E) Development for *ifd-2(bz187)*.** We determined hours from egg into young adult stage. We placed isolated embryos (synchronized by an hour egg lay using 50 animals) onto each plate and monitored individual larvae every day for L4 stage markers. We calculated time of development as the average number of days animals needed to reach adulthood. N > 150, P = 0.38, 3 trials. We calculated statistical significance using an unpaired two-tailed t-test in which mean time to development values of 3 trials were compared using GraphPad Prism ^94^. **F) Survival curve.** *ifd-1(ok2404); bzIs166*[P*_mec-_*4mCherry]. **G) Survival curve.** *ifd-2(bz187); bzIs166*[P*_mec-_*4mCherry]. Graphed as days until death. We scored newly hatched animals for viability every day using a platinum wire for mechanical stimulation; movement response was scored as alive. We transferred animals at least every two days to prevent mix-up with their hatched progeny. We excluded animals that could not be located on the plate from statistical analysis. N > 150, *ifd-1* P = 0.9442; *ifd-2* P = 0.1152, 3 trials, We performed statistical analysis using the survival function and the Gehan-Breslow-Wilcoxon Test of GraphPad Prism ^94^. **H) Osmotic stress.** Percent survival after 24 hours of exposure to 450 mM NaCl concentrated plates; strains *ifd-1(ok2404);bzIs166*[P*_mec-_*4mCherry] and *ifd-2(bz187); bzIs166*[P*_mec-_*4mCherry]. Osmotic stress is associated with proteostasis disruption ^47^. For osmotic stress we used NGM agar with 450 mM NaCl. We stored plates overnight at room temperature and subsequently inoculated with concentrated OP50 overnight culture; after overnight incubation at room temperature plates were ready for assay. We started osmotic stress experiments by transferring L4 larvae onto the bacterial lawn followed by an overnight incubation. Subsequently, we washed in recovery buffer (M9 buffer with 150 mM NaCl) and transferred to normal NGM plates. The viability of each animal was scored after an additional overnight incubation using mechanical stimulation. N > 150, *ifd-1* P = 0.329, *ifd-2* P = 0.76, 3 trials; we used a two-tailed t-test for significance calculation ^94^. **I) Oxidative stress.** Hours on paraquat are graphed vs. survival; strains *ifd-1(ok2404);bzIs166*[P*_mec-_*4mCherry] and *ifd-2(bz187); bzIs166*[P*_mec-_*4mCherry]. Oxidative stress is associated with proteostasis disruption ^48^. For oxidative stress we used 200 mM paraquat in NGM agar. We stored plates overnight at room temperature and subsequently inoculated with concentrated OP50 overnight culture. Following overnight incubation at room temperature plates were ready to use. We started oxidative stress experiments by transferring L4 larvae onto the bacterial lawn. We scored animals hourly for viability using mechanical stimulation with a platinum wire. Control plates did not contain paraquat. N > 150*, ifd-1* ***P = 0.0004, *ifd-2* P = 0.076, 3 trials. We performed statistical analysis using the survival function and the Gehan-Breslow-Wilcoxon Test of GraphPad Prism ^94^. **J) GST-4 expression in *ifd-1(ok2404).*** GST-4 is a glutathione S-transferase that facilitates the Phase II detoxification process. GST-4 expression is regulated by *daf-16* and *skn-1* and GST-4::GFP is a common reporter of cytosolic stress. Maximum intensity per animal reported in AU. GST-4::GFP expression is NS (P = 0.16) in *ifd-1* mutants. N > 30, 3 trials. Two-tailed t-test. Strains: *dvIs19*[(pAF15)P*_gst-4_*GFP::NLS] and *dvIs19*[(pAF15) P*_gst-4_*GFP::NLS]; *ifd-1(ok2404)*. **K) GST-4 expression in *ifd-2(bz187).*** Maximum intensity per animal reported in AU. GST-4::GFP expression is reduced in *ifd-2* animals (P = 0.0256). N > 30, 3 trials. Two-tailed t-test. Strains: dvIs19[(pAF15)P*_gst-4_*GFP::NLS] and dvIs19[(pAF15)P*_gst-4_*GFP::NLS]; *ifd-2(bz187)*. Note that increasing proteostress by disruption of proteosome or autophagy generally increases exopher production ^7^ – this is the opposite effect of the loss of *ifds,* which suppresses exopher formation, arguing against the *ifd* deletions as generators of systemic stress that influence exophers. **L) Mechanosensory touch function** on Ad1, Ad5, and Ad10 for *ifd-1(ok2404);bzIs166*[P*_mec-_*4mCherry] and *ifd-2(bz187);bzIs166*[P*_mec-_*4mCherry]. We lightly stroked animals with a fine-hair pick behind the pharynx five times and recorded response (direction reversal). For example, if an animal responded 2 out of 5 times, the animals had a 40% response rate. NS, One-way ANOVA, Ad1 - 3 trials, N > 90, Ad5 - 1 trial N > 30, Ad10 - 3 trials, N > 90. **M) ALM morphology** of strains *ifd-1(ok2404);bzIs166*[P*_mec-_*4mCherry] and *ifd-2(bz187);bzIs166*[P*_mec-_*4mCherry]. We scored axonal beading and soma outgrowth percentage on Ad2. Beading and outgrowths are associated with age, stress, and neurodegeneration.^95^ NS, One-way ANOVA N > 30, 3 trials. **N) *ifd-2* mutants display normal number of mCherry collections per ALM.** Animals expressing *bzIs166*[P*_mec-4_*mCherry] with the *ifd-1* or *ifd-2* mutation have the same number of mCherry aggregates as wildtype – 4.67, 4.56, and 4.31 mean mCherry collections per cell respectively for control, *ifd-1*, and *ifd-2*. N = 42, 39, 41 respectively, one-way ANOVA with Dunnett’s post-test, 3 trials, P = 0.98 for *ifd-1,* P = 0.79 for *ifd-2*. **O) *ifd-2* mutants display normal size of mCherry collections per ALM.** WT mCherry collections per ALM expressing *bzIs166*[P*_mec-4_*mCherry] measures a mean puncta diameter of 8.34 AU; there is a NS change in the size of mCherry collections in the *ifd-1* mutant (8.57 AU mean diameter) or *ifd-2* mutant (8.88 AU mean diameter). N = 42, 39, 41 respectively, one-way ANOVA with Dunnett’s post-test, 3 trials, P = 0.69 for *ifd-1,* P = 0.15 for *ifd-2*.

**Supplementary Figure 7.**
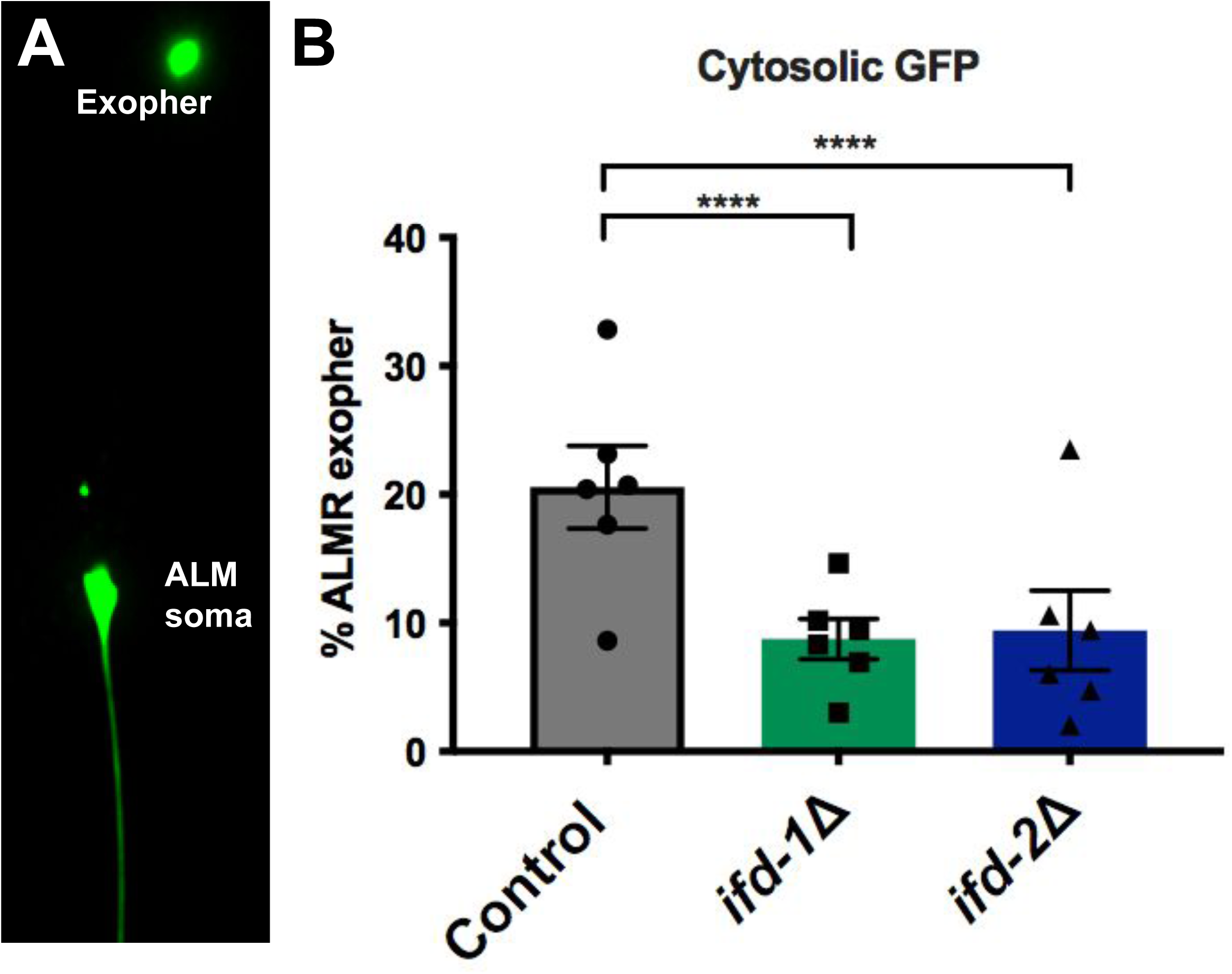
*ifd-1* and *ifd-2* deletions decrease ALMR exophers as reported by a highly expressed cytosolic GFP reporter. **A)** Representative Ad2 exopher produced from touch neurons in *uIs31*[P*_mec-17_*GFP]. **B)** We scored ALMR exopher production on Ad2 in the *ifd-1(ok2404)* deletion mutant and the *ifd-2(bz187)* deletion mutant, each of which also harbored *uIs31*[P*_mec-17_*GFP] integrated transgene. N > 331, 6 trials. *ifd-1*(P < 0.0005); *ifd-2* (P < 0.0005); * P < 0.05, ** P < 0.005, *** P < 0.0005. Data support that *ifd* disruptions modulate exopher levels independently of the fluorescent reporter used for exopher detection. Although GFP is not a physiological substrate, we note that exophers have been identified in amphid neurons by dye filling fluorescence labeling that does not involve transgene reporter introduction^7^.

**Supplementary Figure 8.**
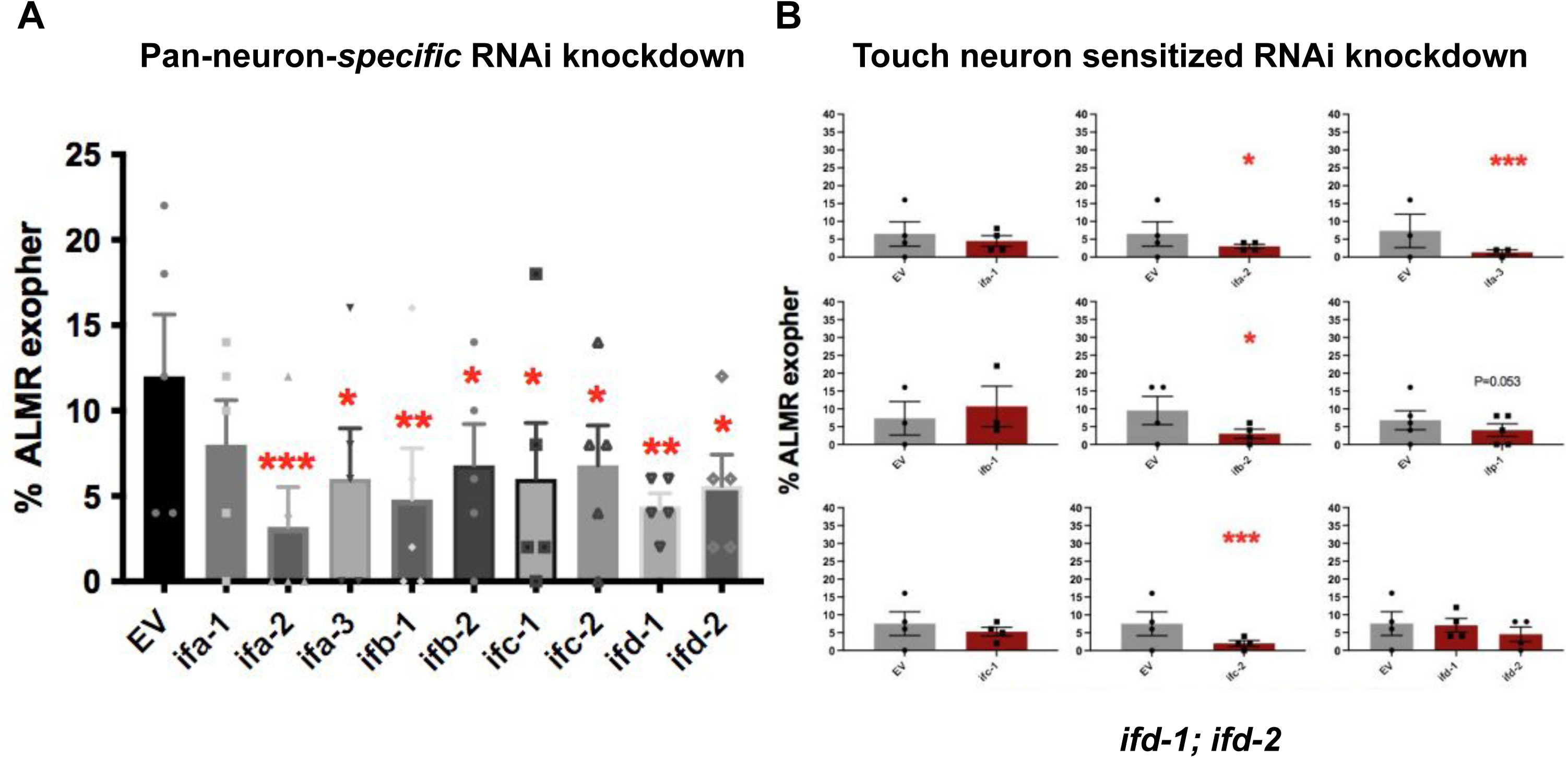
RNAi knockdown of intermediate filament genes can decrease exopher production. **A) Intermediate filament RNAi knockdown in a pan-neuron-specific RNAi strain decreases exopher percentage**. We performed RNAi knockdown of each of 9 *C. elegans* intermediate filament genes, exposing animals to RNAi clones for 2 generations, and assaying ALMR exophers on Ad2, screen strain ZB5184 *bzIs166* [P*_mec_-4*mCherry1] II; *sid-1(qt9)* V; *sqIs71* [P*rgef-1*GFP; P*rgef-1sid-1*;pBS]. Note that in this strain, RNAi should be effective only in all neurons, and not in other tissues; expression in touch neurons (as well as all other neurons) is anticipated to be targeted. Our control RNAi studies have found that use of the P*_rgef-1_* promoter to express dsRNA transporter SID-1 allows most potent neuronal RNAi targeting (gift of M. Hansen), and thus we use P*rgef-1sid-1* for initial RNAi rather than a touch neuron specific promoter strain, in which we find outcomes more variable. Sequence-confirmed RNAi clones were empty vector negative control (EV), *ifa-1*, *ifa-2*, *ifa-3*, *ifb-1*, *ifb-2*, *ifc-1*, *ifc-2*, *ifd-1*, and *ifd-2.* RNAi knockdown N = 250 ALMR/clone, 5 trials. * P < 0.05, ** P < 0.005, *** P < 0.0005. **B) RNAi knockdown of intermediate filament genes in the *ifd-1;ifd-1* double mutant can decrease exopher production.** Intermediate filament RNAi knockdown in the *ifd-1(Δ); ifd-2(Δ)* double mutant background can exacerbate decreases in exopher percentage to less than 5%. We performed RNAi knockdown of 10 *C. elegans* intermediate filament genes, exposing animals to RNAi clones from L4-Ad2, and assaying ALMR exophers on Ad2, screen strain ZB5247: *ifd-1(ok2404); ifd-2(bz187); bzIs166*[P*_mec-_*4mCherry], *uIs71*[(pCFJ90) P_myo*-2*_mCherry + P*_mec-18_*SID-1]. Because some exopher production is evident when *ifd-1* and *ifd-2* are both absent (Figure 4), we infer that a redundant activity or a parallel pathway must also contribute to exopher formation. Note that in this strain, all cells but neurons are amenable to RNAi, however transcripts in touch neurons should be targeted by RNAi as touch neurons express the neuronal dsRNA transporter *sid-1*; disruptions might not act touch neuron autonomously due to susceptibility of other cells. That is to say that although the above RNAi screen strain is touch-neuronally sensitized, without the *sid-1* mutation to disrupt targeting in other cells, we cannot rule out knockdown elsewhere, and therefore effect contribution, by other tissues. Sequence-confirmed RNAi clones were empty vector negative control (EV), *ifa-1, ifa-2, ifa-3, ifb-1, ifb-2, ifp-1, ifc-1, ifc-2*, and control clones - *ifd-1*, and *ifd-2.* Note that RNAi knockdown was performed in the *ifd-1(ok2404); ifd-2(bz187)* double mutant background, and as expected, RNAi targeting to either *ifd-1* or *ifd-2* does not change exopher levels compared to the EV control, suggesting that off-target effect of *if* knockdown are not potent in this study. RNAi knockdowns were 200 ALMR/clone, 4 trials. Note that some experimental RNAi clones were assayed on the same day and therefore used the same empty vector control. Each clone has been graphed separately with day-matched EV controls, for simplicity. Also note that very low baseline exopher levels are present for the *ifd-1(Δ); ifd-2(Δ)* starting strain so that any role implied by these RNAi data requires confirmation by independent genetic perturbation and cell autonomy analysis to confirm. Candidates for further study highlighted by this screen are: *ifa-2, ifa-3, ifb-2, ifc-2,* and potentially *ifp-1.* * P < 0.05, ** P < 0.005, *** P < 0.0005.

**Supplementary Figure 9.**
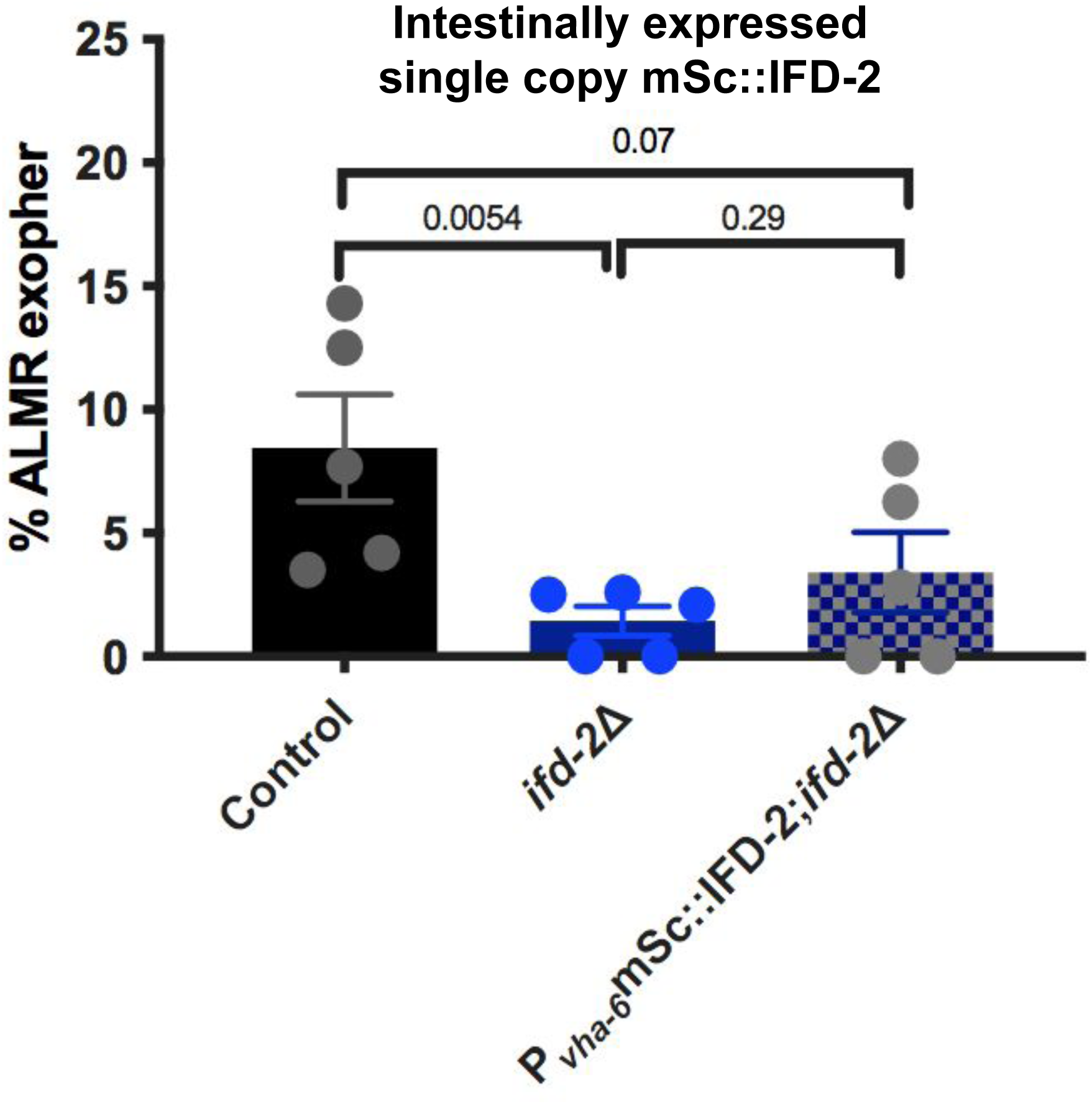
Intestine-specific *ifd-2* expression is not effective for significant rescue of *ifd-2(Δ)* defects in exopher production. *ifd-2* is expressed in intestine and can influence intestinal morphology^94^, raising the question as to whether the intestine might signal non-autonomously to impact exopher production. We therefore expressed *ifd-2* from the *vha-6* intestine-specific promoter in the *ifd-2(Δ)*; mCherry background and scored exopher levels. We compared ALMR exophers at Ad2 in *bzIs166*[P*_mec-4_*mCherry] compared to *bzIs166*[P*_mec-4_*mCherry]; *ifd-2(bz187)* (P=0.0054); *bzIs166*[P*_mec-4_*mCherry] compared to *bzIs166*[P*_mec-4_*mCherry]; *ifd-2(bz187); bzSi45*[P*_vha-6_*mSc::IFD-2] (P = 0.29). *bzIs166*[P*_mec-4_*mCherry]; *ifd-2(bz187)* compared to *bzIs166*[P*_mec-4_*mCherry]; *ifd-2(bz187); bzSi45*[P*_vha-6_*mSc::IFD-2] is P = 0.07, N > 130, 5 trials. Cochran-Mantel-Haenszel test, error bars are SEM. Comparing *ifd-2(bz187)* to *ifd-2(bz187);*P*_vha-6_*mNeonGreen::IFD-2, we find that intestinal expression of mNeonGreen::IFD-2 does not significantly rescue the neuronal exopher phenotype of *ifd-2(bz187),* consistent with a predominant role for *ifd-2* in exophergenesis in the neuron (Figure 4E-G); although the trend toward partial rescue leaves open the possibility of some intestinal contribution. Note: We could not detect expression from native in-line *ifd* promoters or native *ifd* transgenes in touch neurons (data not shown), a reporter outcome we often observe for native promoter-single copy reporters including touch neuron channel *mec-4*, dynein heavy chain *dhc-1*, and *rab-11,* a GTPase required for recycling endosome function, for examples ^34, 96^).

**Supplementary Figure 10.**
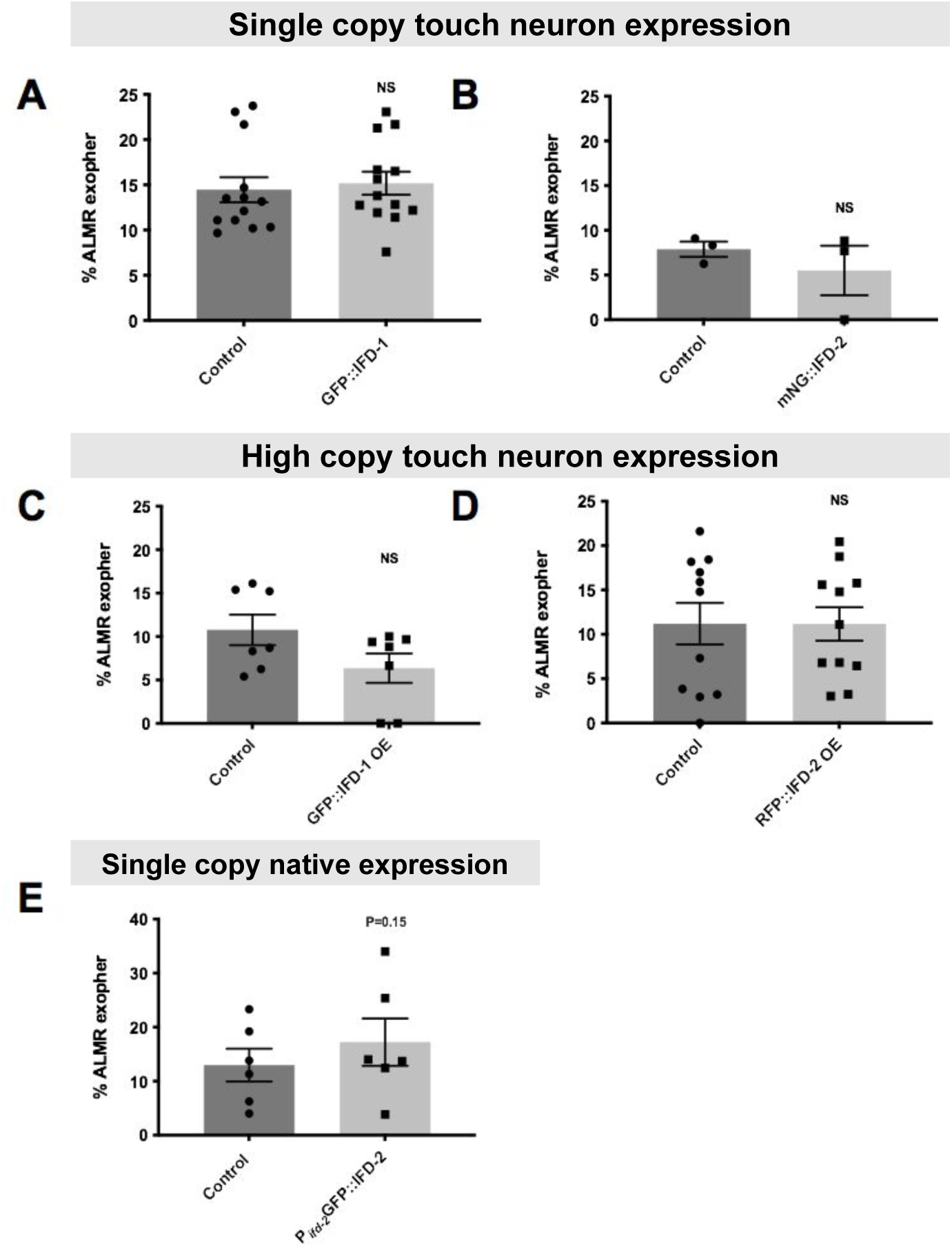
Transgenic IF proteins are neither dominant negative for exopher production nor exopher-elevating on their own. **A)** Single copy *bzIs3*[P*_mec-7_*GFP::IFD-1] does not affect Ad2 ALMR exopher levels. P = 0.92, N > 571/strain, 10 trials, Cochran-Mantel-Haenszel test, error bars are SEM. Strains: control ZB4065 *bzIs166*[P*_mec-4_*mCherry] and ZB4632 *bzIs166*[P*_mec-4_*mCherry];*bzSi3*[P*_mec-7_* GFP::IFD-1]. **B)** Single copy *bzIs37*[P*_mec-7_*mNG::IFD-2] does not affect Ad2 ALMR exopher percentage. P = 0.75, N > 100, 3 trials, Cochran-Mantel-Haenszel test, error bars are SEM. Strains: control ZB4065 *bzIs166*[P*_mec-4_*mCherry] and ZB5091 *bzIs166*[P*_mec-4_*mCherry];*bzSi37*[P*_mec-7_*mNeonGreen::IFD-2]. **C)** Over-expressed (OE) GFP::IFD-1 array does not affect Ad2 ALMR exopher percentage. P = 0.0904, N > 208, 7 trials, Cochran-Mantel-Haenszel test, error bars are SEM. Strains: control ZB4065 *bzIs166*[P*_mec-4_*mCherry] and ZB4501 *bzIs166*[P*_mec-4_*mCherry];*bzEx270*[P*_mec-7_*GFP::IFD-1]. **D)** Over-expressed (OE) RFP::IFD-2 array does not affect Ad2 ALMR exopher percentage. P = 0.987, N > 425, 11 trials, Cochran-Mantel-Haenszel test, error bars are SEM. Strains: control ZB4065 *bzIs166*[P*_mec-4_*mCherry] and ZB4792 *bzIs166*[P*_mec-4_*mCherry];*bzEx253*[P*_mec-7_*RFP::IFD-2]. **E)** An added single copy GFP::IFD-2 transgene expressed from the native *ifd-2* promoter does not change exopher levels. We expressed GFP::IFD-2 from the native *ifd-2* promoter; mCherry background, and scored for exopher levels. We compared ALMR exophers at Ad2 in *bzIs166*[P*_mec-4_*mCherry] to *bzSi76*[P*_ifd-2_*GFP::IFD-2] and found no significant difference, although in two trials, *bzSi76*[P*_ifd-2_*GFP::IFD-2] displayed exopher levels higher than 15%, P = 0.15, 6 trials, N > 300. Our data suggest that the *ifd-*dependent mechanisms influencing exopher levels are unlikely to be strictly dependent on stoichiometric interactions with concentration-limited partner proteins or be anchored in simple enhanced *ifd* expression levels. Lack of over-expression effects are consistent with our observation that size of aggresome compartment per se does not correlate with generally high exopher levels nor does it predict extrusion (Supplementary Figure 10, Supplementary Figure 12).

**Supplementary Figure 11.**
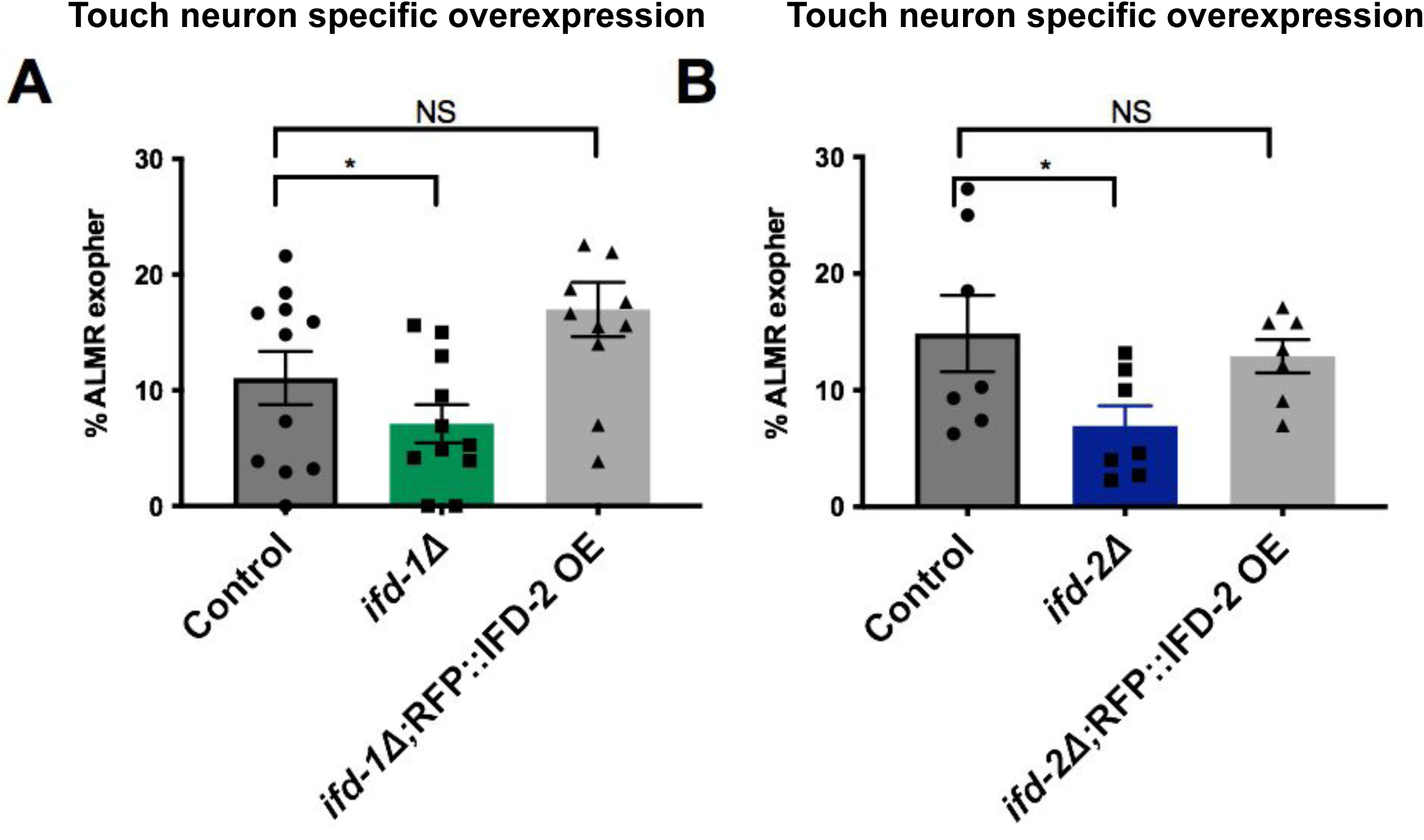
Touch neuron-specific tagged high-copy number IFD transgenes can rescue the IF exopher deficit. **A)** We scored ALMR exophers at Ad2 in *bzIs166*[P*_mec-4_*mCherry] compared to *bzIs166*[P*_mec-4_*mCherry]; *ifd-1(ok2404), (*P < 0.05). *bzIs166*[P*_mec-4_*mCherry] compared to *bzIs166*[P*_mec-4_*mCherry]; *ifd-1(ok2404); bzEx253*[P*_mec-7_*GFP::IFD-2 OE] is NS, P = 0.0642. N > 425, 11 trials. **B)** We scored ALMR exophers at Ad2 in *bzIs166*[P*_mec-4_*mCherry] compared to *bzIs166*[P*_mec-4_*mCherry]; *ifd-2(bz187)* (P < 0.05). *bzIs166*[P*_mec-4_*mCherry] compared to *bzIs166*[P*_mec-4_*mCherry]; *ifd-2(bz187); bzEx253*[P*_mec-7_*RFP::IFD-2 OE] is NS. N > 240, 7 trials. Cross complementation might reflect a need for a particular level for function that cannot be reached from normal gene dosages of *ifd-1* or *ifd-2*.

**Supplementary Figure 12.**
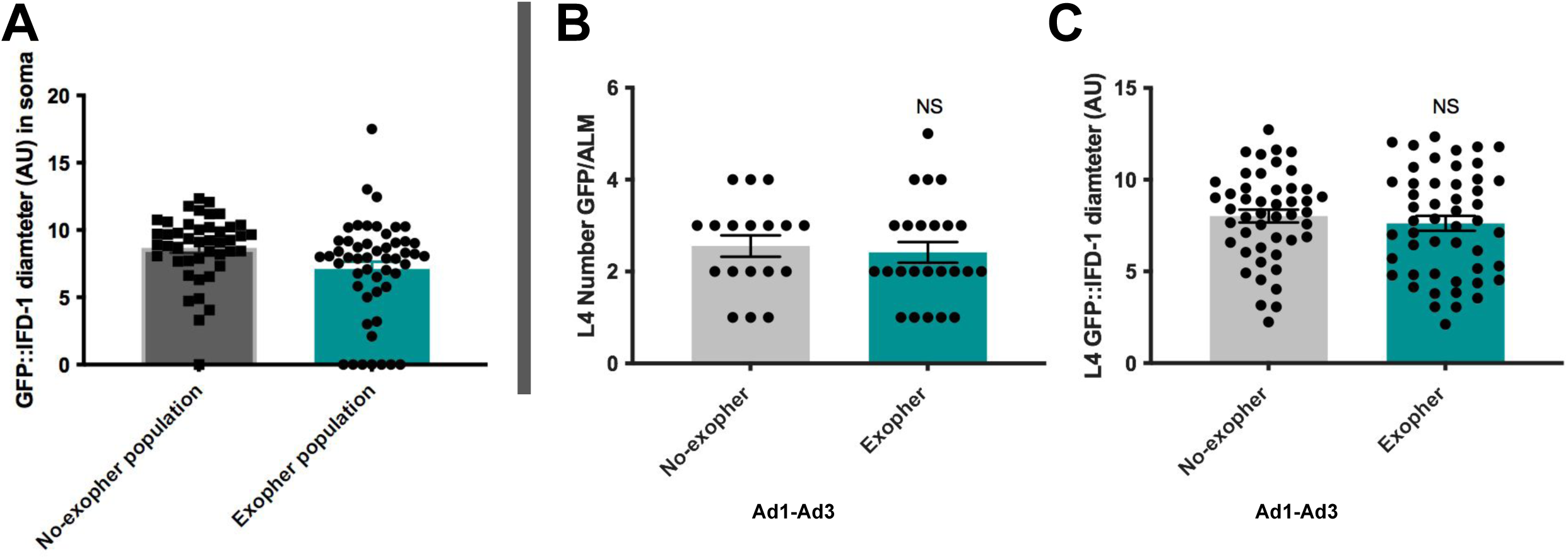
The size of the GFP::IFD-1 concentration does not correlate with exopher rate. **A) GFP::IFD-1-aggresome-like organelles can be cleared from the soma, but the size of aggresome-like organelles that frequently remain in the soma does not correlate with an exopher event.** We imaged 95 Ad3 ALMs from a strain ZB4632 *bzIs166*[P*_mec-4_*mCherry];*bzSi3*[P*_mec-7_*GFP::IFD-1]. 51/95 (54%) neurons displayed an exopher event while 44/95 (46%) did not. We categorized the neurons into two groups: one group that did make an exopher (‘exopher-population’) and a group that did not display an exopher event at time of collection (‘no-exopher population’). We measured the diameter (in AU) of the aggresome-like organelle in the soma in both groups. We found that the ‘exopher+ population’ included several cases in which there was no remaining GFP::IFD-1 aggresome-like organelle in the soma (7/44, or ∼16%) (consistent with the idea that the neuron ejects GFP::IFD-1 aggresome-like organelles discussed in Figure 5B, showing ∼15% GFP::IFD-1-aggresome-like organelle ejection rate). There was one case in which there was no detectable GFP::IFD-1 aggresome-like organelle in the ‘no-exopher’ population. Not considering cases in which there was not a GFP::IFD-1 aggresome-like organelle to measure, we measured no significance in the size of the soma located aggresome-like organelle between the population of animals that made an exopher versus the population that did not make an exopher. P = 0.219 excluding datapoints with 0 aggresome, two-tailed t-test. **B) Early (L4) aggresome-like organelle size does not predict Ad1-Ad3 exophergenesis.** To ask if early aggresome-like organelle size could indicate likelihood of later exophergenesis on adult day 2 or 3, we measured the L4 aggresome-like organelle size and number per cell, and later scored for exopher production. We imaged 216 total L4 neurons and successfully recovered 154/216 animals post-imaging. After 2-3 days on NGM-plates, we scored for exophers (Ad2 and Ad3). Out of 154 animals, 24 had an ALM exopher evident on Ad2/Ad3. To analyze, we binned the animals into two populations: one population that did not make an exopher (‘no-exopher’) and a population that that did make an exopher (‘exopher’). We found no difference in early (L4) aggresome-like organelle number per cell between the two populations. P = 0.67, two-tailed t-test, N successfully scored on Ad2 = 50, 31, 18, 24, and 26 in 5 biologic trials. **C)** Continuing analysis from B, we measured GFP::IFD-1 diameter (AU) in the L4 ALMs in both populations and found NS, P = 0.47 two-tailed t-test.

**Supplementary Figure 13.**
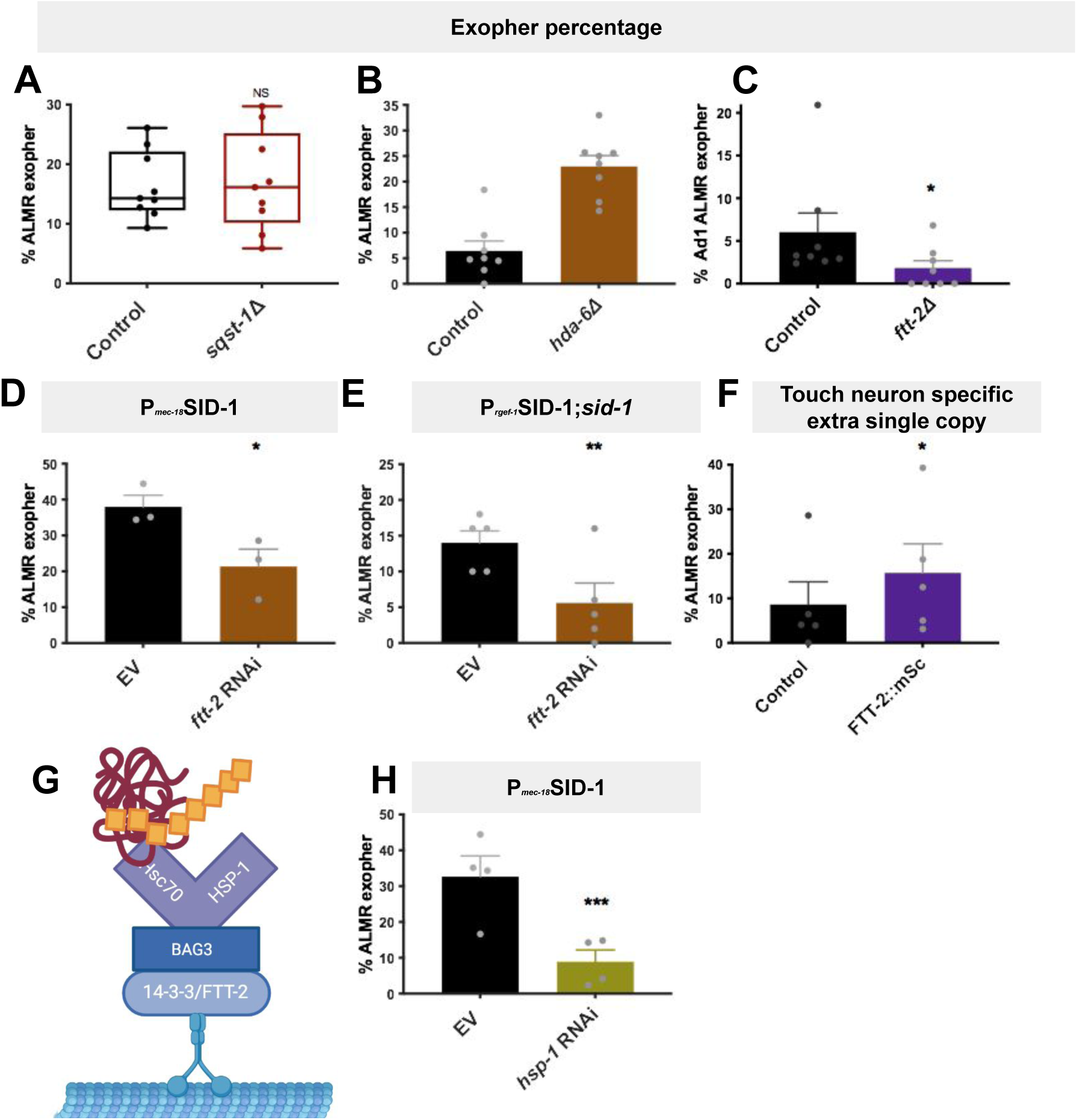
Assessing adaptor protein impact on IFD compartments and exopher production. **Homologs of mammalian aggregate-adapter complex members FTT-2 and HSP-1 have roles in exopher production.** * P < 0.05, ** P < 0.005, *** P < 0.0005. **A) *sqst-1*(*ok2892*) impacts variation in exophergenesis without significantly changing outcome in the *bzIs166*[P*_mec-4_*mCherry] background.** Strains are ZB4065 *bzIs166*[P*_mec-4_*mCherry] and ZB4991 *bzIs166*[P*_mec-4_*mCherry]; *sqst-1*(*ok2892*). We scored ALMR exophers on Ad2 in *bzIs101*[P*_mec-4_*mCherry] compared to *bzIs166*[P_mec-4_mCherry]; *sqst-1(ok2892)IV*. NS, N > 250, 9 trials. Cochran-Mantel-Haenszel, error bars are SEM. **B) *hda-6(ok3203)* increases exophergenesis in the *bzIs166*[P*_mec-4_*mCherry] background on Ad2.** Strains are ZB4801 *bzIs166*[P*_mec-4_*mCherry]; *hda-6(ok3203), bzSi3*[P*_mec-7_*GFP::IFD-1] and ZB4632 *bzIs166*[P*_mec-4_*mCherry]; *bzSi3*[P*_mec-7_*GFP::IFD-1]. P > 0.0005, N > 150, 8 trials. Cochran-Mantel-Haenszel test, error bars are SEM. **C) FTT-2/14-3-3 is required for efficient exopher production.** There is a significant reduction of percentage of ALMR exopher production on Ad1 in the genomic deletion *ftt-2(n4426*) strain expressing *bzIs166*[P*_mec-4_* mCherry]; *bzSi3*[P*_mec-7_* GFP::IFD-1]; P < 0.05, N > 275, 8 trials, Animals measured on Ad1 to avoid animal-bagging. Cochran-Mantel-Haenszel test, error bars are SEM; Ad1. **D) RNAi targeting in all cells + touch neurons supports a 14-3-3/FTT-2 role in efficient exopher production.** We knocked down *ftt-2* in a touch neuron sensitized RNAi strain expressing *bzIs169*[P*_mec-18_*SID-1 P_sng-1_YFP] (other cells in addition to touch neurons can be targeted in this background) from L4 - Ad2. P < 0.05, N > 85, 3 trials, Cochran-Mantel-Haenszel test, error bars are SEM. **E) Pan-neuronal-specific RNAi against *ftt-2* significantly reduces exophergenesis.** We targeted *ftt-2* knockdown using an RNAi clone in a pan-neuronal-specific RNAi strain expressing *sqIs71*[P*_rgef-1_*GFP; P*_rgef-1_*SID-1;pBS]; *sid-1(qt9)*V; *bzIs101*[P*_mec-4_*mCherry] (only neurons subject to *ftt-2* knockdown) to observed significantly reduced Ad2 ALMR exophergenesis. Animals were exposed to the RNAi clone for 2 generations. P = 0.0025, 5 trials, N = 250, Cochran-Mantel-Haenszel test, error bars are SEM. Data are consistent with observations in the *ftt-2(n4426)* putative null mutant in panel C and support FTT-2 function in neurons is required for efficient exopher production. **F) Additional FTT-2 gene dose increases ALMR exophergenesis.** We scored ALMR exopher production on Ad2 in control animals expressing *bzIs166*[P*_mec-4_*mCherry] and *bzIs166*[P*_mec-4_*mCherry]; *bzSi47*[P*_mec-7_*FTT-2::mSc]. Note that the wild type *ftt-2(+)* allele is present and that use of the strong *mec-7* promoter for TN specific expression of the added *ftt-2* copy might elevate expression. Single-copy integrated P*_mec-7_*FTT-2::mSc increases ALMR Ad2 exophergenesis, N > 95, P = 0.035, 3 trials, Ad2. **G) Mammalian aggregate-adapter complex.** Adapted from paper Xu 2013 ^36^ created using BioRender.com. 14-3-3 proteins bind directly to dynein components. 14-3-3 binds BAG3 bound Hsc70/(HSP-1). Hsc70/HSP-1 is able to recognize and bind ubiquitinylated aggregates. The complex moves along the microtubule tracks to deliver to the aggresome compartment. **H) HSP-1 is required for efficient exopher production in a touch neuron-sensitized RNAi strain.** *hsp-1*-targeted RNAi from L4 to Ad2 in a touch neuronal sensitized RNAi strain harboring *bzIs169* [P*_mec-18_sid-1* P_s*ng-1*_YFP]; *bzIs101* [P*_mec-4_*mCherry] (targetting all cells + touch neurons) significantly reduces ALM exophergenesis. Ad2 P < 0.0005, N > 130, 4 trials, Cochran-Mantel-Haenszel test, error bars are SEM. **Discussion.** Note that fertility disruption can be a concern for some knockdowns (^97^ and our unpublished observations). *ftt-2* is expressed in the somatic tissues^98^ and knockdown does not block progeny production; we scored on Ad1 to avoid bagging and retention of progeny that can occur at Ad2. *hsp-1* is a more broadly expressed essential gene that can disrupt fertility; so RNAi knockdown, though anticipated to be enhanced in touch neurons ^55^ and not blocking all progeny production in our experiment, might still impair exophers by compromising fertility or other tissue health. We tested but did not find evidence of a requirement for an *unc-23*/BAG3 mutation or RNAi on exophers (data not shown). BAG3 and UNC-23 are only partially overlapping and we speculate that the HSP-1/Hsc70/FTT-2/14-3-3 complex might be simpler in nematodes as compared to mammals, or another protein may provide the UNC-23 role in the complex. To more firmly establish cell autonomy we constructed degron tagged alleles of *ftt-2* and *hsp-1* for later auxin-induced degradation in touch neurons. We found that *degron::hsp-1* was homozygous lethal; and *degron::ftt-2* was constitutively lowered for exopher production, even in the absence of auxin, suggesting the tagged variant alone reflects the null phenotype. Although we were unable to use AID for touch neuron specific degradation, the *degron::ftt-2* tag serves as an independent allele to confirm the exopher suppression phenotype.

**Supplementary Figure 14.**
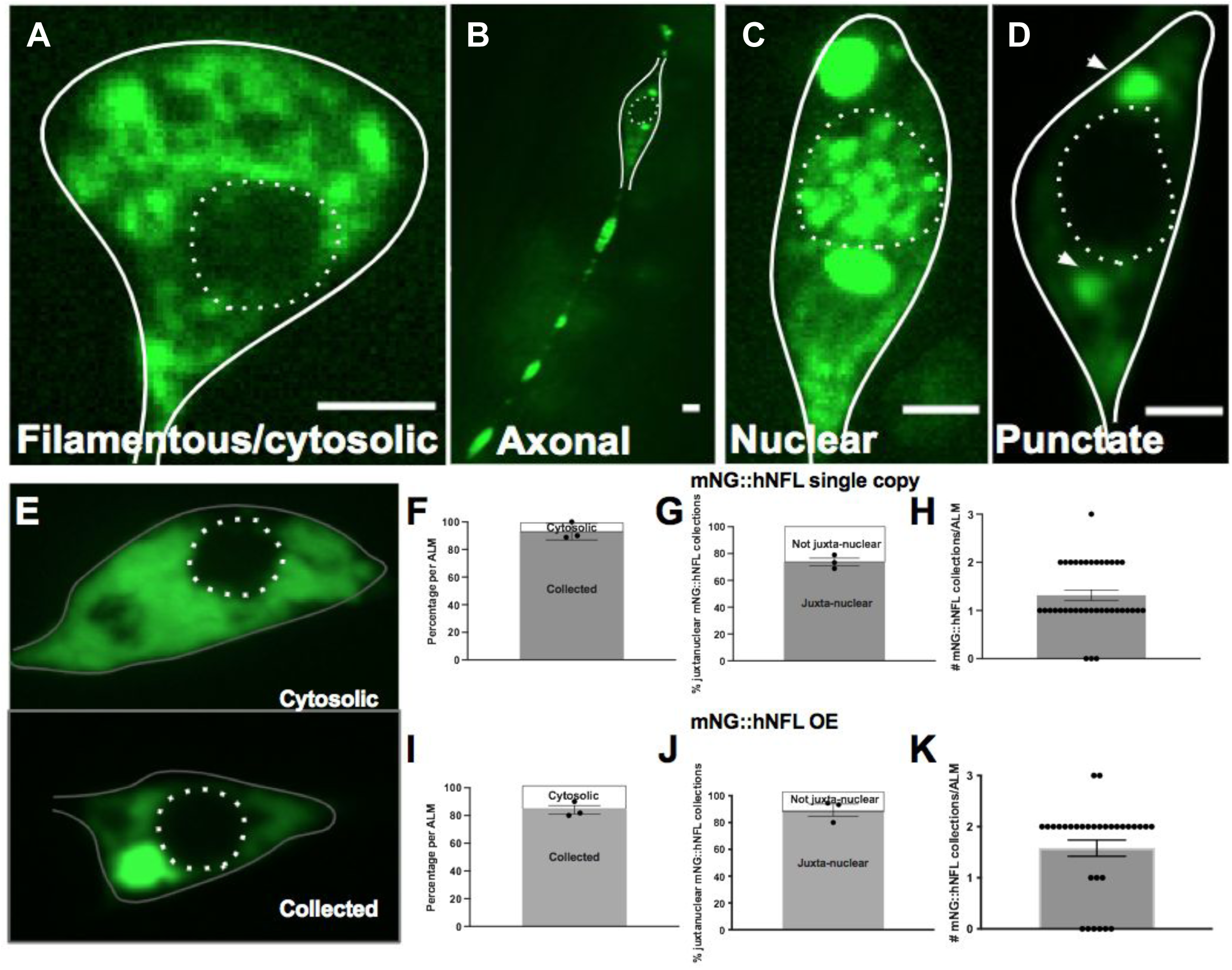
Human intermediate filament Neurofilament Light Chain (hNFL) can adopt multiple localization patterns in *C. elegans* touch neurons, but prominently localizes to 1 or 2 juxtanuclear inclusions. **A-D) Human hNFL can adopt multiple localization patterns when expressed in the *C. elegans* touch neurons – A)** filamentous/cytoplasmic (Strain: ZB5345 *bzEx311*[P*_mec-7_*mNG::hNFL];*bzIs166*[P*_mec-4_*mCherry]; *ifd-2(bz187)*). **B)** axonal (and punctate featured). **C)** nuclear (and punctate featured). And **D)** most commonly, juxtanuclear and punctate. Scale bar = 2 µm. **B-D)** Strain: ZB5175 *bzEx269*[P*_mec-7_*mNG::hNFL];*bzSi34*[P*_mec-7_*mSc::IFD-1]. **E-K) Human mNG::hNFL forms juxtanuclear inclusions in either single copy or high copy array lines. E)** Top - filamentous/ cytosolic localization. Bottom – typical punctate/collected localization. Strain: ZB5593 *bzIs166*[P*_mec-4_*mCherry];*bzEx311*[P*_mec-7_*mNG::hNFL]. Top **(F, G, H**) quantification of integrated single copy strain ZB5332 *bzSi48*[P*_mec-7_* mNG::hNFL]; *bzIs166*[P*_mec-4_*mCherry]. Bottom **(I, J, K)** quantification of high copy array quantification of strain ZB5593 *bzIs166*[P*_mec-4_*mCherry];*bzEx311*[P*_mec-7_*mNG::hNFL]. Collected or cytosolic localization phenotype quantified in **F** and **I**. Juxtanuclear location quantified in **G** and **J**. Number of collected punctate quantified in **H** and **K**. Scale bar = 2 *µm*. Representative of N > 10.

**Supplementary Figure 15.**
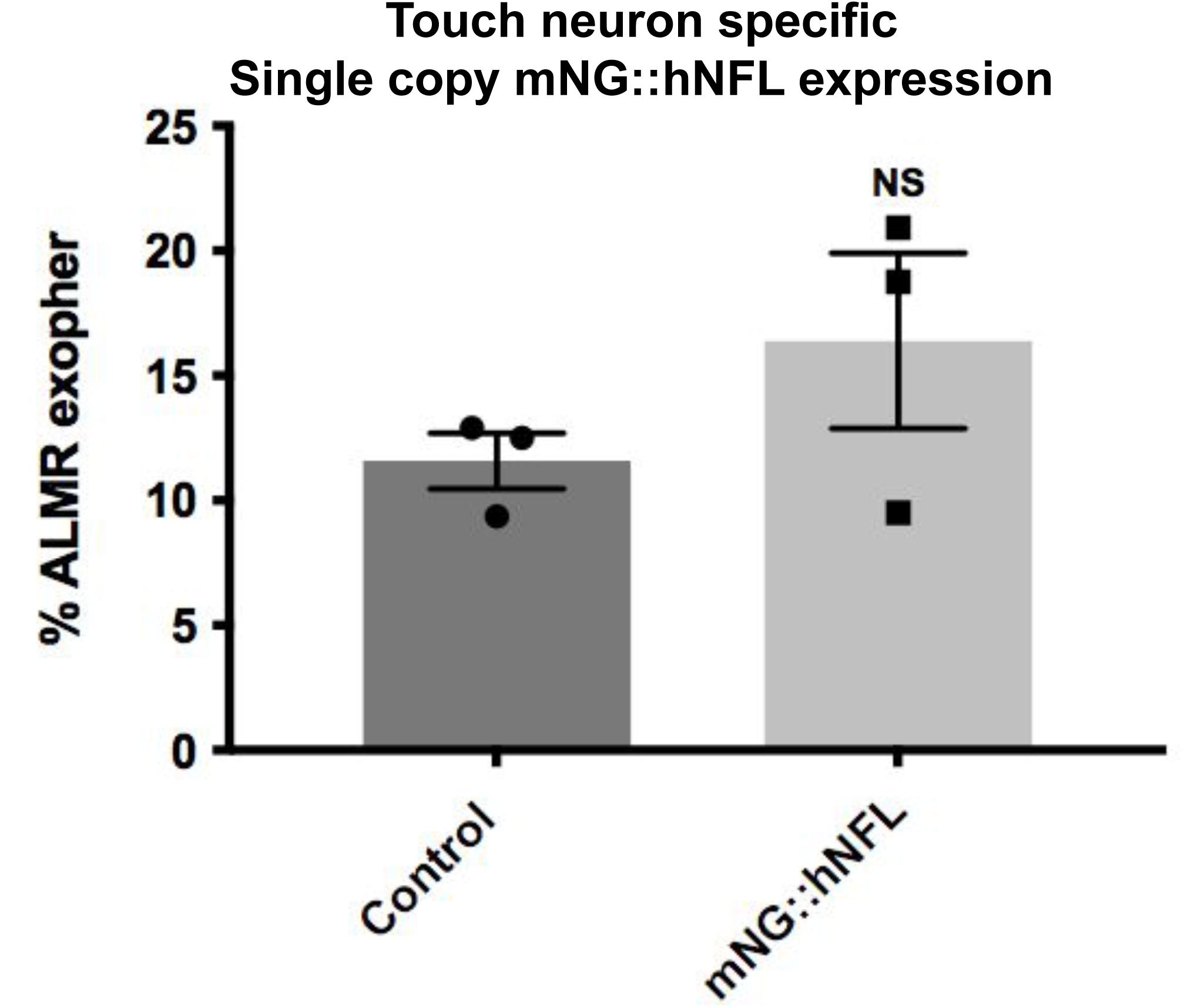
Single copy transgenic hNFL tagged protein is not dominant negative for exopher production nor significantly exopher-elevating on its own. Single copy mNG::hNFL does not significantly affect Ad2 ALMR exopher percentage. P = 0.442, N > 94, 3 trials, Cochran-Mantel-Haenszel test, error bars are SEM. Strains: control ZB4065 *bzIs166*[P*_mec-4_*mCherry] and ZB5332 *bzIs166*[P*_mec-4_*mCherry];*bzSi48*[P*_mec-7_*mNG::hNFL].

**Supplementary Figure 16.**
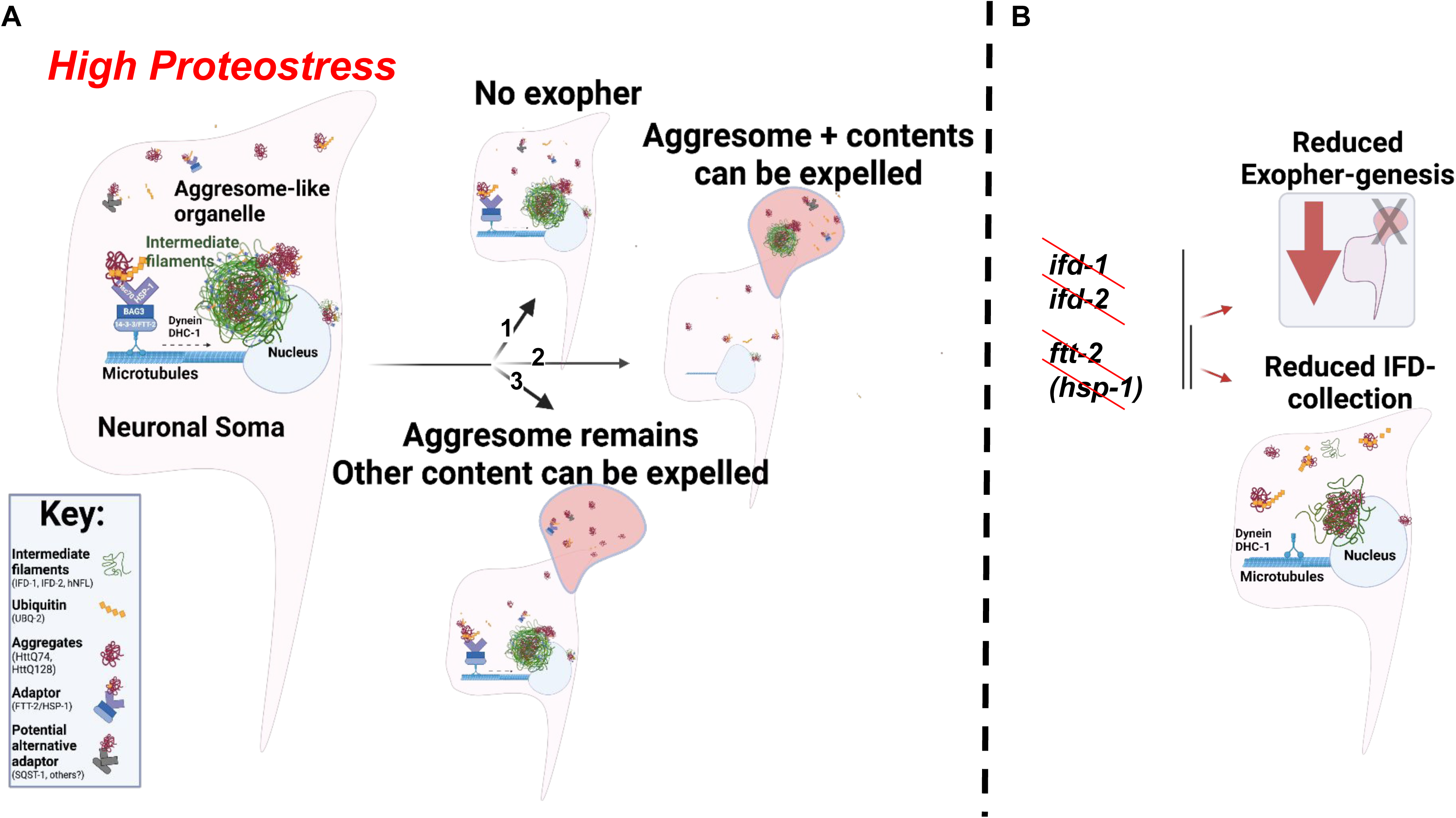
Summary model of *C. elegans* aggresome-like organelle formation, ejection, and the requirements of efficient exopher-genesis. **A)** *C. elegans* mechanosensory touch neurons typically contain 1-3 aggresome-like intermediate filament-decorated collections that can contain ubiquitin (UBQ-2) and aggregated proteins (HttQ74 and HttQ128). Adaptor proteins (FTT-2(14-3-3), from the 14-3-3/BAG3/Hsc70 complex) shuttle ubiquitin tagged aggregated proteins facilitated by dynein transport on the microtubule track for juxtanuclear collection. Furthermore FTT-2(14-3-3) and HSP-1(Hsc70) (as well as ubiquitin and HttPolyQ aggregates, and SQST-1 a potential alternative aggresome-adaptor) can colocalize with the IF-aggresome-like site. One large IF-aggresome-like organelle is depicted, to emphasize detail, and a smaller IF-aggresome-like collection is shown on the other side of the nucleus. Note that the IF-aggresome-like organelle can contain aggregate compactions that are not encased or fully colocalized with IF-protein. In situations of high proteostress, *C. elegans* touch neurons can facilitate aggregate ejection, exophergenesis (outcome 2 and 3). The IF-organelle and its contents can be ejected in the exopher (∼15-50% of exopher events, Figure 5) (option 2), or can remain in the soma as other contents can be expelled (option 3). **B)** Without intermediate filament proteins IFD-1 and IFD-2 there is reduced exophergenesis. IFD-1 and IFD-2 act cell autonomously, within the neuron, in the exopher-pathway. Aggresome adaptor proteins, FTT-2(14-3-3), and HSP-1(Hsc70) of the 14-3-3/BAG3/Hsc70 adaptor complex are also required for efficient exophergenesis, linking proteins in aggresome biology with cellular extrusion functions. FTT-2, which is hypothesized to work with HSP-1 in the IF-aggresome formation pathway, is required for efficient IFD-collection at the aggresome site. With *ftt-2* knockout or knockdown, there is reduced IFD-collection at the juxta nuclear aggresome-like site.

**Supplementary Figure 17.**
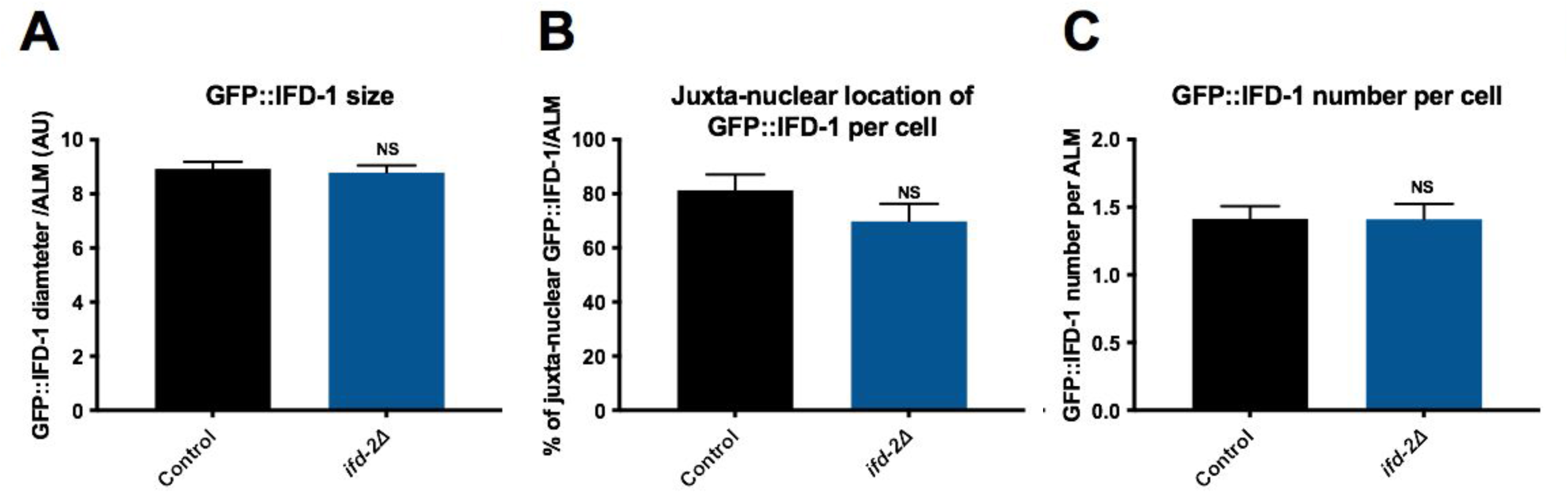
IFD-1-puncta form without *ifd-2*. **A) IFD-1-positive puncta size remains the same in *ifd-2(bz187)*.** Diameter, in µm, of Ad2 IFD-1-positive puncta in ALM neurons remains similar to WT in the *ifd-2(bz187)* background in a strain expressing *bzIs166*[P*_mec-4_* mCherry]; *bzSi3*[P*_mec-7_* GFP::IFD-1]. P = NS, N > 55, 3 trials, two-tailed t-test, error bars are SEM. **B) IFD-1-positive puncta form juxtanuclearly with or without *ifd-2(bz187).*** Percentage of juxtanuclear IFD-1-positve puncta per ALM does not significantly differ in strains expressing *bzIs166*[P*_mec-4_* mCherry]; *bzSi3*[P*_mec-7_* GFP::IFD-1], with or without *ifd-2(bz187)*. P = NS, N > 37, 3 trials, two-tailed t-test, error bars are SEM. Juxtanuclear location assessed in a binary manner. **C) IFD-1-positive puncta number per ALM neuron remains the same in control and *ifd-2(bz187).*** Number of IFD-1-positive inclusions in Ad2 ALM somas does not change in mutant *ifd-2(bz187)* versus WT in strains expressing *bzIs166*[P*_mec-4_* mCherry]; *bzSi3*[P*_mec-7_* GFP::IFD-1]. NS, N > 39, 3 trials, two-tailed t-test, error bars are SEM.

**Supplementary Table 1: List of strains and RNAi details**.

Video link 1: https://youtu.be/zMvtUJJbrzc

Video link 2: https://youtu.be/Gtex9gShdR8

